# Identification of cytoplasmic incompatibility effectors of the reproductive manipulator *Cardinium hertigii*

**DOI:** 10.64898/2026.06.05.730497

**Authors:** Olivia L. Mathieson, Dylan L. Schultz, Matthew R. Doremus, Simina Vintila, Suzanne E. Kelly, Martha S. Hunter, Stephan Schmitz-Esser, Manuel Kleiner

## Abstract

Cytoplasmic incompatibility (CI), a widespread symbiont-induced reproductive manipulation phenotype in arthropods, selectively kills embryonic offspring produced when infected males mate with uninfected females. Most knowledge on CI comes from studying the Pseudomonadota symbiont *Wolbachia* and their CI effector (*cif*) genes, and little is known about CI mechanisms of symbionts without *cif* genes. We identify two putative CI effectors of *Cardinium hertigii c*Eper1, a Bacteroidota symbiont of the parasitoid wasp *Encarsia suzannae*, using a combined proteomics and heterologous expression approach. We show that these genes, CAHE_0406 and CAHE_p0043, have no homology to known *cif* genes, yet impair yeast growth similar to known *cif*s and that CAHE_p0043 localizes to diffuse and fragmented yeast nuclei. These results suggest that CAHE_0406 and/or CAHE_p0043 could play a role in *c*Eper1-induced CI, expanding our current understanding of the diversity of proteins involved in cytoplasmic incompatibility across symbiont lineages.

## Introduction

Heritable bacterial endosymbionts are common in arthropods. These symbionts are often transmitted from mother to offspring by transovarial transmission (symbionts packaged inside the maternal gamete) (Doremus and Hunter, 2020). This reliance on female host reproduction is frequently coupled with modification of host reproduction by the symbiont to favor infected females at the expense of male hosts and/or uninfected individuals. A common symbiont-induced reproductive manipulation is cytoplasmic incompatibility (CI), which promotes the fitness of symbiont-carrying and -transmitting females by causing death of embryonic offspring produced from a symbiont-carrying male and a symbiont-free female (Doremus and Hunter, 2020). Symbiont-induced CI defects associated with sperm will lead to embryonic death unless reversed or prevented in symbiont-carrying zygotes. Symbionts from several bacterial phyla are known to induce CI (Owashi et al., 2024; Pollmann et al., 2022; Rosenwald et al., 2020; Takano et al., 2021, 2017; Umanzor et al., 2025). However, nearly all reported insights into the mechanisms underlying CI come from *Wolbachia* (phylum: Pseudomonadota, class: Alphaproteobacteria) (Duron et al., 2008; Shropshire et al., 2020; Werren et al., 2008), and little is known about CI mechanisms of other symbionts.

*Candidatus* Cardinium hertigii (phylum: Bacteroidota), hereafter *Cardinium*, is a widespread endosymbiont of invertebrates that induces CI in a range of arthropod hosts, including parasitoid wasps in the genus *Encarsia* (Gebiola et al., 2016; Hunter et al., 2003; Mathieson et al., 2025). The *c*Eper1 strain hosted by the wasp *Encarsia suzannae* was the first CI *Cardinium* strain identified (Hunter et al., 2003). Initial analyses of the *c*Eper1 genome revealed no instances of horizontally transferred genes implicated in host cell interaction between *Wolbachia* and *Cardinium* (Penz et al., 2012), and later comparisons to *Wolbachia* genomes found that the *c*Eper1 genome does not encode homologs of *Wolbachia* CI effectors (Beckmann et al., 2017; LePage et al., 2017; Mann et al., 2017). Recently, homologs to *Wolbachia* CI effector (*cif*) genes have been identified in two genomes of *Cardinium* strains not associated with reproductive manipulation (Amoros et al., 2025). Even though CI effector proteins are not shared between CI-inducing *Wolbachia* and *Cardinium* strains, both symbionts cause offspring mortality through similar mitotic defects that result in aberrant chromosomal condensation and chromatin bridging, suggestive of convergent evolution of CI in these distantly related symbionts (Gebiola et al., 2017).

Symbiont localization patterns during *E. suzannae* spermatogenesis provide some insights into the timing of *c*Eper1 CI induction. During male *E. suzannae* pupation, *c*Eper1 infects most developing sperm cysts but is removed along with sperm cytoplasmic contents during spermiogenesis, the nuclear and cellular elongation stage of sperm development that culminates in mature sperm cells (Doremus et al., 2020; Ferree et al., 2019). In *E. suzannae*, the timing of *Cardinium* removal from sperm coincides with the end of host pupation, which suggests the male pupal stage is critical for CI induction (Doremus et al., 2020). These observations, together with mechanisms proposed for *Wolbachia* CI induction (Beckmann et al., 2019; Shropshire and Bordenstein, 2019), suggest two alternate hypotheses for the alteration of male function by *Cardinium*: pre-fertilization modification or post-fertilization toxicity (Doremus et al., 2020). The pre-fertilization modification hypothesis proposes that the reproductive manipulator produces a product that modifies host chromatin or other sperm-associated factors in a way that causes early death in embryogenesis (Doremus et al., 2020). In the post-fertilization toxicity hypothesis, a toxin is produced by the reproductive manipulator during spermatogenesis, remains in the sperm cells after the symbiont cells are removed and becomes toxic when the chromatin decondenses after fertilization (Doremus et al., 2020). In both scenarios for the effect of *Cardinium* on males, however, we assume that the CI phenotype is rescued when sperm from a *Cardinium*-carrying male arrives in an egg of a *Cardinium*-carrying female, allowing mitosis to proceed successfully. While these are the most plausible hypotheses given current evidence, it is possible that *c*Eper1 CI is driven by mechanisms entirely different from *Wolbachia* CI.

*Wolbachia* CI effectors (*cif* genes) are typically found as gene pairs encoded in close proximity in the genome, with one or both required for induction and one as a rescue factor, and both encoding various eukaryotic interaction domains (Amoros et al., 2025; Beckmann et al., 2019; Chen et al., 2019; LePage et al., 2017; Shropshire and Bordenstein, 2019; Wang et al., 2022). In the absence of evidence to the contrary, we might expect *Cardinium* CI factors to be similar, for example consisting of two adjacent genes containing eukaryotic interacting domains. A previous transcriptomic study of adult *Encarsia* has identified genes transcribed by *c*Eper1 as candidates for CI involvement that share these characteristics (e.g., paired genomic localization, eukaryotic interacting proteins) (Mann et al., 2017). These include ankyrin repeat domain-containing proteins, outer membrane protein beta-barrel domain-containing proteins, a DEAD box RNA helicase, and several uncharacterized proteins previously identified through genomics (Penz et al., 2012), some of which are shared across other *Encarsia*-associated *Cardinium* strains (Schultz et al., 2025).

Functional analyses are necessary to narrow the list of potential *Cardinium* CI effectors. While the transcriptome of *Cardinium* in adult *Encarsia* suggested some candidate effectors for CI (Mann et al., 2017), the localization studies indicated that CI induction in *E. suzannae* by the *c*Eper1 strain occurs primarily at the pupal, not the adult, stage (Doremus et al., 2020). Further, studies to confirm translation and assess the function of candidate proteins have not yet been conducted in *Cardinium*. A gel-based proteomics approach was critical for identifying *Wolbachia* effector proteins in *Culex pipiens* (Beckmann and Fallon, 2013), but the *Cardinium* host *E. suzannae* is roughly 1/100th of the size of a *C. pipiens* mosquito and thus a different, more sensitive proteomics approach was needed. Lastly, heterologous expression of CI effector candidates in *Saccharomyces cerevisiae* proved an effective screening strategy for induction and rescue potential of *Wolbachia* CI effectors (Beckmann et al., 2017), providing a promising approach for screening *Cardinium* CI effector candidates.

Here, we investigated the CI-inducing *Cardinium c*Eper1 strain in the parasitoid wasp *E. suzannae* to identify and assess effectors associated with *Cardinium*-induced CI. We identified putative CI effectors by combining previous cytological and genomic insights with highly sensitive LC-MS/MS-based differential proteomics, measuring host tissue-, sex-, and life stage-specific proteomes of the *c*Eper1 strain. We then heterologously expressed putative CI effectors and homologs from other *Encarsia*-associated *Cardinium* strains in yeast to assess their impact on cell growth and localization within yeast cells. Based on these results, we propose two putative *Cardinium* CI effectors that reduce yeast growth, and highlight several other potential effectors based on protein abundance patterns.

## Results

### *c*Eper1 proteome across host tissues, sexes, and life stages

We generated two proteomics datasets to compare *Cardinium* protein abundances across pupal and adult male and female hosts as well as between reproductive organs and heads (**Figure 1A**). The first dataset included three different fractions generated by differential pelleting of whole wasp homogenates. The three fractions analyzed were an initial cell debris/heavy pellet (P), a *Cardinium* enrichment pellet (PS), and the supernatant (S). The second dataset was from complete lysates of specific dissected tissues (ovaries, testes, heads).

**FIGURE 1.**
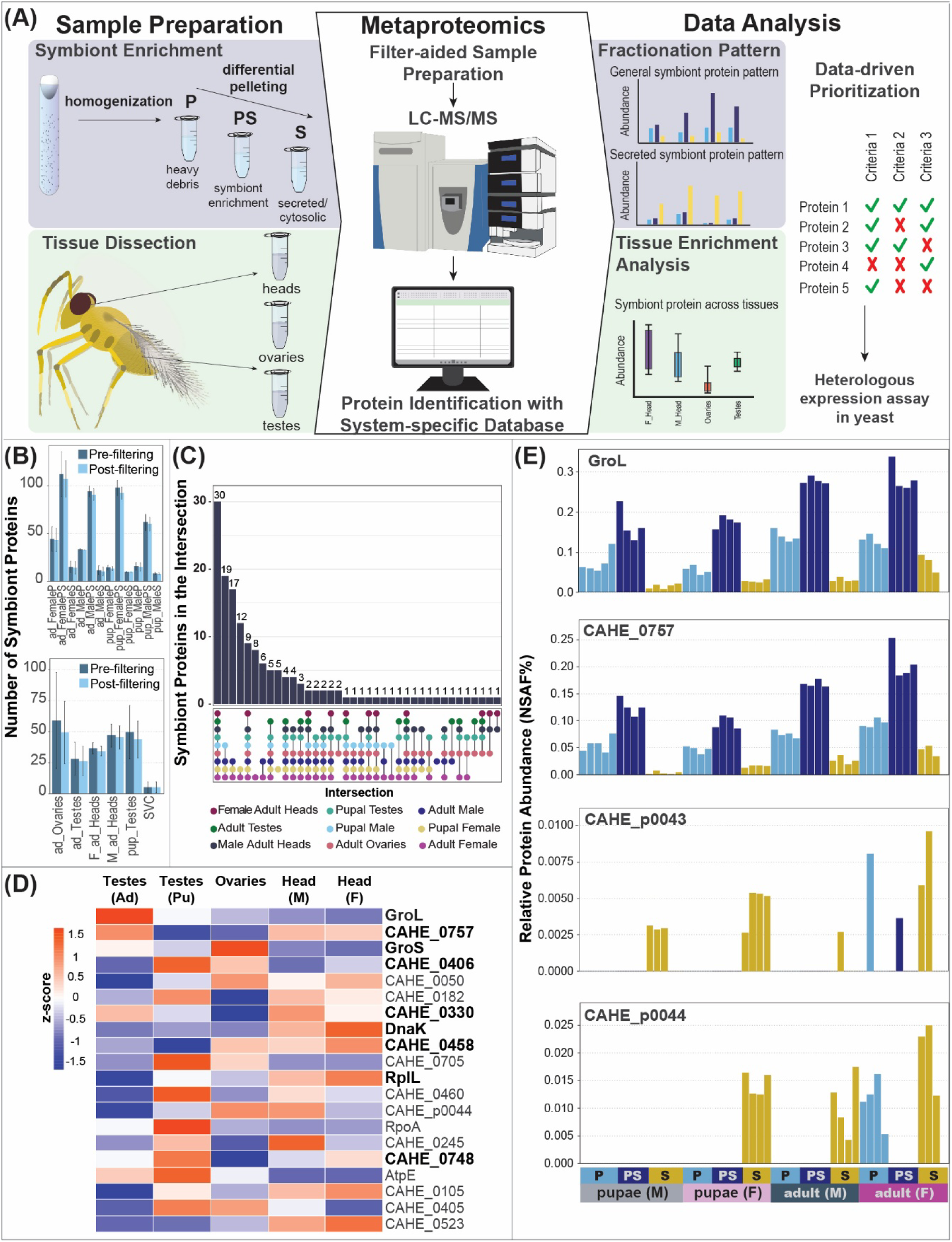
Identification of CI effector candidates in the *c*Eper1 proteome. **(A)** Overview of methods and analysis approaches for identifying proteinaceous effectors using proteomics. **(B)** Mean number of *Cardinium* proteins detected per sample type at a false discovery rate of 5% and classified as a master protein before and after filtering for detection in at least 50% of at least one sample type. Error bars indicate standard deviations (n =4, except for S and P fractions of pupal males where n = 5). SVC = adult seminal vesicle contents, ad = adult, pup = pupae, F = female, M = male, and P, PS, and S = sample fraction. **(C)** Overlap of *Cardinium* protein detection across sample types in the filtered datasets. The y-axis indicates the number of proteins shared by the sample intersections (x-axis). The colors of the points in the intersection matrix indicate sample type. **(D)** Comparison of abundances of the top 20 *Cardinium* proteins in pupal testis proteomes across five tissue types: adult (Ad) testes, pupal (Pu) testes, ovaries, male (M) head, and female (F) head. Protein abundances (average orgNSAF%) were centered across the row using a z-score transformation, with dark blue indicating lower abundance and red indicating higher abundance. Proteins are listed from most abundant in pupal testes (top) to least abundant (bottom) and bolding indicates that the protein was detected in the seminal vesicle proteome. **(E)** Detection pattern of *Cardinium* proteins (CAHE_p0043, CAHE_p0044, CAHE_0757, and GroL) across fractions obtained from whole wasp homogenates by differential centrifugation. The x-axis shows individual samples grouped by fraction, host sex, and life stage. Yellow = symbiont secretome fraction (S), dark blue = symbiont cell enrichment fraction (PS), and light blue = host cell debris fraction (P).

We detected 11,285 *E. suzannae* proteins and 210 *c*Eper1 proteins in our fractionated sample dataset, with *Cardinium* making up an average of 0.75% (± 0.77 SD) of the proteinaceous biomass in the samples (**Supplementary Data 1**). *Cardinium* proteins were most abundant in the symbiont enrichment fraction (PS) with 143 *Cardinium* proteins detected in at least two replicates of one sample type (sex, life stage). Due to the higher coverage of the *Cardinium* proteome in the PS fraction compared to the P and S fractions, we are referring to the PS fraction when comparing symbiont protein abundances between host sexes and life stages unless otherwise stated.

We detected fewer *Cardinium* proteins in pupae (**Figure 1B**; Male = 59.75 mean ± 7.09 SD; Female = 92.25 mean ± 6.50 SD) compared to adults (**Figure 1B**; Male = 90.50 mean ± 6.24 SD; Female = 106.8 ± 19.1 SD), and overall, detected fewer *Cardinium* proteins in males compared to females in both life stages. In contrast, we detected more host proteins in the pupal life stage compared to the adult life stage (**Figure S1**). In the male and female reproductive and head tissue proteomes, we detected a total of 8,052 *E. suzannae* and 149 *c*Eper1 proteins. The tissue-specific samples had a higher *Cardinium* proteinaceous biomass than the fractionated whole wasp with an average of 2.02% (± 1.80 SD) with the highest biomass contribution being in the pupal testes (4.72% mean ± 2.07% SD) (**Supplementary Data 1**), suggesting enrichment of *Cardinium* in pupal testes relative to the rest of the body. We found that 30 *Cardinium* proteins were consistently detected across all sample types (**Figure 1C**). Generally, these proteins were central to symbiont physiology (e.g., GroL, EF-Tu, DnaK, ribosomal proteins, and ATPase subunits); however, they also included proteins of unknown function, such as CAHE_0757, a predicted lipoprotein, which was among the most abundant *Cardinium* proteins consistently detected. Another abundant *Cardinium* protein was the tail sheath protein (CAHE_0458) of the type VI secretion system (T6SS), which we consistently detected in higher abundance in male versus female hosts (q = 3.32e-04, Welch’s T-test with Benjamini-Hochberg FDR correction), regardless of life stage (q = 0.01, both adult and pupae, Welch’s T-test with Benjamini-Hochberg FDR correction, **Figure S2**). However, other components of the T6SS had differing detection patterns, including CAHE_0459 which was enriched in females (q = 1.46e-06, sex only; q = 0.064, adult; q = 1.30e-04, pupae; Welch’s T-test with Benjamini-Hochberg FDR correction, **Figure S2**). We also detected 123 proteins in unique intersects of sample types (**Figure 1C**), suggesting that our proteomics approach captured several snapshots of *Cardinium* physiology. We leveraged the specific detection patterns of *Cardinium* proteins to identify proteins potentially involved in CI by (1) identifying proteins enriched in CI-relevant host sex, life stage, and tissue-types and (2) identifying potentially secreted *Cardinium* proteins.

### *Cardinium* proteins with putative CI involvement based on differential abundance patterns across tissues, life stage and sex

Prior work highlighted the pupal testes as the focal life stage and tissue for CI induction (Doremus et al., 2020). The 20 most abundant *Cardinium* proteins in pupal testes included several proteins that were detected across nearly all sample types (GroL, GroS, DnaK, EF-Tu, RpoA, RplL, PepA (CAHE_0748), CAHE_0458 and CAHE_0757; **Figure 1D**). There were also several T6SS proteins (CAHE_0458, CAHE_0460), an ATP synthase subunit (AtpE), ABC transporter substrate-binding protein (CAHE_0245), and uncharacterized proteins (CAHE_0406, CAHE_0405, CAHE_0523, CAHE_0105, CAHE_p0044, CAHE_0705, CAHE_0182, CAHE_0050) (**Figure S3**). Nine of the 20 most abundant pupal testes proteins were also detected in adult seminal vesicle samples (**Figure 1D**). These included CAHE_0757 and CAHE_0406, both uncharacterized predicted lipoproteins. We detected CAHE_0757 consistently across all tissue types (**Figure S1**). CAHE_0406 was much less abundant than CAHE_0757 and was only detected in one out of four seminal vesicle samples. We observed a trend of higher abundance of CAHE_0406 in pupal testes relative to other tissues (**Figure 1D, S1**). CAHE_0405, another protein detected in the top 20 proteins in the pupal testis, was adjacent to CAHE_0406 in the genome (**Figure S4**). We observed a similar detection pattern for CAHE_0405 to that of CAHE_0406, only detecting the protein in pupal testes, adult ovaries, and adult male heads (**Figure 1D, S1**). In the fractionated whole wasps, we observed increased detection of CAHE_0405 in pupal males relative to pupal females (q = 0.094, Welch’s T-test with Benjamini-Hochberg FDR correction; **Figure S1**) as well as adult males (q = 0.11, Welch’s T-test with Benjamini-Hochberg FDR correction; **Figure S1**). CAHE_0406 was detected consistently at a higher abundance than CAHE_0405 in the symbiont fractions (PS) enriched from whole wasps (**Figure S1**). The shared pattern of enrichment of CAHE_0406 and CAHE_0405 in the pupal testis and the adult ovary proteomes combined with adjacent positioning in the genome suggest possible involvement in CI. In addition, we included CAHE_0757 as a candidate due to its high abundance across all tissues and consistent detection in the seminal vesicle samples (3 out of 4 replicates).

### Identification of putatively secreted *Cardinium* proteins as CI effector candidates

We used the proteomes from fractionated whole wasps to identify putative secreted effectors by comparing the detection pattern of proteins in the *Cardinium* proteome across the three fractions generated by differential pelleting. Each fraction had a distinct symbiont proteome composition (e.g., overall number of proteins, identity of proteins), making standard statistical testing challenging. Therefore, we relied on manual inspection of protein abundance to group proteins according to detection patterns. Generally, *Cardinium* proteins were most abundant in the symbiont enrichment fraction (PS), with fewer symbiont proteins being detected in the cell debris fraction (P) and even fewer to no proteins detected in the supernatant fraction (S) (**Figure 1E**; **Supplementary Data 2**). CAHE_p0043 and CAHE_p0044, two adjacent plasmid-encoded uncharacterized proteins (**Figure S5)**, were exceptions to this pattern, as they were detected almost exclusively in the S fraction and were mostly absent in the PS fraction (**Figure 1E**). This contrasts with CAHE_0757, which was detected in all fractions and most enriched in the PS fraction, similar to the pattern of GroL detection (**Figure 1E**). This fractionation pattern indicates that CAHE_p0044 and CAHE_p0043 are likely secreted and potentially interact directly with the host. We also detected both CAHE_p0044 and CAHE_p0043 in adult testes and ovaries, and in pupal testes (**Figure S1**), with CAHE_p0044 being in the 20 most abundant symbiont proteins detected in the pupal testis samples (**Figure 1D**). However, neither of these proteins were detected in the seminal vesicles. We considered CAHE_p0043 and CAHE_p0044 CI effector candidates due to the genomic co-localization of these genes combined with the detection pattern suggesting secretion. CAHE_0406 and CAHE_0405 did not exhibit the secretion pattern that was observed for CAHE_p0043 and CAHE_p0044 (**Figure S6**).

### Heterologous expression of CI effector candidates impacts yeast growth

We screened the five *c*Eper1 CI candidate genes identified by proteomics (CAHE_0406, CAHE_0405, CAHE_p0044, CAHE_p0043, CAHE_0757) for toxicity using yeast heterologous expression assays (Sun et al., 2022; Xiao et al., 2021) (**Figure 2A,C**). We confirmed induction of the heterologous gene using proteomics on induced and uninduced batch cultures (**Figure 2B,D**).

**FIGURE 2.**
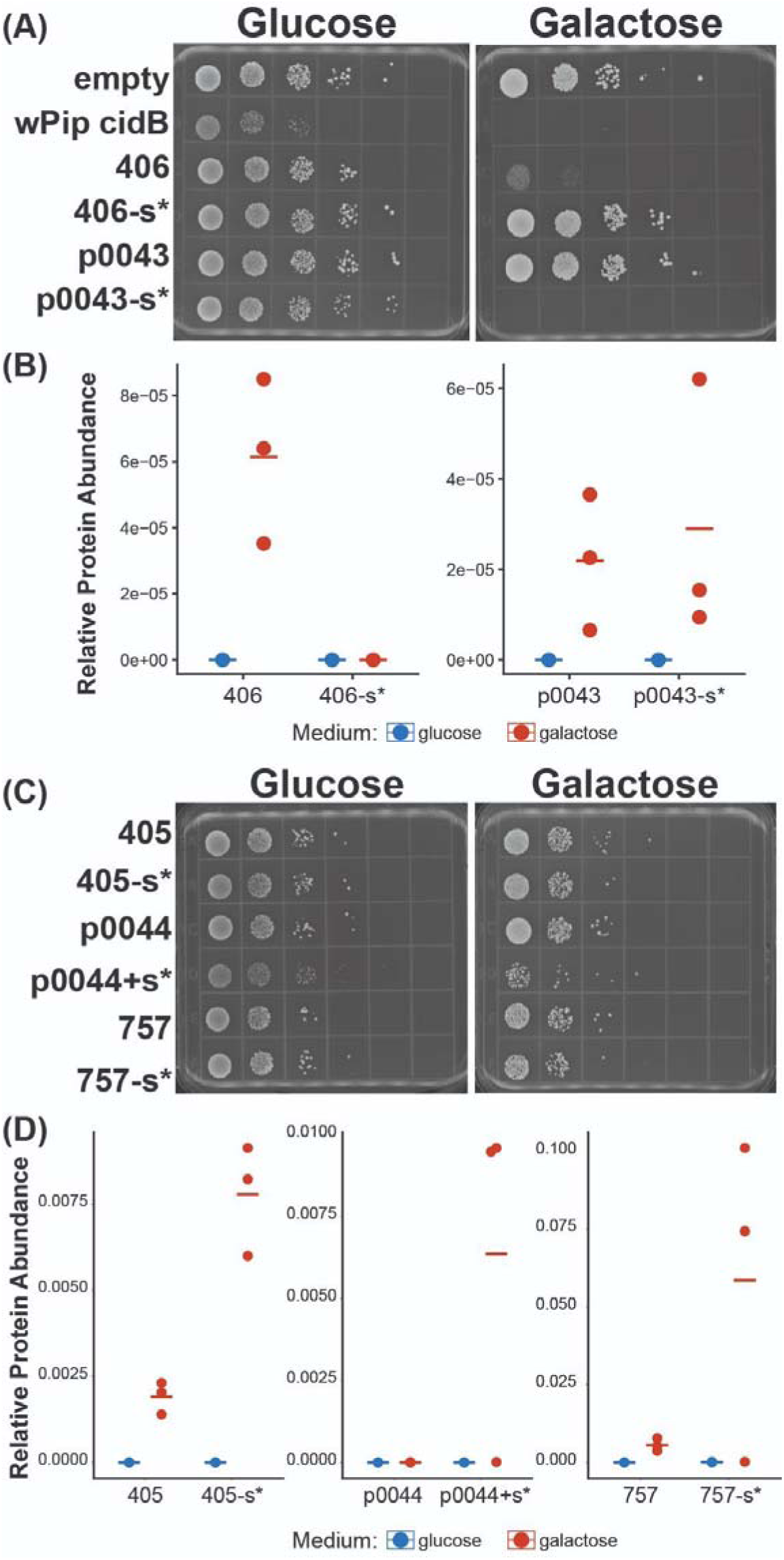
Heterologous expression of *Cardinium c*Eper1 CI candidates in yeast. **(A)** and **(C)** are spot plates of 10x dilution series of *Saccharomyces cerevisiae* expressing individual candidate effectors from a high-copy galactose inducible vector (pYES-DEST52). Left plates show growth on DOB agar containing 2% glucose where the genes of interest are off and serve as negative controls (uninduced). Right plates are DOB agar with 2% galactose for induction of the genes of interest. Lack of growth on these plates indicates that the expressed gene is toxic to yeast growth. For plates in panels **(A)** and **(C)**, ‘s’ indicates the predicted signal peptide was removed (-) from (CAHE_0406, CAHE_0405, CAHE_p0043, and CAHE_0757) or added (+) to the construct (CAHE_p0044) and ‘*’ indicates the construct was codon optimized for *S. cerevisiae*. **(B)** and **(D)** show proteomic detection of the proteins heterologously expressed in *S. cerevisiae* when grown on glucose (uninduced) and galactose (induced) media. Each point is the abundance (TSS normalized AUC quantification) of the protein in one replicate and horizontal lines indicate the mean of three replicates.

When expressing CAHE_0406 in *S. cerevisiae* from the high-copy plasmid pYES-DEST52 under control of the galactose-inducible promoter GAL1, we observed reduction in yeast growth compared to the empty vector control, but not complete lethality as observed in the *w*Pip *cidB* positive control (**Figure 2A)**. We did not observe reduction in yeast growth associated with expression of the other unaltered candidates (CAHE_0405, CAHE_p0044, CAHE_p0043, and CAHE_0757) (**Figure 2A,C**). We detected expression of all unaltered candidates in batch cultures when induced, except for CAHE_p0044 (**Figure 2B,D**). All candidate protein sequences included a predicted signal peptide, except CAHE_p0044, which notably did not encode a signal peptide in the predicted sequence. To mimic potential maturation of secreted *Cardinium* effectors through cleavage of signal peptides, a common occurrence in the maturation process of secreted bacterial proteins (Beckwith, 2013), we generated additional constructs with predicted signal peptides removed. We observed no change in yeast growth for CAHE_0405 and CAHE_0757 when we removed their signal peptides compared to full length constructs (**Figure 2C,D)**. The removal of the predicted signal peptide for CAHE_0406 resulted in a total loss of toxicity when expressed in *S. cerevisiae*, resulting in growth similar to that of the empty vector control; however, we were not able to detect CAHE_0406 without the signal peptide with proteomics, likely due to the small size of the protein (**Figure 2A,B)**. Strikingly, we observed strong toxicity in *S. cerevisiae* when CAHE_p0043 was expressed without its signal peptide (CAHE_p0043^-signal^), similar to *Wolbachia w*Pip *cidB* (**Figure 2A**). Adding the predicted upstream signal peptide to CAHE_p0044 did not alter the protein’s impact on yeast growth when expressed (**Figure 2C,D**). Reduced growth in yeast associated with expression of full-length CAHE_406 and CAHE_p0043^-signal^ suggests that these proteins are strong candidates for CI induction by *Cardinium c*Eper1.

We attempted to assess the potential of CAHE_0405 and CAHE_p0044 as possible rescue factors for CAHE_406 and CAHE_p0043^-signal^, respectively, based on their proximity to the putative modification factors on the *Cardinium* genome and plasmid using a similar approach to that employed to elucidate *Wolbachia* CI rescue factors (Beckmann et al., 2019, 2017; Murphy and Beckmann, 2024; Sun et al., 2022; Xiao et al., 2021). However, our results were inconclusive as to what effect co-expression of the CI candidates with their neighboring gene has in this yeast model (**Figure S7**).

### Presence of CI effector candidates across *Cardinium* strains

Multiple *Cardinium* strains can induce CI in their host, so we searched for homologs of the putative CI effectors in other strains to assess conservation of the effector candidates across *Cardinium* strains and gather potential evidence of involvement in CI. We used BLASTp to search for full-length or nearly full-length homologs of CI candidate proteins identified by proteomics (CAHE_p0043, CAHE_p0044, CAHE_0405, CAHE_0406, and CAHE_0757) in 25 *Cardinium* genomes and the sister taxon to *Cardinium*, *Candidatus* Amoebophilus asiaticus 5a2 (**Table S1**). We found that CAHE_0757 exhibited broad distribution across multiple *Cardinium* genomes with and without known manipulation phenotypes (**Figure 3A**), albeit with a low amino acid sequence identity of <30% in most cases. Notably, a CAHE_0757 homolog is missing from *c*Sfur, a known CI-causing strain found in the planthopper *Sogatella furcifera*, suggesting that CAHE_0757 is not involved in CI in *S. furcifera*.

**Figure 3.**
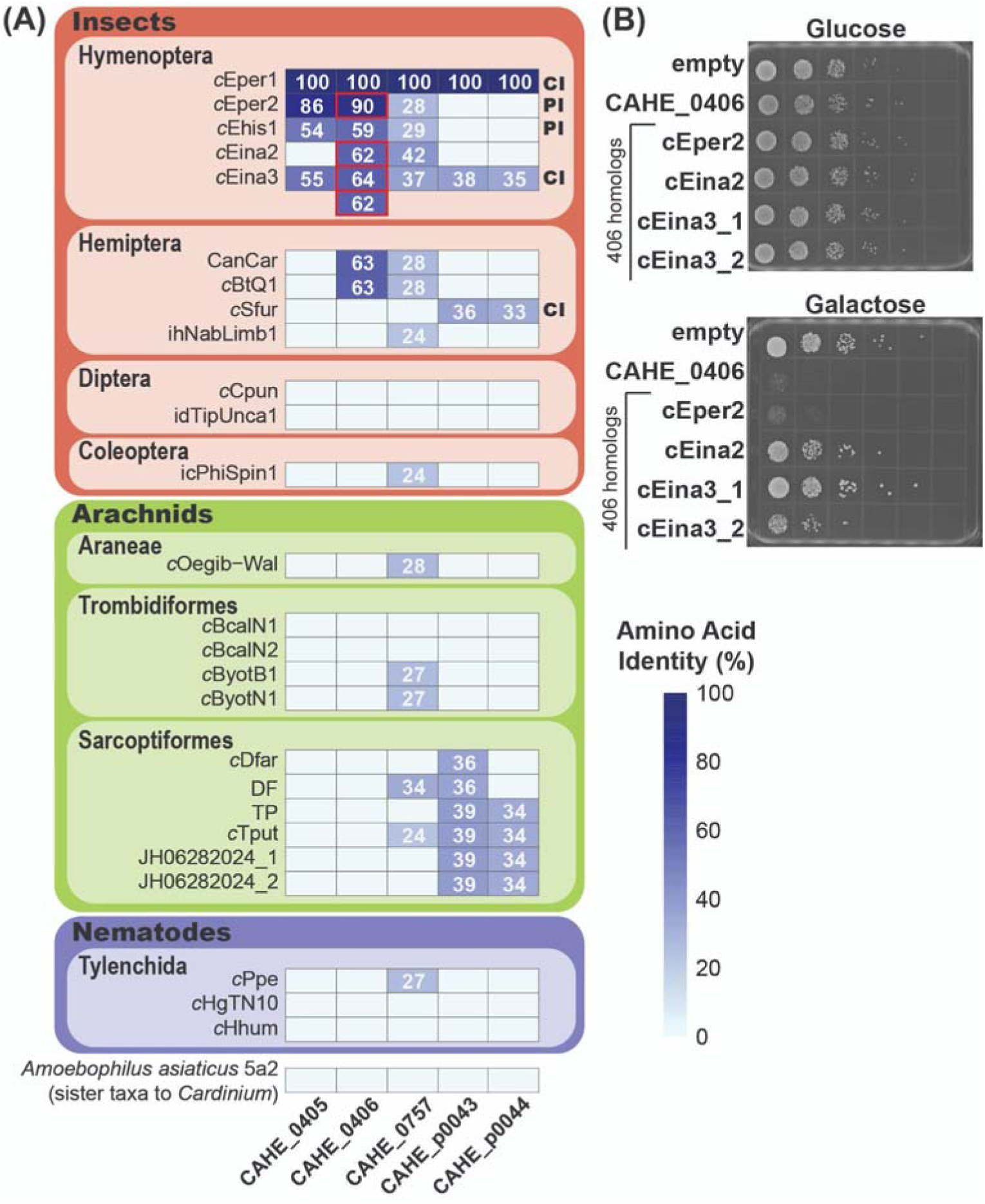
Identification and screening of CAHE_0406 homologs. **(A)** Amino acid identities (%) of full-length homologs of *c*Eper1 CI candidate proteins in other *Cardinium* genomes. Strains are grouped according to host taxonomy. There were two homologs (64% and 62% amino acid identity) for CAHE_0406 in CI-causing *c*Eina3. Known reproductive manipulation phenotypes are indicated by initials: cytoplasmic incompatibility (CI) and parthenogenesis (PI). Homologs selected for further screening are indicated by red framing. **(B)** Images are of spot plates of 10x dilution series of *Saccharomyces cerevisiae* expressing individual candidate effectors from a high-copy galactose inducible vector (pYES-DEST52). Top plate shows growth on DOB agar + 2% glucose where the genes of interest are not induced and serve as negative controls. Bottom plate is DOB agar with 2% galactose for induction of the genes of interest. Lack of growth on these plates indicates that the expressed gene is toxic to yeast growth.

In contrast, we found full-length homologs of both CAHE_p0043 and CAHE_p0044 in the only two other known CI strains, *c*Eina3 and *c*Sfur with published genomes (**Figure 3A**). We also identified full-length homologs of CAHE_p0043 and CAHE_p0044 in four genomes of *Cardinium* infecting *Tyrophagus putrescentiae* mites and homologs to CAHE_p0043 in *Cardinium* strains DF and *c*Dfar infecting the house dust mite *Dermatophagoides farinae* (**Figure 3A**). These mite-associated *Cardinium* strains have not been characterized for reproductive manipulation phenotypes. We found full-length homologs of CAHE_0406 only in *Encarsia*- and *Bemisia tabaci*-associated *Cardinium* genomes. We only found homologs of CAHE_0405, the neighboring gene of CAHE_0406, in the genomes of three *Encarsia*-associated strains (*c*Eper2, *c*Ehis1, and *c*Eina3). All genomes encoding homologs of both CAHE_0405 and CAHE_406 are associated with reproductive manipulation phenotypes (*c*Eper2: Parthenogenesis Induction (PI), *c*Ehis1: PI, and *c*Eina3: CI).

Due to their high similarity to CAHE_0406, we expressed the homologs of CAHE_0406 encoded by *c*Eper2 (WP_432451371.1) and *c*Eina2 (WP_432446694.1), and two homologs encoded by *c*Eina3 (WP_432466820.1 with 62% amino acid identity to CAHE_0406 and WP_432466821.1 with 64% amino acid identity to CAHE_0406) in yeast to screen for toxicity. We did not screen the protein encoded by *c*Ehis1 with 59% amino acid identity to CAHE_0406 (**Figure 3A**) because of its length (149 aa versus 60 aa). When expressed in yeast, two of the homologs did not impact yeast growth compared to the empty control: WP_432446694.1 in the asymptomatic *c*Eina2 strain and WP_432466820.1 in the CI strain *c*Eina3 (*c*Eina3_1) (**Figure 3B**). The homolog WP_432466821.1 in the CI strain *c*Eina3 (*c*Eina3_2) caused a minor reduction in yeast growth compared to the empty control, though notably less than did CAHE_0406 (**Figure 3B**). The highly similar (90%) WP_432451371.1 homolog encoded by the PI strain *c*Eper2 showed similar lethality to CAHE_0406 when induced (**Figure 3B, Figure S8**).

### Localization of candidate CI proteins in *S. cerevisiae* suggests interactions with eukaryotic **nuclei**

To determine if CAHE_p0043^-signal^ and CAHE_0406 toxicity is indicative of CI-involvement, we observed the localization of CAHE_0406 and CAHE_p0043^-signal^ proteins fused with C-terminal enhanced green fluorescent protein (eGFP) when expressed in *S. cerevisiae* across 2-8 hours of induction by fluorescence microscopy (**Figure 4A-C**), as previously implemented to screen *Wolbachia* effectors in yeast cells (Rice et al., 2017). We expected CAHE_p0043^-signal^+eGFP and CAHE_0406+eGFP to localize to the *S. cerevisiae* nucleus or cause similar cytological defects to those observed in naturally CI-affected *E. suzannae* embryos (Gebiola et al., 2017).

**Figure 4.**
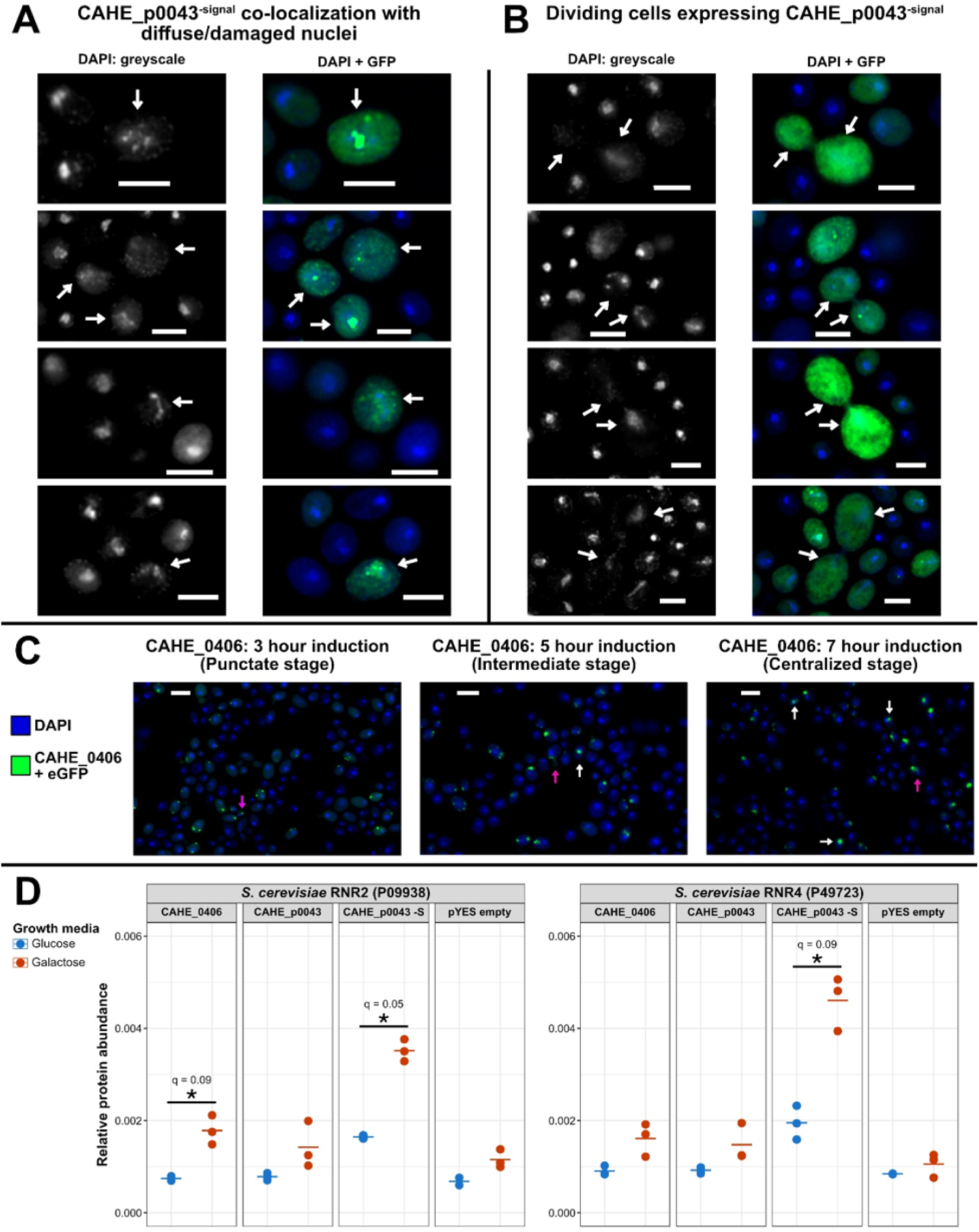
Localization of CAHE_p0043 and CAHE_0406 CI candidates fused with eGFP and RNR expression response in *S. cerevisiae*. **(A)** Example images of CAHE_p0043^-signal^ co-localizing with yeast nuclei which appear diffuse or damaged. **(B)** Examples of dividing *S. cerevisiae* cells expressing CAHE_p0043^-signal^. For **A** and **B**, cultures were induced for 6 to 8 hours, white arrows indicate cells or features of interest, and white bars at the bottom of each image indicate a scale of 3 µm. **(C)** Progression of CAHE_0406 localization across hours 3 to 7 of induction. Pink arrows indicate cells which are dividing or have recently divided, while white arrows indicate instances of CAHE_0406 and DAPI co-localization. White bars at the top of images indicate a scale of 5 µm. **(D)** Proteomic detection of ribonucleoside-diphosphate reductase (RNR) proteins in *S. cerevisiae* with *c*Eper1 constructs (without eGFP tag) induced (red) and uninduced (blue). “-S” indicates the signal peptide was removed. Protein abundance was quantified using an area under the curve method and normalized via total sum scaling. Statistical significance (* indicates q < 0.1) was determined by performing pairwise Welch’s T-tests on log2 transformed data corrected for multiple hypothesis testing with Benjamini-Hochberg FDR correction.

Localization of CAHE_0406+eGFP in yeast cells did not display consistent targeting to the eukaryotic nucleus (**Figure 4C**). Instead, early (2-5 hours induction) CAHE_0406+eGFP localization is highly punctate within a cell, with the intensity and number of separate spots increasing with longer incubation periods. At 6 hours of induction, CAHE_0406+eGFP localizes to a larger centralized area of eGFP fluorescence with variable subcellular localization (**Figure 4C**).

Localization of CAHE_p0043^-signal^+eGFP was distinct from CAHE_0406+eGFP. We observed limited fluorescence of CAHE_p0043^-signal^+eGFP prior to 6 hours of induction. Between 6 and 8 hours of induction, CAHE_p0043^-signal^+eGFP was often observed diffusely throughout the yeast cell cytoplasm, creating a hazy appearance usually consisting of many small dots of fluorescence with varying intensities. CAHE_p0043^-signal^+eGFP was sometimes present in condensed bright areas, similar to CAHE_0406+eGFP localization post-7-hour induction (**Figure 4A,C**). In the case of CAHE_p0043^-signal^+eGFP, we often observed these hotspots to be co-localized with, or with what appeared to be the remains of, the yeast nucleus.

In addition to the distinct localization pattern of CAHE_p0043^-signal^+eGFP, we noticed many yeast cells with diffuse, misshapen, fragmented, or completely absent nuclei in 6 to 8-hour CAHE_p0043^-signal^+eGFP induction cultures (**Figure 4A**; e.g., typical nuclei at 3, 5, or 7-hour induction timepoints of CAHE_0406, **Figure 4C**). In most cases, irregular nuclei occurred in cells which also had fluorescence signal from CAHE_p0043^-signal^+eGFP, and we found multiple instances of condensed eGFP fluorescence near or overlapping with irregular nuclei. Yeast cells undergoing cell division and producing CAHE_p0043^-signal^+eGFP exhibit irregularities suggestive of unsuccessful cell division, and we rarely observed normal dividing cells producing CAHE_p0043^-signal^+eGFP signal (**Figure 4B**). In a few instances, we observed abnormally large budding cells where the budding cell contained a diffuse DAPI signal and its large daughter cell was completely devoid of DAPI signal (**Figure 4B**). In another case, both the daughter and parent cells contained DAPI signal, but both nuclei appeared fragmented instead of consisting of single condensed areas of DAPI signal (**Figure 4B**).

Proteomics of yeast producing CAHE_p0043^-signal^ revealed enrichment of ribonucleoside-diphosphate reductase (RNR) proteins RNR2 and RNR4 compared to uninduced conditions (q < 0.1, Welch’s T-test with Benjamini-Hochberg FDR correction; **Figure 4D**), as well as MEC1-mediated checkpoint protein HUG1 which was only detected in yeast carrying the CAHE_p0043^-signal^ construct (**Figure S9**). Increased expression of these genes indicates DNA damage (see discussion). We also observed enrichment of RNR2 along with FMP52 and FMP45 in the yeast proteome when CAHE_0406 was expressed (**Figure S9**).

## Discussion

We applied complementary approaches of proteomics and heterologous yeast expression to identify *Cardinium*-produced effectors of the *c*Eper1 strain responsible for CI induction in *E. suzannae*. We found two proteins, CAHE_0406 and CAHE_p0043^-signal^, that both showed distinctive proteomic detection patterns consistent with our predicted model for *Cardinium* CI and were toxic when heterologously expressed in *S. cerevisiae* (**Table 1**). The cytology of CAHE_p0043^-signal^ in yeast showed disruptive interactions between the protein and the nuclei of yeast cells, and two yeast ribonucleoside-diphosphate reductase (RNR) proteins associated with DNA damage were upregulated upon CAHE_p0043^-signal^ expression, suggesting involvement in CI. Combined, these results suggest that one or both of these proteins contribute to *Cardinium*-host cell interactions and possibly *Cardinium*-induced CI.

**TABLE 1.** Evidence for putative cytoplasmic incompatibility (CI) effectors identified in this study. Bolded ‘Yes’ indicates evidence supporting a role in CI. ‘N/A’ indicates there is no applicable evidence.

| Evidence | CAHE_p0044 | CAHE_p0043 | CAHE_0406 | CAHE_0405 | CAHE_0757 |
| --- | --- | --- | --- | --- | --- |
| Size (AA length) | 200 | 312 | 60 | 149 | 681 |
| Domains/Annotation | Knr4/Smi1-like domain | prokaryotic lipoprotein | prokaryotic lipoprotein | prokaryotic lipoprotein | prokaryotic lipoprotein |
| Possible partner | CAHE_p0043 | CAHE_p0044 | CAHE_0405 | CAHE_0406 | none predicted |
| Secreted based on proteome | **Yes** | **Yes** | No | No | No |
| Pupal testes detection | **Yes** | **Yes** | **Yes** | **Yes** | **Yes** |
| Seminal vesicle detection | No | No | **Yes**<sup>1</sup> | No | **Yes** |
| Absence in non-reproductive organs | No | No | No | No | No |
| Toxicity in yeast | No | **Yes**, without signal peptide | **Yes**, with signal peptide | No | No |
| Localization in yeast | N/A | co-localizes with disrupted nuclei (-signal) | no nuclear localization | N/A | N/A |
| Yeast DNA damage | N/A | **Yes**, distinct cytology with RNR2, RNR4, and HUG1 upregulated | Unclear from cytology but RNR2, FMP45, and FMP52 upregulated | N/A | N/A |

<sup>1</sup>Based on detection in 1 replicate out of 4

CAHE_0406 and CAHE_p0043^-signal^ were both toxic to yeast and exhibited genomic structure and expression patterns suggestive of CI involvement. CAHE_0406 was among the 20 most abundant *Cardinium* proteins detected in pupal testes (**Figure 1**), the life stage and tissue where *Cardinium* CI induction occurs in *E. suzannae* (Doremus et al., 2020). CAHE_p0043 was also detected in pupal testes and exhibited a detection pattern indicative of secretion (**Figure 1**). Both CAHE_0406 and CAHE_p0043 have a neighboring gene that was also detected (**Figure 1**, **S4**, **S5**). We used paired genes as part of our selection criteria based on the current models of *Wolbachia* CI which rely on two neighboring *cif* genes for induction and rescue of the phenotype (Beckmann et al., 2019; Shropshire and Bordenstein, 2019; Wang et al., 2022). These genes CAHE_0406 and CAHE_p0043 also have homologs in *Cardinium* strains of other host taxa **(Figure 3)**. Another common genomic feature of *Wolbachia cif* genes is association with prophage and other mobile genetic elements (Amoros et al., 2025; LePage et al., 2017). While neither of the two *Cardinium* CI candidates is associated with a prophage, which are absent in *Cardinium* (Mann et al., 2017), CAHE_p0043 is encoded on the *c*Eper1 plasmid amongst transposases (**Figure S5**).

There are only few commonalities between *Wolbachia cif* genes and the two *Cardinium* CI candidates. CAHE_0406 and CAHE_p0043 are enriched in reproductive tissues, as expected, cause toxicity in yeast, and, like *Wolbachia cif* genes, are located on the genome or plasmid next to another enriched protein that is a candidate for a rescue factor. There the similarity appears to end. CAHE_0406 (60 amino acids) and CAHE_p0043 (312 amino acids) are much smaller than *Wolbachia cif*s (e.g., *w*Pip CidB at 1200 amino acids and CidA at 500 amino acids) (Sicard et al., 2021). They also show no homology to *Wolbachia cifs*. Both lack the known functional domains of *Wolbachia cifs*, such as nuclease and deubiquitylase domains (Beckmann et al., 2017; Chen et al., 2019). Taken together, the distinctive size and lack of predicted functional domain overlap between *Wolbachia cifs* and our candidates, CAHE_0406 and CAHE_p0043, suggest that these proteins may induce CI through a different mechanism or interact with different host targets. CAHE_0406 and CAHE_p0043 are predicted lipoproteins. Bacterial lipoproteins, which are typically covalently bound to the outer membrane of the cell after signal peptide cleavage (Kovacs-Simon et al., 2011; Smithers et al., 2021), have been implicated in host-interaction effects including immune evasion and modulation, and cell adhesion and invasion (Cole et al., 2021; Wilson and Bernstein, 2016). Symbiont enrichment-specific detection of CAHE_0406 in our proteomic data supports membrane localization of this protein, though CAHE_0406 requires its predicted signal peptide for toxicity in yeast (**Figure 2**; **Figure S6**). The best match from the Swiss-Prot database and closest match in amino acid length for CAHE_0406 when performing a BLASTp search against the UniProtKB + Swiss-Prot database is an 86 amino acid Omega-theraphotoxin-Hhn1f3 (identity = 38.2%, score = 76, E-value = 2.1; matrix = BLOSUM62, gapalign = true; **Table S2**) of *Cyriopagopus hainanus* (Chinese bird spider), which belongs to the Huwentoxin-1 family and acts through inhibiting ion-channels in the membrane of target cells (Liu et al., 2003; Peng et al., 2002). The shared homology between CAHE_0406 and this toxin, which primarily overlaps with the signal peptide region, suggests that CAHE_0406 may require the signal peptide for localization in the host cell as a part of its mechanism of action, and thus removal of the lipoprotein signal peptide-like region from the protein may inhibit activity. CAHE_0406+eGFP localization when expressed in *S. cerevisiae* suggested that the protein may be packaged into vesicles over time as it appears to transition from a punctate presence throughout the cell cytoplasm to condensing into single compact areas of protein with inconsistent subcellular localization, similar to the localization of autophagosomes or other membrane-bound vesicles (Gross et al., 2024; Torggler et al., 2017). Based on this localization pattern, it seems that CAHE_0406+eGFP does not regularly localize to or interact with the *S. cerevisiae* nucleus. However, it may be that CAHE_0406 interacts with other central pathways in host cells, such as membrane-trafficking (Berardi et al., 2025; Carpinone et al., 2018), in a way that slows cell growth and proliferation. Proteomic results showing increased expression of autophagy protein ATG8 and reticulons RTN1 and RTN2 in yeast under CAHE_0406 expression seem to support this (**Figure S9**). Reticulons are major structural regulators of the endoplasmic reticulum (ER) and are suggested to regulate ER inheritance in yeast (Piña et al., 2016). ATG8 has been shown to increase in expression under ER stress induced autophagy in yeast (Yorimitsu et al., 2006), though it is typically associated with starvation stress. Interestingly, despite CAHE_0406 not appearing to localize to yeast nuclei, the proteomic results show increased expression of FMP45 and FMP52 when CAHE_0406 is expressed (**Figure S9**), suggesting DNA damage may be occurring. Specifically, FMP45 has been shown to exhibit increased expression in yeast treated with a double-strand break inducer (Cong et al., 2025) and FMP52 expression has been shown to respond to a photolesion inducer (Dardalhon et al., 2007), though the molecular roles of these specific proteins in DNA damage response remain unclear. Follow-up studies are needed to determine which eukaryotic proteins and pathways CAHE_0406 interacts with and how it produces a toxic effect in *S. cerevisiae*.

In contrast, our data suggests that CAHE_p0043 is a fully secreted protein, not attached to *Cardinium*, and requires removal of the predicted signal peptide for toxicity (**Figure 1, 2**). CAHE_p0043 has few useful annotations through BLASTp as it matches mostly to much larger proteins and to no proteins reviewed through Swiss-Prot. Interestingly, many of these larger proteins appear to be involved with the cell cycle or nucleic acid interaction based on protein family and domain predictions (e.g., AAA ATPase family proteins, a core motif found in CifB proteins (Amoros et al., 2025; Martinez et al., 2021), and protein C3H1-type domains (Carballo et al., 1998; Lai et al., 1999); **Table S2**). However, CAHE_p0043 does not appear to share these domains or protein family designations. Most of what we can use to speculate on the possible mechanism of CAHE_p0043 is the toxicity of the protein when expressed in yeast after signal peptide removal (**Figure 2**), its predicted and observed nuclear localization (**Figure 4; Table S3**), and the apparent secretion of the protein from symbiont cells (**Figure 1**) which we might expect involves cleavage of the signal peptide leading to secretion of the mature protein (Beckwith, 2013). This suggests that both signal peptide cleavage and, perhaps, specific subsequent eukaryotic cell trafficking of the protein is critical for CAHE_p0043 activity.

Prediction of CAHE_p0043^-signal^ eukaryotic cell localization using DeepLoc2.1 (Ødum et al., 2024; Thumuluri et al., 2022) revealed increased probability of nuclear localization and a nuclear export signal, which were not predicted for CAHE_p0043 with the signal peptide (**Table S3**). This suggests that removal of the signal peptide during secretion of the protein possibly allows for nuclear localization. When expressed in *S. cerevisiae*, we observed bright, condensed areas of CAHE_p0043^-signal^ which often co-localized with the DAPI signal of yeast nuclei or with what appeared to be the remains of yeast cell nuclei. We also noticed that dividing yeast cells expressing CAHE_p0043^-signal^ were very rare and had abnormalities ranging from diffuse and fragmented nuclei to large daughter cells with no DNA. Further, proteomics of induction assays in yeast revealed enrichment of ribonucleoside-diphosphate reductases (RNRs) in *S. cerevisiae* expressing CAHE_p0043^-signal^. These proteins are responsible for catalyzing the rate-limiting step in the production of deoxyribonucleotides (Sanvisens et al., 2013). Upon DNA damage in yeast, the synthesis of deoxyribonucleotides is increased 6-8 fold via upregulation of RNR proteins (Bu et al., 2019; Chabes et al., 2003). RNR2 and RNR4 are both known to increase in abundance in response to DNA damage, and both were significantly enriched when CAHE_p0043^-signal^ was expressed (**Figure 4**) (Elledge and Davis, 1989; Gasch et al., 2001; Huang and Elledge, 1997). We also observed increased expression of HUG1 in response to CAHE_p0043^-signal^ expression (**Figure S9**). HUG1 is a proposed RNR-interacting protein that is typically induced by DNA damage in yeast and has been previously used as a genotoxicity reporter in screening assays (Kim and Siede, 2011; Lee et al., 2007; Wei et al., 2013). Together, localization and proteomics results suggest that the toxic effect of CAHE_p0043^-signal^ may be due to interaction with or disruption of the eukaryotic nucleus and damage to chromatin, potentially leading to failed cell division (**Figure 4**), as we might expect for CI.

Three of our candidates (CAHE_0757, CAHE_0405, and CAHE_p0044) did not impact yeast growth (**Figure 2**; **Table 1**). CAHE_0757, a predicted lipoprotein, was consistently detected in high abundance across host sex, life stage, and tissue. These detection patterns were similar to highly abundant housekeeping proteins like GroL. The housekeeping-like detection patterns combined with the lack of toxic phenotype when expressed in yeast suggest that this protein is not involved in CI induction and is more likely a central component of *Cardinium* physiology or host interaction. CAHE_0405 and CAHE_p0044 share similar proteomic detection patterns and genomic occurrence across *Cardinium* lineages with their neighboring toxicity-inducing genes, CAHE_0406 and CAHE_p0043 (**Figure 1, S1, & S6**). The shared detection patterns of genes in each pair suggest that interactions within the pairs may be conserved, and therefore CAHE_0405 and CAHE_p0044 should be considered as possible candidates for CI involvement or potential rescue factors. Interestingly, CAHE_p0044 contains a predicted SMI1/KNR4 domain. The SMI1/KNR4 domain was characterized in a protein involved in fungal cell wall stress tolerance (Kroll et al., 2025; Martin-Yken et al., 2002). This domain has also been suggested to be involved in detoxifying nucleic acid degrading toxins (Zhang et al., 2011). We attempted rescue assays as done in *Wolbachia* studies (Beckmann et al., 2017; Murphy and Beckmann, 2024; Sun et al., 2022); however, we were unable to achieve rescue (**Supporting Information; Figure S7**).

In addition to the CI candidate proteins assessed in this study, we detected nearly 24% of the 887 predicted protein coding sequences available in UniProt (downloaded: 29 Dec 2020) for *Cardinium c*Eper1 through our differential proteomics strategy, providing valuable insights into symbiont physiology. We identified additional proteins potentially involved in host interactions and possibly CI (**Table S4**). These include CAHE_0669 (predicted peptidyl-prolyl sis-trans isomerase), CAHE_0435 (ankyrin repeat-containing protein), CAHE_0706 (collagen triple helix repeat-containing protein), CAHE_0705 (uncharacterized putative lipoprotein), and components of the type VI secretion system (CAHE_0458, CAHE_0459, CAHE_0460, CAHE_0036, and CAHE_0409; **Figure S2**). Discussion about these proteins can be found in the **Supporting Information**.

### Future work and outlook

Here, we provide evidence for involvement of two *Cardinium* proteins in CI. Future work is needed to explore the structure and mechanism of CAHE_0406 and CAHE_p0043 to verify and understand their role in CI. One question is how these effectors are delivered from infected males to uninfected embryos and another is the timing of their effects on gametes and developing embryos, which are also active questions in *Wolbachia* CI research (Beckmann et al., 2017; Horard et al., 2022; Wang et al., 2022). One approach to interrogate this in *Encarsia* could be through proteomic measurements of spermathecae, which are female organs for the storage of sperm until fertilization. Detection of wPip_0282 (*Wolbachia cif*) in mosquito spermathecae was early evidence for its involvement in CI (Beckmann and Fallon, 2013), and detection of CAHE_0406 or CAHE_p0043 in the spermathecae would be evidence of effector transmission by sperm. Emerging low input and single-cell proteomics techniques will be imperative for gaining insights into tissue-specific physiologies of *Cardinium* and transmission dynamics of CI effectors in its tiny insect host. Another remaining question is how the toxic effects of CAHE_0406 and CAHE_p0043 are reversed upon fertilization of an infected egg. Although we attempted to rescue toxicity effects of CAHE_0406 and CAHE_p0043 on yeast by co-expressing their neighboring genes (CAHE_0405 and CAHE_p0044, respectively; **Supporting Information, Figure S7**), our results were inconclusive. It is possible that these proteins, or their mechanisms of action, are not entirely compatible with the yeast-based approaches used to elucidate *Wolbachia* CI rescue. For this reason, future research should use other molecular methods (e.g., transgenic expression in insect lines) to interrogate such mechanisms (Chaverra-Rodriguez et al., 2020; Werren et al., 2009; Yang et al., 2025; Zhang et al., 2024). Lastly, more work is required to identify potential molecular host targets of CAHE_0406 and CAHE_p0043. Follow-up work should determine if CAHE_p0043^-signal^ degrades DNA *in vitro* and identify *in vivo* host targets (Beckmann et al., 2017; Chen et al., 2019; Xiao et al., 2021) to further elucidate the mechanism of *Cardinium* CI in *E. suzannae*.

In summary, our work presents the first *Cardinium* proteomic data and CAHE_0406 and CAHE_p0043^-signal^ as CI factor candidates of *Cardinium* based on homology patterns across *Cardinium* lineages, detection of these proteins across host sex, life stage, and tissue, and ability of these proteins to disrupt yeast growth. These candidates are entirely distinct from the *Wolbachia cif* genes, supporting previous proposals of convergence and, likely, distinctive mechanisms of the CI phenotype in these distantly related symbiont lineages. Our findings provide a broader basis for the exploration of CI across symbiont taxa and suggest remarkable convergence of the CI phenotype among symbiont lineages.

## Materials and Methods

### Insect lines and verification of infection status

*Encarsia suzannae* is a minute (<1 mm long) parasitoid wasp that develops within whiteflies. *Cardinium*-carrying *E. suzannae were* originally collected from South Texas, USA, and were maintained at The University of Arizona in Tucson, AZ, USA at 27°C with a 16h:8h day:night light cycle. Female *E. suzannae* were raised on whiteflies (*Bemisia tabaci*) on cowpea (*Vigna unguiculata*), while males, which develop as hyperparasitoids (parasitic on other developing parasitoids), were raised on another whitefly parasitoid, *Eretmocerus emiratus* (Hunter et al., 2003). Neither *B. tabaci* nor *E. emiratus* were infected with *Cardinium*. Cured wasp lines were established by treating infected wasps with rifampicin and maintained for >10 generations prior to use in experiments (Hunter et al., 2003). To confirm the presence of *Cardinium* in *E. suzannae* cultures, we extracted wasp and symbiont DNA using previously established protocols for Chelex extractions (Doremus et al., 2020; Hunter et al., 2003). Symbiont presence was confirmed with *Cardinium* diagnostic primers and the PCR product was visualized on a 1% agarose gel (Harris et al., 2010).

### Differential proteomics

We compared male and female, pupal and adult, and reproductive organs and heads using differential proteomics to identify symbiont proteins associated with specific host physiologies and CI. Our detailed proteomics methods can be found in the **Supporting Information.**

To compare symbiont proteins between sexes and life stages, we prepared 4-5 replicates each of symbiont-carrying adult males, pupal males, adult females, and pupal females, with 100 live wasps per replicate (**Table S5**). Pooling was necessary to ensure sufficient biomass for sample preparation. We thoroughly homogenized live wasps in phosphate buffered saline (PBS; 0.01 M phosphate, 0.0027 M KCl, and 0.137 M NaCl, pH 7.4) with a protease inhibitor cocktail (cOmplete EDTA-free Protease Inhibitor Cocktail, Roche) using a glass homogenizer (KIMBLE DUALL Tissue Grinder, DWK Life Sciences) before performing differential pelleting to fractionate homogenates based on previously described methods (Hinzke et al., 2018). This approach yielded 3 fractions: a cell debris/heavy pellet (P), a *Cardinium* enrichment pellet (PS), and a supernatant (S) (**Figure S10**). The S fraction was lyophilized overnight to concentrate. The P, PS, and S fractions were suspended in an SDT lysis buffer (4% w/v sodium dodecyl sulfate, 100 mM Tris-HCl pH 7.6, 0.1 M dithiothreitol) and heated at 95°C for 10 minutes to lyse cells and denature proteins. To compare proteins across host tissues, we dissected wasps in PBS to harvest heads (to serve as a control against the reproductive tissues) and reproductive organs (testes+seminal vesicle or ovaries) for proteomic measurement. We pooled organs suspended in SDS lysis buffer (4% w/v sodium dodecyl sulfate, 100 mM Tris-HCl pH 7.6) to obtain four to five replicates of each tissue sample with 50-75 tissues in each. We homogenized head samples to disrupt the insect cuticle prior to further processing. Organ samples were then heated at 95°C for ten minutes to lyse cells and denature proteins. We loaded sample lysates of fractionated wasps and organ samples onto 10 kDa PES centrifugal filter units (VWR) to prepare peptide eluates using filter aided sample preparation protocol (Wiśniewski, 2019) with modified centrifugation times.

We used nanoflow liquid chromatography and tandem mass spectrometry (LC-MS/MS) to measure proteomic contents of samples (Mordant and Kleiner, 2021; Violette et al., 2024) (**Tables S6-S8**). Resulting raw spectral data were searched using ProteomeDiscoverer 2.3 (Thermo Scientific), as previously described (Mordant and Kleiner, 2021), against a custom database constructed from protein sequences predicted from the *c*Eper1 genome (UniProt: UP000003996) and *E. suzannae* transcriptome of male and female adult wasps (Schultz et al., 2022) and pupae, as well as potential contaminants (https://www.thegpm.org/crap/). We constructed the database and reduced redundancy using CD-HIT following previous recommendations (Blakeley-Ruiz and Kleiner, 2022; Fu et al., 2012; Li and Godzik, 2006). UniProt Accessions, Locus Tags, and GenBank identifiers for all *c*Eper1 proteins can be found in **Supplementary Data 3**. We quantified proteins by peptide spectral match (PSM) counts and merged search results using custom python scripts, filtering for proteins inferred with 5% FDR confidence and classified as a master protein. We identified differentially abundant symbiont proteins between host sexes, life stages, fractions, and tissues using differential statistics. We used normalization (NSAF or orgNSAF), filtering, transformation, and imputation strategies appropriate to specific comparisons (Florens et al., 2006; Kleiner, 2017; Mueller et al., 2010). A detailed overview of all statistical comparisons performed can be found in **Supplementary Data 4** and **5**. Because of the low number of replicates, we assessed significance using an alpha value of 0.1 in this study. We analyzed our proteomic data using R (version 4.2.2) using code that we adapted, in part, from code generated through the use of ChatGPT, Google Gemini, and various collaborative programming blogs for the purpose of data cleaning, high throughput code adaptation (e.g., for loops), figure cleaning, and general error debugging. Images, data figures generated in R, and other visualization tools were formatted using Adobe Illustrator. For an overview of data analysis tools, packages, and software used in this manuscript see **Table S9**.

### Heterologous expression of *Cardinium* proteins in yeast

We heterologously expressed five *c*Eper1 protein candidates (CAHE_0757, CAHE_0405, CAHE_0406, CAHE_p0043, and CAHE_p0044) in *Saccharomyces cerevisiae* and compared toxicity associated with expression of candidate genes to an empty vector (negative control) and to the *Wolbachia w*Pip *cidB* CI gene (WP0283 / CAQ54391.1) toxicity (positive control). For this, the candidate genes and the *w*Pip *cidB* gene were synthesized by gene synthesis (ThermoFisher GeneArt) and subcloned into the Gateway cloning entry vector pDONR221 (Katzen, 2007). We used Invitrogen Gateway LR Clonase II to integrate the synthesized genes into Gateway compatible expression plasmids from their entry vector **(Table S10)** (Alberti et al., 2007), following the manufacturer’s protocol with an extended incubation period (2-5 hours).

We transformed reaction products into chemically competent ccdB-sensitive StrataClone *Escherichia coli* (Agilent) via heat shock following the manufacturer’s protocol. To isolate *E. coli* transformants, we plated on LB media containing 100 µg/mL ampicillin and tested colonies for susceptibility to LB containing 30 µg/mL chloramphenicol to screen for gene of interest insertion. We confirmed insert size with PCR and agarose gel electrophoresis, and further confirmed transformants with Sanger sequencing. The plasmids of *E. coli* transformants containing the correct insert were purified using the Invitrogen Quick Plasmid Miniprep kit. We then transformed those plasmids into *S. cerevisiae* strain BY4741 (MATa, his3Δ1, leu2Δ0, met15Δ0, ura3Δ0) using lithium acetate pretreatment and electroporation as previously described (Thompson et al., 1998). Yeast transformants were grown on drop-out base (DOB) medium containing 2% glucose (MP Biomedicals) supplemented with synthetic complete (SC) mixture (MP Biomedicals) lacking uracil (for strains transformed with pYES) and/or leucine (for strains transformed with pAG415). We selected and screened colonies for the correct size insert via PCR as outlined above. Yeast coexpression strains were generated by sequential transformation: one plasmid was transformed into *S. cerevisiae* BY4741 and confirmed, followed by transformation of the second plasmid into the singly transformed yeast strain and plated onto media lacking both leucine and uracil. The presence of both plasmids was confirmed via PCR and growth on media lacking leucine and uracil. All *S. cerevisiae* transformants were grown at 30°C, unless otherwise noted. Additional information on the generated yeast constructs can be found in **Table S11** and the methods can be found in the **Supporting Information**.

CI candidate genes CAHE_0757, CAHE_0405, CAHE_0406, and CAHE_p0043 were predicted to encode lipoprotein signal peptides (Sec/SPII) via SignalP 6.0 in “slow” mode (Teufel et al., 2022). To account for possible change in function following signal peptide cleavage, we generated additional constructs of these four genes without their predicted signal peptides. For each gene, we removed the signal peptide region using the predicted cleavage site output by SignalP 6.0 and added a new start codon to the beginning of the peptide region. CAHE_p0044 had no predicted signal peptide for the original protein on GenBank. We analyzed the DNA sequence upstream of the predicted CAHE_p0044 and found an in-frame start codon 27 amino acids upstream of the original start codon. With the addition of the 27 amino acids from the alternate CAHE_p0044 start codon, SignalP 6.0 predicted a signal peptide (lipoprotein signal peptide (Sec/SPII)) for the longer CAHE_p0044 (CAHE_p0044^+signal^). All five genes with signal peptides removed (or added) were subject to the same cloning and confirmation procedure as outlined above.

We tested for toxicity of all five CI candidate proteins using yeast spot plate assays, similar to previous studies (Beckmann et al., 2017). We grew transformed yeast strains containing expression plasmids with candidate genes overnight in DOB broth containing 2% glucose with SC lacking uracil and/or leucine at 30°C, 200 rpm. Cultures were pelleted, washed once with 1 mL of sterile water, and resuspended in 1 mL of sterile 1x PBS (Fisher Bioreagents). We diluted washed cultures to 0.2 OD600 with 1x PBS, which were then serially diluted in ten-fold steps to a final dilution of 10^-5^. Twenty microliters of each dilution step were then plated onto DOB agar containing 2% glucose (no induction) or 2% galactose (induction of gene of interest) lacking uracil (for single expression) or lacking both uracil and leucine (for co-expression) and incubated at 30°C until sufficiently grown (approximately 3 days with 2% glucose or 4-5 days with 2% galactose; coexpression strains were given approximately 2 extra days). We obtained images of plates using an Epson Perfection V850 Pro scanner.

We measured expression of *c*Eper1 CI candidate proteins from pelleted batch cultures of induced and uninduced yeast in triplicates (**Supplemental Data S6)**. Protein digests were prepared from cell pellets using a single-pot sample preparation method. Resulting peptide extracts were pelleted and supernatant was retained for measurement via LC-MS/MS and protein identification and quantification with a custom database. Full details on yeast proteomics are available in the **Supporting Information** and **Supplemental Data S7**.

### CI candidate protein localization in *S. cerevisiae*

*S. cerevisiae* was transformed with the low-copy pAG415 plasmid containing either CAHE_p0043^-signal^ or CAHE_0406, both of which lacked a stop codon. This created a C-terminal fusion of either protein with enhanced green fluorescent protein (eGFP), which was encoded by the plasmid vector backbone. Cultures were grown to stationary phase in DOB media containing 2% raffinose and lacking leucine for approximately two days at 30°C, 200 rpm. The stationary phase cultures were then pelleted and standardized to 0.3 OD600 in 6 mL of inducing DOB 2% galactose broth lacking leucine and grown for 2, 3, 4, 5, 6, 7, and 8 hours in separate biological replicates at 30°C and 200 rpm for induction. After incubation, 3 mL were pelleted and resuspended in 400 µL of 4% paraformaldehyde solution in 1x PBS and fixed for 20 minutes at room temperature in the dark. One milliliter of 1x PBS was added, then cells were pelleted and resuspended in 1 mL of fresh 1x PBS. This wash step was repeated for a total of two washes. Finally, the fixed cell pellet was resuspended in 500 µL of 1 µg/mL DAPI in PBS and incubated for 2 hours at room temperature in the dark. The DAPI-stained cells were concentrated by pelleting and resuspending in 50-60 µL of remaining supernatant. One to six microliters were then added to a field on a PTFE-printed microscope slide. After drying, a coverslip was applied with CitiFluor AF1 antifadent solution (Electron Microscopy Sciences) and sealed with clear nail polish. Slides were imaged on a Leica DMi8 inverted fluorescence microscope fitted with a Leica K5 camera. Images were taken and processed in the LAS X software using 1000x total magnification and fluorescence at the emission wavelengths of 525 nm for eGFP and 460 nm for DAPI. Brightfield images were also taken to visualize whole yeast cells. Background noise and blur was removed from images via Leica THUNDER technology using the large volume computational clearing method.

## Data Availability

Proteomic data, raw mass spectrometry files, and databases used are deposited in the ProteomeXchange Consortium via the PRIDE partner repository (Deutsch et al., 2023; Perez-Riverol et al., 2022) and will be provided upon request. See supplemental data S8-S11 for normalized data used in analyses and S12 for functional annotation of symbiont proteome.

## Author Contributions

Conceptualization, M.S.H., S.S., and M.K.; methodology, O.L.M., D.L.S., M.R.D., S.V., S.E.K., M.S.H., S.S., and M.K.; investigation, O.L.M., D.L.S., M.R.D., S.V., S.E.K.; formal analysis, O.L.M. and D.L.S.; writing - original draft, O.L.M. and D.L.S.; writing - review & editing, O.L.M., D.L.S., M.R.D., S.E.K., M.S.H., S.S., and M.K.; visualization - O.L.M., D.L.S., and M.R.D.; supervision M.S.H., S.S., and M.K.; funding acquisition, M.S.H., S.S., and M.K.

## Acknowledgments

This work was funded by the National Science Foundation awards #2426305 and #2003107 to M. Kleiner, #2002987 and #2426304 to S. Schmitz-Esser, and #2002934 and #2426306 to M.S. Hunter. We made all LC-MS/MS measurements in the Molecular Education, Technology, and Research Innovation Center (METRIC) at North Carolina State University, which is supported by the State of North Carolina, USA. We would also like to thank Cole Andersen for helping with yeast cultures and assays.

## Supplementary Materials

**Supporting Information**: Additional methods, results, supplementary figures, and discussion.

**Supplementary Data 1:** Summary statistics of LC-MS/MS run and database search.

**Supplementary Data 2:** Abundance of *c*Eper1 proteins across fractionated samples quantified by peptide spectral match counts normalized to Normalized Spectral Abundance Factors Percentages (NSAF%). Bars are colored according to fraction (P = light blue, PS = dark blue, S = gold).

**Supplementary Data 3:** Conversion of protein accessions (UniProt) to GenBank identifiers and locus tags.

**Supplementary Data 4:** Results from statistical analysis of fractionated sample metaproteomes.

**Supplementary Data 5:** Results from statistical analysis of tissue sample metaproteomes.

**Supplementary Data 6:** Sample name, construct, and media for yeast proteomic measurements.

**Supplementary Data 7:** Results from statistical analysis of yeast proteomic response to expression of CAHE_0406, CAHE_p0043, CAHE_0406^-signal^, CAHE_p0043^-signal^, and empty pYES plasmid under induction and noninduction conditions.

**Supplementary Data 8:** Normalized spectral abundance factors for the *c*Eper1 proteome from fractionated normalized to *c*Eper1 (orgNSAF).

**Supplementary Data 9:** Normalized spectral abundance factors (NSAF) for the full *c*Eper1-*Encarsia suzannae* metaproteome from fractionated samples.

**Supplementary Data 10:** Normalized spectral abundance factors for the *c*Eper1 proteome from tissue samples normalized to *c*Eper1 (orgNSAF).

**Supplementary Data 11:** Total sum scaled area under the curve abundance of yeast proteomes.

**Supplementary Data 12:** Functional annotation of *c*Eper1 proteome.

