## Supporting Information for "Identification of cytoplasmic incompatibility effectors of the reproductive manipulator *Cardinium hertigii*"

### Supplementary Methods

##### Whole wasp fractionated sample preparation

To identify symbiont proteins enriched in specific host sexes or life stages or secreted into the host, we separated symbionts from wasp tissue by differential centrifugation. For each sex and life stage combination we used four to five replicates (**Table S5, Supplementary Data 1**). For each replicate, we suspended 100 live wasps in 1x phosphate-buffered saline (PBS) with a protease inhibitor (1 tablet per 10 ml of PBS, Complete Mini EDTA-free Protease Inhibitor Cocktail Tablets, Roche 04693159001) and homogenized them to disrupt tissues using a glass homogenizer (Kimble Duall Tissue Grinders, DWK Life Sciences, 885460-0022). A differential centrifugation method was used to obtain three fractions (Hinzke et al., 2018). For the first fraction the disrupted samples were centrifuged at 1,000 x g for 2 minutes. The pellet was retained and will hereafter be referred to as the P fraction. The resulting supernatant was then centrifuged at 15,000 x g for 7 minutes for all sample types, except males which were centrifuged at 10,000 x g for 10 minutes. The resulting pellet and the supernatant were retained, and hereafter are referred to as the PS and S fractions. All samples were stored at -80°C until further processing. Prior to protein extraction, the S fraction was lyophilized overnight to concentrate. On the day of protein extraction, the P and S fractions were suspended in 120 μl of SDT lysis buffer (4% w/v sodium dodecyl sulfate, 100 mM Tris-HCl pH 7.6, 0.1 M dithiothreitol), while the PS fraction was suspended in 60 μl of SDT lysis buffer.

We confirmed symbiont enrichment in the fractions using Fluorescence *in situ* hybridization (FISH) following a previously described protocol (Daims et al., 2004) with *Cardinium* 16S rRNA specific Cy3 double-labeled probes (Ch1162 5’- (Cy3)- TTGACCTCATCCTCTCCT (Cy3-Q) - 3) (Doremus et al., 2020). Pellets from each fraction were resuspended in 1x PBS, then air dried on coverslips. Samples were then fixed with 4% paraformaldehyde for 20 min at room temperature. Fixative was removed and samples were washed three times with ddH2O. Samples were then dehydrated using a series of 3 min rinses in 50, 80, and 96% ethanol. Dehydrated samples were incubated with 20 μl hybridization buffer (10% formamide, 0.9 M NaCl, 20 mM Tris HCL (pH 8), 0.01% SDS) and 2 μl cy3 probes (30 ng/μl) in the dark at 46°C for ~ 2 hours. Samples were submerged in wash buffer (0.45 M NaCl, 20 mM Tris HCl (pH 8)) for 10-15 min in a 48°C water bath. Samples were then rinsed in ice cold ddH2O for 1-2 seconds and dried with compressed air. Host nuclei and symbiont DNA were stained with 10 μl DAPI (10 μg/mL) for 5 min in the dark, followed by three rinses with ddH2O. Coverslips were mounted to slides with mounting media (80% glycerol, 20% TBST with 2% *n*-propyl-gallate (Sigma)) and stored in the dark at 4°C until visualization. Images were collected using a Zeiss LSM 880 inverted confocal microscope using a 63X lens with oil immersion. DAPI and cy3 fluorescence were captured using a 405 and 561 nm excitation beam, respectively. Final images were processed using the Zen Blue Software (Zeiss).

##### Tissue sample preparation

To obtain tissue samples, wasps were dissected in ~40 μl of 1X PBS buffer. Dissections included removing the head (to serve as a control against the reproductive tissues) and the reproductive organs (testes+seminal vesicle or/ovaries). Tissues were suspended in 4% w/v sodium dodecyl sulfate, 100 mM Tris-HCl pH 7.6 either manually with very fine “minuten” pins or pipetted with a 20 μl pipette tip coated in Tween 20. Approximately 30-50 tissue samples were collected in a batch, then stored at -20°C until used for protein extraction. Tissue dissections were thawed and pooled to obtain four to five replicates of each tissue sample with 50-75 tissues in each. Heads were homogenized prior to cell lysis to disrupt the insect cuticle, while testis, ovary, and sperm samples dissolved in the lysis buffer without disruption.

**Filter aided sample preparation for wasp proteomics**

All samples were heated to 95°C for 10 minutes to lyse the cells. A peptide mixture was obtained from each sample using a modified filter-aided sample preparation (Wiśniewski, 2019). Briefly, the full sample lysate was loaded onto 10 kDa VWR modified PES centrifugal filters with UA buffer (8 M urea in 0.1 M Tris/HCl, pH 8.5) using a ratio of 60 μl of sample lysate to 400 μl of UA, then centrifuged at 14,000 x g for 20 minutes (or until there was no standing liquid remaining on the filter). This step was repeated for the P- and S-fractions to load the full lysate amount. All filters were then washed with an additional 200 μl of UA and centrifuged as before, before treating the samples with 100 μl of IAA (0.05 M iodoacetamide in UA buffer). For this treatment, samples were mixed for 1 minute at 600 rpm in a heating and cooling shaker (Benchmark Scientific Inc.) and incubated at room temperature for 20 minutes without mixing. Samples were then centrifuged as before. The samples were subjected to a series of washing steps: 100 μl of UA centrifuged as before then repeated twice and 100 μl of ABC (50 mM ammonium bicarbonate) centrifuged as before then repeated twice. Filters were transferred to clean collection tubes. Trypsin (0.5-1.0 μg in 40 μl of ABC per sample) was added to the proteins retained on the filter units and mixed for 1 minute at 600 rpm in a thermo-mixer, before overnight digestion at 37°C in a wet chamber. After digestion, samples were centrifuged as before. Lastly, 50 μl 0.5 M NaCl was added and mixed for 1 minute at 600 rpm in a thermo-mixer. Filter units were centrifuged as before to elute the peptides off the filters. This final eluate was retained for further analysis.

##### LC-MS/MS measurement of wasp proteome

Peptide eluate concentrations were determined using the Pierce MicroBCA assay (Thermo Fisher Scientific). For each sample, 600 ng of peptide mixture was loaded at flow rate of 5 μl min^-1^ using an UltiMate 3000 RSLCnano liquid chromatograph (Thermo Fisher Scientific) with loading solvent A (2% acetonitrile, 0.05% trifluoroacetic acid) onto a nano trap cartridge (300 μm i.d. x 5 mm) packed with C18 Acclaim PepMap100 (Thermo Fisher Scientific). After loading, the nano trap cartridge was put in line with a 75 μm, 75 cm EASY-Spray analytical column packed with PepMap RSLC C18 (Thermo Fisher Scientific) heated to 60°C. The peptides were separated using a 140 minute gradient at a flow rate of 300 nL min^-1^ previously implemented (Mordant and Kleiner, 2021). Briefly, the gradient began at 95% eluent A (0.1% formic acid) and ended at 99% eluent B (0.1% formic acid, 80% acetonitrile. A wash step was performed after each sample run by injecting 20 μl of acetonitrile with 99% eluent B to decrease carryover between samples. For fractionated whole wasp samples, an Easy-Spray source connected the analytical column to a Q Exactive HF Hybrid Quadrupole-Orbitrap mass spectrometer (Thermo Fisher Scientific) which was used to measure eluted peptides producing MS^1^ and MS^2^ spectra acquired using a previously published method (Mordant and Kleiner, 2021). In brief, MS1 precursor scans were acquired with a scan range of 380 to 1,600 m/z at 60,000 resolution with a 200 ms maximum injection time (IT) and 3e6 AGC target. The top 15 most abundant precursor ions were selected using a 25 s dynamic exclusion duration and fragmented using 24% normalized collision energy. MS2 scans were acquired at 15,000 resolution with a 100 ms maximum IT and 5e4 AGC target. Tissue specific samples were measured using an Orbitrap Exploris 480 mass spectrometer (Thermo Fisher Scientific) using a previously published method (Violette et al., 2024). Here, MS1 precursor scans were acquired and precursor ions were selected for fragmentation following the method detailed above. Selected precursor ions were subjected to a 27% normalized collision energy applied in the HCD cell. MS2 spectra were acquired at 15,000 resolution with a maximum IT of 50 ms and an AGC target of 1e5. Full details of the methods used for the LC-MS/MS runs can be found in **Tables S6** and **S7**.

##### Protein identification

To identify peptides from the spectral data, we used the database search engine SEQUEST HT in Proteome Discoverer 2.3 (Thermo Scientific). We quantified proteins using peptide spectral matches (PSMs). We constructed a protein sequence database, using previously outlined recommendations (Blakeley-Ruiz and Kleiner, 2022). Our database included protein sequences predicted from a preliminary assembly of the adult and pupal male and female *E. suzannae* transcriptome (Schultz et al., 2022) and *Cardinium c*Eper1 proteome (<https://www.uniprot.org/proteomes/UP000003996>) (Penz et al., 2012). We also included possible contaminant protein sequences specific to the system including the proteomes of the host of *E. suzannae*, *B. tabaci* MEAM1 (<https://www.ncbi.nlm.nih.gov/datasets/genome/GCF_001854935.1/>), and its symbionts *Hamiltonella defensa* (<https://www.uniprot.org/proteomes/UP000216438>), *Portiera aleyrodidarum* (<https://www.uniprot.org/proteomes/UP000215611>), and *Rickettsia sp.* MEAM1 (<https://www.uniprot.org/proteomes/UP000215668>). To construct the insect portion of the database, male and female adult *E. suzannae* transcriptome-derived protein sequences were merged and clustered at 95% similarity to remove redundancy using CD-HIT (Version 4.7) (Fu et al., 2012; Li and Godzik, 2006). This reduced the number of sequences from 179,209 to 142,008. The pupal transcriptome was clustered at 95% similarity using CD-HIT (no protein sequences clustered), and then compared to the adult proteome to identify unique sequences using CD-HIT-2D at 90% similarity resulting in 92,329 unique protein sequences. The unique pupal sequences were then merged with the adult sequences and clustered at 95% similarity, resulting in 229,024 *E. suzannae* sequences in total. Similarly, the *B. tabaci* proteome was compared to the *E. suzannae* proteome to identify unique sequences at a 90% similarity. These unique sequences were merged with *E. suzannae* protein sequences and clustered at 95% similarity. This resulted in an insect protein database of 242,183 sequences. The reference proteomes of *C. hertigii c*Eper1, *H. defensa*, *P. aleyrodidarum*, and *Rickettsia sp.* MEAM1 were merged with the insect portion of the database without clustering. Common laboratory contaminants were accounted for in the database by including sequences from the cRAP database (<https://www.thegpm.org/crap/>). The resulting database had a total of 245,985 protein sequences (98.45% insect sequences, 0.36% *C. hertigii* *c*Eper1 sequences). PD results were exported as text files, merged and filtered to obtain a matrix of peptide spectral match (PSM) counts for master proteins identified at a 5% false discovery rate (FDR). Functional protein annotations were curated using results from BLASTKoala (Kanehisa et al., 2016), SignalP 5.0 (Almagro Armenteros et al., 2019b), TargetP 2.0 (Almagro Armenteros et al., 2019a), TMHMM 2.0 (Krogh et al., 2001), InterProScan (Blum et al., 2025; Jones et al., 2014; Paysan-Lafosse et al., 2025), MANTIS (Queirós et al., 2021), and DeepLoc 2.0/2.1 (Ødum et al., 2024; Thumuluri et al., 2022) (**Supplementary Data 12**).

##### Yeast expression strain preparation

We selected five candidates from our proteomic analyses for heterologous expression in *Saccharomyces cerevisiae* based on a combination of expression data, curated protein annotations, previous comparative genomics work (Schultz et al., 2025), and expected characteristics based on current cytoplasmic incompatibility literature. Evidence supporting candidate selection can be found in **Table 1**.

*Cardinium* *c*Eper1 CI candidate genes CAHE_0757, CAHE_0405, CAHE_0406, CAHE_p0043, and CAHE_p0044, along with the *Wolbachia* wPip *cidB* CI gene (WP0283 / CAQ54391.1) were ordered for synthesis through ThermoFisher GeneArt gene synthesis (**Table S11**) and subcloned into the Gateway entry vector pDONR221 (Katzen, 2007). Invitrogen Gateway LR Clonase II was used to swap the synthesized genes from their entry vector into the desired expression plasmids shown in **Table S10** (Alberti et al., 2007). The manufacturer’s protocol was used with an extended incubation period (2-5 hours). The LR reaction products were then immediately transformed into chemically competent ccdB-sensitive StrataClone *Escherichia coli* (Agilent) via heat shock following the manufacturer’s protocol and plated onto LB media containing 100 µg/mL ampicillin for selection. Colonies were picked from the initial transformation and restruck onto LB + 100 µg/mL ampicillin for maintenance and onto LB + 30 µg/mL chloramphenicol to confirm the presence of the inserted gene of interest. Colonies which did not grow on LB agar containing chloramphenicol were screened for the correct size insert via PCR and 1% agarose gel electrophoresis using primers targeting the GAL1 promoter (5’ AATATACCTCTATACTTTAACGTC 3’) and CYC1 terminator (5’ GCGTGAATGTAAGCGTGAC 3’) present on the yeast plasmids. Some transformants were also selected for additional confirmation via Sanger sequencing. Transformants with the correct inserts were subject to plasmid miniprep using the Invitrogen Quick Plasmid Miniprep kit.

Miniprepped plasmids were then transformed into *S. cerevisiae* strain BY4741 (MATa, his3Δ1, leu2Δ0, met15Δ0, ura3Δ0) using lithium acetate pretreatment and electroporation (Thompson et al., 1998). Briefly, an 8 mL overnight culture of *S. cerevisiae* BY4741 was grown at 200 rpm, pelleted, then washed twice with 1 mL of ice-cold sterile water. The pellet was then suspended and incubated at 30°C for 30 minutes in 2 mL of LiAc/DTT/TE solution containing 0.1 M lithium acetate, 10 mM dithiothreitol, 10 mM Tris–HCl (pH 7.5), and 1 mM EDTA. Following incubation, cells were kept on ice, pelleted, and washed twice with ice-cold sterile water before being resuspended in 1 mL then 200 µL of ice-cold 1 M sorbitol. Five hundred to 1 µg of plasmid DNA was added to the treated cells, and the solution was incubated on ice for 5 minutes before electroporation in a 2 mm electroporation cuvette at 1.5 kv, 25 µFd, and 200 Ω using a Bio-Rad Gene Pulser Xcell electroporation system. One milliliter of ice-cold 1 M sorbitol was added immediately after electroporation, and cuvette contents were plated on drop-out base (DOB) medium containing 2% glucose (MP Biomedicals) supplemented with synthetic complete (SC) mixture (MP Biomedicals) lacking uracil (for strains transformed with pYES) and/or leucine (for strains transformed with pAG415). Colonies were picked and screened for the correct size insert via PCR as outlined above. Yeast was grown at 30°C unless otherwise noted.

We generated yeast coexpression strains by sequential transformation: one plasmid was transformed into *S. cerevisiae* BY4741 via the above method, plated onto DOB media lacking either leucine or uracil, then confirmed via PCR. Next, we transformed the second plasmid into the yeast strain containing one expression plasmid and plated onto media lacking both leucine and uracil. The presence of both plasmids was confirmed via PCR and growth on media lacking leucine and uracil. For more info on all yeast constructs used for this study, see **Table S11**.

CI candidate genes CAHE_0757, CAHE_0405, CAHE_0406, and CAHE_p0043 were all predicted to encode lipoprotein signal peptides (Sec/SPII) via SignalP 6.0 in “slow” mode (Teufel et al., 2022). Although we do not know the fate of each of these CI candidate proteins and whether they are processed by signal peptide cleavage and exported, we wanted to make sure to also test these proteins as they are predicted to be post signal peptide cleavage. Therefore, we generated additional constructs of these four genes by removing the predicted signal peptide up to the predicted cut site given by SignalP and adding a new start codon to the beginning of each gene. In the case of CAHE_p0044, there was no predicted signal peptide for the original protein on GenBank, but an analysis of the plasmid DNA sequence revealed an in-frame start codon 27 amino acids upstream of the predicted CAHE_p0044 start codon, suggesting the GenBank predicted CAHE_p0044 start codon may be incorrect. With the addition of the 27 amino acids from the alternate CAHE_p0044 start codon, SignalP predicted the same type of signal peptide for the longer CAHE_p0044 (named p0044^-signal^) as was predicted for the other four CI candidate proteins (lipoprotein signal peptide (Sec/SPII)). All five genes with signal peptides removed (or added) were subject to the same cloning and confirmation procedure as outlined above. All were then tested for toxicity in yeast along with their unaltered counterparts.

##### Yeast heterologous expression assays

Yeast spot plate assays to test for toxicity of the CI candidate proteins were performed similar to previous studies (Beckmann et al., 2017). Strains containing expression plasmids with genes of interest were grown overnight in DOB broth containing 2% glucose with SC lacking uracil and/or leucine at 30°C, 200 rpm. Cultures were pelleted, washed once with 1 mL of sterile water, and resuspended in 1 mL of sterile 1x PBS. Washed cultures were diluted to 0.2 OD600 with 1x PBS then serially diluted 1:10 to a final dilution of 10^-5^. Twenty microliters of each dilution step were then plated onto DOB agar containing 2% glucose (no induction) or 2% galactose (induction of gene of interest) lacking uracil (for single expression) or lacking both uracil and leucine (for coexpression) and incubated at 30°C until sufficiently grown (approximately 3 days with 2% glucose or 4-5 days with 2% galactose; coexpression strains were given approximately 2 extra days). Plates were then imaged on an Epson Perfection V850 Pro scanner.

##### Proteomics Analysis on Yeast Cultures

We prepared proteomic samples from cultures of transform *Saccharomyces cerevisiae* to verify expression of our CI candidate constructs. Uninduced yeast cultures were grown at 30°C for 14-24 hours at 200 rpm in DOB 2% glucose broth with synthetic complete mixture (SC) lacking uracil (single expression) or DOB 2% glucose broth with SC lacking uracil and leucine (co-expression). For induction of CI candidate genes, cultures were grown in DOB 2% galactose media with SC -uracil or -uracil - leucine at 30°C for 24-48 hours at 200 rpm. Due to toxicity of some candidates when expressed in yeast, were not able to standardize induction times across constructs and needed to adapt the procedure by initially growing toxic candidate protein expressing yeast on DOB 2% raffinose media with SC -uracil or -uracil - leucine for 24-30 hours before resuspending in DOB 2% galactose media with SC and allowing for induction for 14-17 hours prior to collection. Using this procedure, we prepared 3 replicates of our 17 constructs uninduced and induced. We pelleted 1.5-4.5 ml of each culture, removing the media prior to storage at -80°C.

For proteomics, we thawed the samples on ice and then transferred 0.5-1 µl of the yeast pellets to 40 µl of HPLC-grade water with 5 mM DTT using sterile pipette tips. We froze samples at -80°C for 1 hour before heating samples to 90°C for 30 minutes. We repeated this freeze-thaw cycle (total of two freeze-thaw cycles) and then cooled samples to 12°C. After cooling, we briefly spun down the samples with a benchtop centrifuge and sonicated the samples in a water bath for 5 minutes. After sonication, samples were briefly vortexed and spun down. Then, we added 2 µl of 300 mM iodoacetamide (final concentration of 15 mM in sample) and incubated samples in the dark for 30 minutes. We prepared samples using a low-input, single-pot sample preparation method adapted from Petelski et al. (2021). We added 8.8 µl of a master mix consisting of 100 ng/µl Trypsin Gold (Promega) reconstituted in 50 mM glacial acetic acid, 500 mM TEAB, and 1.08 U benzonase nuclease prepared in HPLC water to each sample. We then incubated samples in a thermocycler held at 37°C for 3 hours (lid heated to 52°C). After, we added 13.3 µl of 0.5% hydroxylamine, vortexed briefly, spun down, and incubated at room temperature for 30 minutes. We then centrifuged samples at 10,000 x g for 5 minutes to pellet any particulate debris before transferring supernatants to fresh HPLC vials for 1D-LC-MS/MS.

We loaded 2 µl of each sample onto a PepMap Neo trap cartridge (Thermo Scientific, cat#: 174500) using a nano-flow Vanquish Neo UHPLC system (Thermo Scientific). We separated sample peptides on an Easy-Spray PepMap Neo column (Thermo Scientific, cat#: ES75750PN) using a 70 minute reverse-phase gradient at a 0.250 µl/min flow rate as follows: 95% eluent A (0.1% formic acid) to 28% eluent B (80% acetonitrile, 0.1% formic acid) over 39 minutes, 28% to 40% eluent B over 14 minutes, 40% to 99% eluent B over 2 minutes, and then held at 99% eluent B for 15 minutes. We included a 15 minute oscillating (eluent A to eluent B) wash injection between samples. Eluting peptides were ionized by electrospray ionization in an Easy-Spray source prior to measurement with an Orbitrap Eclipse mass spectrometer (Thermo Scientific) using a data dependent acquisition method. We acquired precursor scans in the Orbitrap mass analyzer with a scan range of 380-1,600 m/z at a resolution of 60,000, a 200 ms maximum injection time, and a 300% (1.2e6 ions) AGC target. The top 15 ions from each precursor scan with a dynamic exclusion of 25 s were selected for fragmentation and subjected to 27% normalized collision energy in the HCD cell. We then acquired MS2 scans of fragmented peptides in the Orbitrap mass analyzer at a resolution of 15,000, a 50 ms maximum injection time, and 200% (1.0e5 ions) AGC target. Full details of the methods used for the LC-MS/MS runs can be found in **Table S8**.

We built a custom proteomics database using available UniProt/Swiss-Prot protein sequences of *S. cerevisiae* from NCBI (TaxID: 4932), all available protein sequences from *S. cerevisiae* BY4741 from NCBI (TaxID: 1247190), and the 5 candidate proteins from *c*Eper1. We also included common laboratory contaminants (reduced cRAP database, <http://www.thegpm.org/crap/>). We searched raw spectral data against this database using Proteome Discoverer 2.3 (Thermo Scientific). We used the Sequest HT node with the following settings: Trypsin (Full) allowing for a maximum of 2 missed cleavages, peptide length of 6 to 144 amino acids, 10 ppm precursor mass tolerance, 0.1 Da fragment mass tolerance, and a maximum of 3 equal and 4 dynamic modifications allowed per peptide. The dynamic modifications included in our method were oxidation (+15.995 Da [M]), deamidation (+0.984 Da [N, Q, R]), and N-terminus acetyl group (+42.011 Da). We also included a static carbamidomethylation (+57.021 Da [C]) modification. We calculated peptide FDR confidence using the Percolator node, retaining only PSMs with FDR < 0.05. We used the Protein FDR Validator node for protein inference and classified proteins as high confidence (FDR < 0.01) or medium confidence (0.0 1< FDR < 0.05). The proteins were quantified using an area under the curve (AUC) method with the Minora Feature Detector node (minimum chromatographic trace length of 5 non-zero points and high PSM confidence). In the consensus step, we used the Feature Mapper node (Perform RT Alignment = FALSE) and the Precursor Ions Quantifier Node (use unique + razor peptides, considering protein groups for uniqueness, and calculate protein abundance based on summed abundances). We exported results from Proteome Discoverer and compiled into a single table with a custom python script before processing in R (version 4.2.2) with stringr (version 1.6.0), tidyr (verison 1.3.1), and dplyr (version 1.1.4). The dataset was filtered to remove contaminant proteins and transformed via total sum scaling. Because expression of *c*Eper1 constructs was captured through presence-absence patterns in all instances, we deemed statistical assessment of construct expression unnecessary. To assess global differential expression patterns associated with toxic constructs (CAHE_0406 and CAHE_p0043^-signal^) for significance, we subset data by construct, replaced missing values with half of the lowest value in each dataset, and log2 transformed the data prior to testing Welch’s t-test with Benjamini-Hochberg FDR correction. A major confounding variable in our comparisons is the medium. To account for this, we repeated the same analyses on the yeast proteome with the empty pYES construct and non-toxic versions of the constructs (CAHE_0406^-signal^ and CAHE_p0043) to assess impact of induction medium. Comparisons were visualized using the following packages: ggplot2 (version 4.0.2), ggbeeswarm (version 0.7.3), and ggpubr (version 0.6.3).

### Supplementary Results

##### Coexpression of CI candidate proteins to test for rescue

We attempted to screen for rescue of CAHE_0406 toxicity in yeast similar to previous studies with *Wolbachia* which found that expression of a toxic CifB with its cognate CifA led to a restoration of yeast growth, mimicking the induction and rescue of natural CI (Beckmann et al., 2019, 2017; Murphy and Beckmann, 2024; Sun et al., 2022; Xiao et al., 2021). To accomplish this, we utilized the low-copy Gateway-compatible plasmid pAG415GAL-ccdb-eGFP (Addgene plasmid # 14193) with subcloned CAHE_0406 (Alberti et al., 2007). However, we saw no toxicity caused by CAHE_0406 to yeast when expressed from this low-copy plasmid. To account for this loss of toxicity, we generated coexpression constructs with the toxic CAHE_0406 expressed from the same high-copy pYES-DEST52 plasmid as previously along with a potential antidote (CAHE_0405) expressed from the low-copy pAG415 plasmid. However, we noted no toxicity in our positive control consisting of expressing CAHE_0406 from pYES-DEST52 in *S. cerevisiae* which was also transformed with an empty pAG415 plasmid, despite detection of the protein in the proteome (**Figure S7**). Therefore, we could not test rescue factors for CAHE_0406 using the two-plasmid system in *S. cerevisiae* since CAHE_0406 is not toxic to yeast when expressed from a low-copy plasmid or when expressed in co-expression with a secondary plasmid.

We also investigated rescue of CAHE_p0043^-signal^ toxicity by coexpressing this CI candidate protein from the low copy pAG415 vector along with both versions of CAHE_p0044 (with and without added signal peptide) expressed from the high-copy pYES-DEST52. However, we observed complete lethality in all coexpression constructs containing CAHE_p0043^-signal^, suggesting that CAHE_p0044 may not be a potential rescue factor for CAHE_p0043-mediated toxicity (**Figure S7**). However, this result remains inconclusive given the lack of detection of CAHE_p0043 in the co-expression assay (**Figure S7**).

### Supplementary Discussion

##### Yeast coexpression highlights issues with elucidating mechanisms of rescue

Because of the models proposed in Doremus *et al*., and CI research at large, we were expecting the CI phenotype to be rescued by a neighboring protein (Doremus et al., 2020; Wang et al., 2022). The *Wolbachia cifs* are encoded adjacent to each other and were found to act like a standard toxin/antitoxin system when coexpressed in *S. cerevisiae:* CifB was toxic to yeast when expressed from a low-copy plasmid unless it was coexpressed with the correct cognate CifA on a high-copy plasmid (Beckmann et al., 2017; Murphy and Beckmann, 2024; Sun et al., 2022). Both of the top *c*Eper1 CI candidate proteins have an adjacent gene, CAHE_0405 and CAHE_p0044, with similar expression patterns to their toxic counterparts, and therefore were our main candidates for rescue of their cognate toxic candidates.

Our observation that CAHE_0406 expressed from a high-copy plasmid was not toxic when the low-copy pAG415 plasmid was also present (despite being toxic when singly expressed from the high-copy plasmid) highlights a general challenge with our CI candidate genes and the yeast system. We hypothesize that the toxicity of CAHE_0406 may be dose-dependent. Therefore, its toxicity may be negated due to a reduced protein load resulting from its expression on a low-copy plasmid or from coexpression with another gene on a different plasmid under control of a competing GAL1 promoter. It is unclear if CAHE_0406 forms a multimeric protein, which may explain why protein dose is important in its toxicity, or if other factors are involved, such as an inability of the small CAHE_0406 protein to localize correctly on its own, meaning that high dosages are needed to ensure some of the protein is present in the correct subcellular area of action. A previous study has highlighted the importance of high transcription to the induction of CI, finding that the differences in CI strength across various *Drosophila*-*Wolbachia* systems are largely explained by the levels of *cifB* transcripts, with higher levels of *cif* transcripts usually found in hosts displaying stronger CI (Shropshire et al., 2022). It is possible that CAHE_0405 does rescue CAHE_0406-mediated toxicity, but additional work is needed to elucidate the relationship between these two potentially CI-relevant proteins.

We hypothesized that coexpression of CAHE_p0043^-signal^ and a form of CAHE_p0044 (with or without signal peptide) may result in rescue similar to CifB/CifA coexpression given their proximity and the annotation of CAHE_p0044. Although the predicted SMI1/KNR4 domain in CAHE_p0044 was originally characterized as part of a protein important for fungal cell wall stress tolerance (Kroll et al., 2025; Martin-Yken et al., 2016), it has also been suggested to be involved in detoxifying nucleic acid degrading toxins (Zhang et al., 2011). Further, CAHE_p0044^-signal^ was predicted to localize to the eukaryotic nucleus (0.828), which would be expected for a CI rescue factor. However, no rescue of CAHE_p0043^-signal^ toxicity was observed for either version of CAHE_p0044. It is possible that dosage effects may play a role in our lack of rescue of CAHE_p0043 toxicity: perhaps the amount of CAHE_p0043^-signal^ produced in coexpression was more than could be nullified by the amount of CAHE_p0044 available, even with the toxin expressed from a low-copy plasmid. Further, since we did not test a positive control of rescue of *Wolbachia w*Pip CidB via coexpression, we cannot be certain that rescue is possible in our yeast system. We also observed much greater toxicity of *w*Pip CidB at lower temperatures than in previous studies (Beckmann et al., 2019, 2017), so it is possible that our yeast expression system is more susceptible to toxic effects than those from prior studies. As a result, we cannot confirm if CAHE_p0044 is a rescue factor for CAHE_p0043^-signal^, but the possibility cannot be entirely ruled out. It is also possible that, if CAHE_0406 or CAHE_p0043 cause CI, rescue of the phenotype may be due to proteins other than CAHE_0405 and CAHE_p0044, or perhaps via an entirely different unknown mechanism. Additional research is needed to explore rescue of these CI candidate proteins.

##### Additional proteins with potential CI involvement identified from metaproteomics data

CAHE_0669, a predicted peptidyl-prolyl cis-trans isomerase, was uniquely detected in the male reproductive organs. Peptidyl-prolyl cis-trans isomerases have been shown to play a role in the maturation of secreted proteases (Lyon and Caparon, 2003) and nucleases (Wiemels et al., 2016), as well as intracellular pathogen virulence factor secretion and pathogenesis (Alonzo et al., 2009; Alonzo and Freitag, 2010). The tissue-specific detection of this protein suggests possible involvement in activating CI factors or promoting the intracellular lifestyle of *Cardinium* in the male reproductive organs. Another protein of interest is CAHE_0435, an ankyrin repeat protein that was enriched in pupal testes. Bacterial ankyrin repeat domains are indicative of protein-protein interaction and have been implicated in diverse functions (Al-Khodor et al., 2010), such as CI and other reproductive manipulation phenotypes in conjunction with other predicted domains (Amoros et al., 2025; Hochstrasser, 2023; Pollmann et al., 2022; Siozios et al., 2013). CAHE_0465, which was detected and enriched in the male secretome (**Supplementary Data 4**), is a probable protease with a PDZ domain. This protein also matches to the pro-apoptotic serine protease NMA111 family (PANTHER: PTHR46366), which is a protein family responsible for downregulating apoptosis inhibitors and increasing apoptotic phenotypes when upregulated in *Candida albicans* (Nam et al., 2022). This protein also has a predicted signal peptide, and, if secreted by the symbiont, could interact with apoptotic pathways in its host cell as a toxin (Kaushal et al., 2023). CAHE_0706 and CAHE_0705 are another possible CI pair, similar to the paired genomic localization of *Wolbachia cif* genes and the *Cardinium* candidates proposed in this study, CAHE_p0043/p0044 and CAHE_0405/0406 pairs. CAHE_0706, a 336 amino acid predicted collagen triple helix repeat-containing protein, was detected in the secretome with a secreted pattern. Bacterial collagens have been suggested to be involved in host-microbe interaction, potentially enhancing invasion and evasion of the immune system (Yu et al., 2014). CAHE_0705 is an 80 amino acid lipoprotein of unknown function that was among the 20 most abundant symbiont proteins in the pupal testes, and was only detected in the pupal testes and adult ovaries, suggesting tissue-specific tropism (**Figure 1D, S3**).

We also observed type VI secretion system (T6SS) components among the most abundant proteins in nearly all sample types (**Figure S2**). This is consistent with previous transcriptomic evidence from the *c*Eper1 strain (Mann et al., 2017). *Cardinium* and its closest relative, *Ameobophilus asiaticus*, have a unique arsenal-like T6SS (subtype iv) with several T6SS aligning on the cell wall to make a structure resembling stacked lines of filaments (Böck et al., 2017; Mathieson et al., 2025; Penz et al., 2012). The targets of this secretion system and its effectors are not known; however, given the prominence of this complex feature in the proteome it likely plays an integral role in symbiont function and lifestyle, such as host interaction or microbe-on-microbe competition (Chen et al., 2019; Wood et al., 2020). Interestingly, we observed sex-specific enrichment of T6SS of individual components which might suggest a role in CI induction (**Figure S2**), though this will need to be confirmed with more absolute quantification measures before speculating on host-sex specific roles.

### Supplementary Tables

**Table S1. Genomes of *Cardinium* and sister taxa (*Cand.* Amoebophilus asiaticus 5a2) used in this study.** Known phenotype: cytoplasmic incompatibility (CI) and parthenogenesis (PI).

| **Cardinium strain (phenotype)** | **GenBank genome accession** | **Host taxa** |
| --- | --- | --- |
| **Insects** | | |
| **Hymenoptera** | | |
| ***c*Eper1 (CI)** | **GCA_000304455.1** | ***Encarsia suzannae*** |
| *c*Eper2 (PI) | GCA_050864935.1 | *Encarsia tabacivora* |
| *c*Ehis1 (PI) | GCA_050864915.1 | *Encarsia hispida* |
| *cEina2* | GCA_050864895.1 | *Encarsia partenopea* |
| ***c*Eina3 (CI)** | **GCA_050864855.1** | ***Encarsia partenopea*** |
| **Hemiptera** | | |
| CanCar | GCA_004300865.1 | *Bemisia tabaci* |
| *c*BtQ1 | GCA_000689375.1 | *Bemisia tabaci* |
| ***c*Sfur (CI)** | **GCA_003351905.1** | ***Sogatella furcifera*** |
| ihNabLimb1 | GCA_965196625.1 | *Nabis limbatus* |
| **Diptera** | | |
| *c*Cpun | GCA_004354815.1 | *Culicoides punctatus* |
| idTipUnca1 | GCA_964020025.1 | *Tipula unca* |
| **Coleoptera** | | |
| icPhiSpin1 | GCA_964030745.1 | *Philonthus spinipes* |
| **Arachnids** | | |
| **Araneae** | | |
| *c*Oegib-Wal | GCA_936981045.1 | *Oedothorax gibbosus* |
| **Trombidiformes** | | |
| *c*BcalN1 | GCA_022810505.1 | *Brevipalpus californicus* |
| *c*BcalN2 | GCA_022810445.1 | *Brevipalpus californicus* |
| *c*ByotB1 | GCA_022763005.1 | *Brevipalpus yothersi* |
| *c*ByotN1 | GCA_022810525.1 | *Brevipalpus yothersi* |
| **Sarcoptiformes** | | |
| *c*Dfar | GCA_007559345.1 | *Dermatophagoides farinae* |
| DF | GCA_025268635.1 | *Dermatophagoides farinae* |
| TP | GCA_025215055.1 | *Tyrophagus putrescentiae* |
| *c*Tput | GCA_030441515.1 | *Tyrophagus putrescentiae* |
| JH06282024_1 | GCA_036541165.1 | *Tyrophagus putrescentiae* |
| JH06282024_2 | GCA_036541185.1 | *Tyrophagus putrescentiae* |
| **Nematodes** | | |
| **Tylenchida** | | |
| *c*Ppe | GCA_003788695.1 | *Pratylenchus penetrans* |
| *c*HgTN10 | GCA_003176915.1 | *Heterodera glycines* |
| *c*Hhum | GCA_028766965.1 | *Heterodera humuli* |
| **Amoebozoa (sister taxa)** | | |
| *Amoebophilus asiaticus* 5a2 | GCA_000020565.1 | *Acanthamoeba spp.* |

**Table S2. CAHE_0406 and CAHE_p0043 annotations based on results of BLASTp against the UniProtKB+SwissProt protein database.** The top 5 matches are shown for each protein. Also included for CAHE_0406 are the top 3 matches that are reviewed SwissProt entries (indicated by *****). CAHE_p0043 did not have matching proteins from SwissProt.

| **Protein Name** | **UniProt Entry** | **Length (AA)** | **Protein Families** | **Organism** | **Identity**  **(%)** | **Score** | **E-value** |
| --- | --- | --- | --- | --- | --- | --- | --- |
| **CAHE_0406** (query) | **K0P5U4 (UPI00027EA19A)** | **60** |  | ***c*Eper1** |  |  |  |
| Chaperone protein ClpB | A0A292YGM0 | 870 | ClpA/ClpB family | *Lebetimonas natsushimae* | 72.7 | 88 | 0.06 |
| DUF7918 domain-containing protein | A0A6A7AXD3 | 1,051 |  | *Plenodomus tracheiphilus* IPT5 | 35.7 | 85 | 0.16 |
| Uncharacterized protein | A0A834I932 | 203 |  | *Rhynchophorus ferugineus* | 56.4 | 84 | 0.2 |
| Uncharacterized protein | A0ABM5HJ22 | 834 | MAP7 family | *Drosophila rhopaloa* | 46.9 | 84 | 0.22 |
| Uncharacterized protein | A0A1M5XPD9 | 283 |  | *Clostridium collagenovorans* DSM 3089 | 56.2 | 83 | 0.29 |
| Omega-theraphotoxin Hhn1f 3***** | D2Y2E9 | 86 | Neurotoxin 10 (Hwtx-1) family, 17 (Hntx-9) subfamily | *Cyriopagopus hainanus* | 38.2 | 76 | 2.1 |
| Nucleoid-associated protein CTN_1899***** | B9KAU2 | 118 | YbaB/EbfC family | *Thermotoga neapolitana* (strain ATCC 49049/DSM 4359/NBRC 107923/NS-E) | 51.6% | 74 | 5.3 |
| Omega-theraphotoxin Hs2a***** | P61104 | 86 | Neurotoxin 10 (Hwtx-1) family, 17 (Hntx-9) subfamily | *Cyriopagopus schmidti* | 38.2 | 73 | 6.1 |
| **CAHE_p0043 (query)** | **K0P6Z1 (UPI00027E9E2E)** | **312** |  | ***c*Eper1** |  |  |  |
| ATP-dependent Zn protease | A0ABQ0J2M3 | 701 | AAA ATPase family | ‘*Chrysanthemum coronarium*’ phytoplasma | 23.1 | 112 | 0.02 |
| C3H1-type domain-containing protein | A0A1J1H6Q0 | 2,254 |  | *Plasmodium relictum* | 24.6 | 108 | 0.09 |
| Uncharacterized protein | W7K1G7 | 1,845 |  | *Plasmodium falciparum* (isolate NF54) | 22.8 | 106 | 0.15 |
| Protein-tyrosine-phosphatase | A0A1I2K578 | 243 | Metallo-dependent hydrolases superfamily, CpsB/CapC family | *Salegentibacter agarivorans* | 30.1 | 103 | 0.15 |
| Uncharacterized protein | A0A5K1K8R5 | 2,756 |  | *Plasmodium falciparum* (isolate 3D7) | 22.7 | 106 | 0.15 |

**Table S3. Predicted eukaryotic cell localization prediction by DeepLoc 2.1 of symbiont proteins.** Localization of CI candidate constructs tested in this study and housekeeping GroL were predicted by running DeepLoc 2.1 in slow mode.

| **Gene** | **Localization** (probability) | **Membrane association**  (probability) | **Predicted signals** |
| --- | --- | --- | --- |
| **CAHE_0405** | Extracellular (0.9374) | Soluble (0.7290) | Signal peptide |
| **CAHE_0406** | Extracellular (0.5886) | Soluble (0.6560) | None |
| **CAHE_0757** | Extracellular (0.7616) | Soluble (0.6360) | None |
| **CAHE_p0044+signal** | Extracellular (0.7267) | Soluble (0.6800) | Nuclear localization signal |
| **CAHE_p0044** | Nucleus (0.8284)  Extracellular (0.6405) | Soluble (0.8280) | Nuclear localization signal |
| **CAHE_p0043-signal** | Cytoplasm (0.5161)  Nucleus (0.6517) | Soluble (0.7880) | Nuclear export signal |
| **CAHE_p0043** | Endoplasmic reticulum (0.5237) | Soluble (0.5460) | Signal peptide |
| **CAHE_0757-signal** | Cytoplasm (0.5749) | Soluble (0.6840) | Nuclear localization signal |
| **CAHE_0406-signal** | Cytoplasm (0.5423) | Soluble (0.8840) | None |
| **CAHE_0405-signal** | Nucleus (0.5681) | Soluble (0.7670) | None |
| **GroL** | Cytoplasm (0.7091) | Soluble (0.7270) | Nuclear export signal |

**Table S4. Evidence for additional putative cytoplasmic incompatibility (CI) effectors identified in this study.** Bolded ‘**Yes**’ indicates evidence supporting a role in CI.

| **Evidence** | **CAHE_0706** | **CAHE_0705** | **CAHE_0435** | **CAHE_0669** | **CAHE_465** |
| --- | --- | --- | --- | --- | --- |
| **Size (AA length)** | 336 | 80 | 392 | 314 | 485 |
| **Domains/Annotation** | Collagen triple helix repeat-containing lipoprotein | prokaryotic lipoprotein | Ankyrin repeat containing protein | Peptidyl-prolyl cis-trans isomerase | Serine protease with PDZ domain |
| **Possible partner** | CAHE_0705 | CAHE_0706 | none predicted | none predicted | none predicted |
| **Secreted based on proteome** | **Yes** | No | No | No | No |
| **Pupal testes detection** | **Yes** | **Yes** | **Yes** | **Yes** | **Yes** |
| **Seminal vesicle detection** | No | No | No | No | No |
| **Absence in non-reproductive organs** | No | **Yes** | **Yes** | **Yes** | No |
| **Toxicity in yeast** | not tested | not tested | not tested | not tested | not tested |

**TABLE S5. Samples of *Encarsia suzannae* carrying *Cardinium c*Eper1 used for proteomics.**

| **Sample** **Name** | **Sample** **Type** | **Host** **Sex** | **Host** **Life** **Stage** | **Number** **Of** **Specimens** **Pooled** |
| --- | --- | --- | --- | --- |
| **Pup_SPM_P1** | Fractionated - P | male | pupae | 100 |
| **Pup_SPM_P2** | Fractionated - P | male | pupae | 100 |
| **Pup_SPM_P3** | Fractionated - P | male | pupae | 100 |
| **Pup_SPM_P4** | Fractionated - P | male | pupae | 100 |
| **Pup_SPM_P5** | Fractionated - P | male | pupae | 100 |
| **Pup_SPM_PS1** | Fractionated - PS | male | pupae | 100 |
| **Pup_SPM_PS2** | Fractionated - PS | male | pupae | 100 |
| **Pup_SPM_PS3** | Fractionated - PS | male | pupae | 100 |
| **Pup_SPM_PS4** | Fractionated - PS | male | pupae | 100 |
| **Pup_SPM_S1** | Fractionated - S | male | pupae | 100 |
| **Pup_SPM_S2** | Fractionated - S | male | pupae | 100 |
| **Pup_SPM_S3** | Fractionated - S | male | pupae | 100 |
| **Pup_SPM_S4** | Fractionated - S | male | pupae | 100 |
| **Pup_SPM_S5** | Fractionated - S | male | pupae | 100 |
| **SLF_P1** | Fractionated - P | female | pupae | 100 |
| **SLF_P2** | Fractionated - P | female | pupae | 100 |
| **SLF_P3** | Fractionated - P | female | pupae | 100 |
| **SLF_P4** | Fractionated - P | female | pupae | 100 |
| **SLF_PS1** | Fractionated - PS | female | pupae | 100 |
| **SLF_PS2** | Fractionated - PS | female | pupae | 100 |
| **SLF_PS3** | Fractionated - PS | female | pupae | 100 |
| **SLF_PS4** | Fractionated - PS | female | pupae | 100 |
| **SLF_S1** | Fractionated - S | female | pupae | 100 |
| **SLF_S2** | Fractionated - S | female | pupae | 100 |
| **SLF_S3** | Fractionated - S | female | pupae | 100 |
| **SLF_S4** | Fractionated - S | female | pupae | 100 |
| **SPF_P1** | Fractionated - P | female | adult | 100 |
| **SPF_P2** | Fractionated - P | female | adult | 100 |
| **SPF_P3** | Fractionated - P | female | adult | 100 |
| **SPF_P4** | Fractionated - P | female | adult | 100 |
| **SPF_PS1** | Fractionated - PS | female | adult | 100 |
| **SPF_PS2** | Fractionated - PS | female | adult | 100 |
| **SPF_PS3** | Fractionated - PS | female | adult | 100 |
| **SPF_PS4** | Fractionated - PS | female | adult | 100 |
| **SPF_S1** | Fractionated - S | female | adult | 100 |
| **SPF_S2** | Fractionated - S | female | adult | 100 |
| **SPF_S3** | Fractionated - S | female | adult | 100 |
| **SPF_S4** | Fractionated - S | female | adult | 100 |
| **SPM_P1_2** | Fractionated - P | male | adult | 100 |
| **SPM_P2_2** | Fractionated - P | male | adult | 100 |
| **SPM_P3_2** | Fractionated - P | male | adult | 100 |
| **SPM_P4_2** | Fractionated - P | male | adult | 100 |
| **SPM_PS1_2** | Fractionated - PS | male | adult | 100 |
| **SPM_PS2_2** | Fractionated - PS | male | adult | 100 |
| **SPM_PS3_2** | Fractionated - PS | male | adult | 100 |
| **SPM_PS4_2** | Fractionated - PS | male | adult | 100 |
| **SPM_S1_2** | Fractionated - S | male | adult | 100 |
| **SPM_S2_2** | Fractionated - S | male | adult | 100 |
| **SPM_S3_2** | Fractionated - S | male | adult | 100 |
| **SPM_S4_2** | Fractionated - S | male | adult | 100 |
| **SSp1** | seminal vesicle contents | male | adult | 66 |
| **SSp2** | seminal vesicle contents | male | adult | 62 |
| **SSp3** | seminal vesicle contents | male | adult | 60 |
| **SSp4** | seminal vesicle contents | male | adult | 62 |
| **EP1** | Tissue - testes | male | pupae | 60 |
| **EP2** | Tissue - testes | male | pupae | 56 |
| **EP3** | Tissue - testes | male | pupae | 56 |
| **EP4** | Tissue - testes | male | pupae | 52 |
| **EP5** | Tissue - testes | male | pupae | 53 |
| **EA1** | Tissue - testes | male | adult | 51 |
| **EA2** | Tissue - testes | male | adult | 51 |
| **EA3** | Tissue - testes | male | adult | 64 |
| **EA4** | Tissue - testes | male | adult | 60 |
| **SIOV1** | Tissue - ovaries | female | adult | 57 |
| **SIOV2** | Tissue - ovaries | female | adult | 57 |
| **SIOV3** | Tissue - ovaries | female | adult | 59 |
| **SIOV4** | Tissue - ovaries | female | adult | 55 |
| **SIMB1** | Tissue - head | male | adult | 64 |
| **SIMB2** | Tissue - head | male | adult | 64 |
| **SIMB3** | Tissue - head | male | adult | 64 |
| **SIMB4** | Tissue - head | male | adult | 69 |
| **SIFB1** | Tissue - head | female | adult | 84 |
| **SIFB2** | Tissue - head | female | adult | 72 |
| **SIFB3** | Tissue - head | female | adult | 73 |
| **SIFB4** | Tissue - head | female | adult | 67 |

**TABLE S6. LC-MS/MS methods for fractionated samples.**

| **Parameter** | **Setting** |
| --- | --- |
| Wash step included? | One wash step between samples |
| Block randomization? | YES |
| Peptide load? | 600 ng |
| **Liquid Chromatography** | |
| LC | UltiMate 3000 RSLCnano Liquid Chromatograph NCS-3500RS Nano ProFlow (Thermo Fisher Scientific) |
| LC trap column | Nano trap cartridge, 300 µm i.d. x 5 mm, packed with Acclaim PepMap100 C18, 5 µm, 100 angstrom, nanoViper (Thermo Fisher Scientific, PN ES905, 160454) |
| LC separation column | 75µm × 75cm analytical EASY-Spray column packed with PepMap RSLC C18, 2 µm material (Thermo Fisher Scientific, SN C20296606) |
| Column temperature (separation) | 60°C |
| flow | 300 nl/min |
| Solvent A | H2O + 0.1% formic acid (FA) |
| Solvent B | 80% acetonitrile + 20% H2O + 0.1% FA |
| Step1: 5% to 31% B | 102 min |
| Step2: 31% to 50% B | 18 min |
| Step3: 99% B | 20 min |
| **Mass Spectrometry** | |
| MS | Q Exactive HF Hybrid Quadrupole-Orbitrap Mass Spectrometer (Thermo Fisher Scientific) |
| Ion source | NSI |
| Spray voltage | Static |
| Positive ion (V) | 2000 |
| Transfer tube temperature | 275°C |
| **Global Settings** |  |
| Expected LC peak width (s) | 15 |
| Default charge state | 2 |
| Internal mass calibration | User-defined lock mass |
| mass tolerance (ppm) | 15 |
| lock mass injection | FALSE |
| lock mass injection | 445.12003, Positive |
| acquisition start time (min) | 15 |
| acquisition end time (min) | 140 |
| **Full Scan** |  |
| Orbitrap Resolution | 60000 |
| Scan range (m/z) | 380-1600 |
| RF lens (%) | 40 |
| Normalized AGC target (%; ions) | 3.00E+06 |
| Max IT (ms) | 200 |
| Microscans | 1 |
| Polarity | Positive |
| **Filters** |  |
| Peptide match | Preferred |
| Intensity threshold | 5.00E+03 |
| Charge state exclusion | Exclude +1 |
| Include undetermined charge states | TRUE |
| Multiple charge | All |
| Dynamic exclusion duration (s) | 25 |
| DE: exclude isotopes | TRUE |
| **ddMS2** |  |
| Fragment tolerance | 0.1 Da |
| Precursor tolerance | 10 ppm |
| Multiplex ions | FALSE |
| Isolation window (m/z) | 1.2 |
| Isolation offset (m/z) | 0 |
| Normalized collision energy | 24 |
| Orbitrap resolution | 15000 |
| Scan range mode | Define first mass |
| Fixed first mass | - |
| Normalized AGC Target (%; ions) | 5.00E+04 |
| Max IT (ms) | 100 |
| Microscans | 1 |
| Loop count | 15 |
| MSX count | 1 |
| TopN | 15 |

**TABLE S7. LC-MS/MS method used for measurement of tissue samples.**

| **Parameter** | **Setting** |
| --- | --- |
| Wash step included? | One wash step between samples |
| Block randomization? | YES |
| Peptide load? | 600 ng |
| **Liquid Chromatography** | |
| LC | UltiMate 3000 RSLCnano Liquid Chromatograph NCS-3500RS Nano ProFlow (Thermo Fisher Scientific) |
| LC trap column | Nano trap cartridge, 300 µm i.d. x 5 mm, packed with Acclaim PepMap100 C18, 5 µm, 100 angstrom, nanoViper (Thermo Fisher, PN ES905, 160454) |
| LC separation column | 75µm × 75cm analytical EASY-Spray column packed with PepMap RSLC C18, 2 µm material (Thermo Fisher Scientific, SN C20296606) |
| Column temperature (separation) | 60°C |
| flow | 300 nl/min |
| Solvent A | H2O + 0.1% formic acid (FA) |
| Solvent B | 80% acetonitrile + 20% H2O + 0.1% FA |
| Step1: 5% to 31% B | 102 min |
| Step2: 31% to 50% B | 18 min |
| Step3: 99% B | 20 min |
| **Mass Spectrometry** | |
| MS | Orbitrap Exploris480 (Thermo Fisher Scientific) |
| Ion Source | NSI |
| Spray voltage | Static |
| Positive Ion (V) | 2000 |
| Transfer tube temperature | 275°C |
| **Global Settings** |  |
| Expected LC peak width (s) | 15 |
| Advanced peak determination | TRUE |
| Default charge state | 2 |
| Internal mass calibration | User-defined lock mass |
| Mode | Scan-to-scan |
| Mass tolerance (ppm) | 15 |
| Lock mass injection | FALSE |
| Lock mass injection | 445.12003, Positive |
| Acquisition start time (min) | 15 |
| Acquisition end time (min) | 140 |
| **Full Scan** |  |
| Orbitrap resolution | 60000 |
| Scan range (m/z) | 380-1600 |
| RF lens (%) | 40 |
| Normalized AGC target (%; ions) | 300; 3e6 |
| Max IT (ms) | 200 |
| Microscans | 1 |
| Polarity | Positive |
| **Filters** |  |
| Monoisotopic peak determination | Peptide (relax when too few precursors are found) |
| Intensity threshold | 5.00E+03 |
| Include charge state(s) | 2-6 |
| Include undetermined charge states | TRUE |
| Dynamic exclusion mode | Custom |
| DE: Exclude after n times | 1 |
| DE: Exclusion duration (s) | 25 |
| DE: mass tolerance | ppm |
| DE: low, high | 10, 10 |
| DE: exclude isotopes | TRUE |
| **ddMS2** |  |
| Multiplex ions | FALSE |
| Isolation Window (m/z) | 1.4 |
| Isolation Offset (m/z) | 0.3 |
| Reported Mass | Offset mass |
| Collision Energy Type | Normalized |
| HCD collision energy (%) | 27 |
| Orbitrap Resolution | 15000 |
| Scan range mode | Define first mass |
| First mass (m/z) | 100 |
| Normalized AGC Target (%; ions) | 100; 1e5 |
| Max IT (ms) | 50 |
| Microscans | 1 |

**TABLE S8. LC-MS/MS method used for measurement of yeast samples.**

| **Parameter** | **Setting** |
| --- | --- |
| Wash step included? | One wash step between samples |
| Block randomization? | YES |
| **Liquid Chromatography** | |
| LC | VanquishNeo (Thermo Scientific) |
| LC trap column | PepMap Neo 5 μm C18 300 μm X 5 mm Trap Cartridge (Thermo Scientific, REF 174500) |
| LC separation column | Easy-Spray PepMap Neo 2 μm C18 75 μm X 750 mm  (Thermo Scientific, PN ES75750PN) |
| Column temperature (separation) | 60°C |
| Flow | 250 nl/min |
| Solvent A | H2O + 0.1% formic acid (FA) |
| Solvent B | 80% acetonitrile + 20% H2O + 0.1% FA |
| Step1: 5% to 28% B | 39 min |
| Step2: 28% to 40% B | 14 min |
| Step3: 40% to 99% B | 2 min |
| Step4: 99% B | 15 min |
| **Mass Spectrometry** | |
| MS | Orbitrap Eclipse Mass Spectrometer (Thermo Scientific) |
| Ion Source | NSI |
| Spray voltage | Static |
| Positive Ion (V) | 2000 |
| Transfer tube temperature | 275°C |
| **Global Settings** |  |
| Advanced Peak Determination | TRUE |
| Default charge state | 2 |
| Internal mass calibration | User-defined lock mass |
| Lock mass injection | 445.12003, Positive |
| Acquisition start time (min) | 10 |
| Acquisition end time (min) | 70 |
| **Full Scan** |  |
| Orbitrap resolution | 60000 |
| Scan range (m/z) | 380-1600 |
| RF lens (%) | 40 |
| Normalized AGC target (%; ions) | 300% (1.2e6) |
| Max IT (ms) | 200 |
| Microscans | 1 |
| Polarity | Positive |
| **Filters** |  |
| MIPS mode | Peptide (relax when too few precursors are found) |
| Intensity threshold | 5.00E+03 |
| Include charge state(s) | 2-6 |
| include undetermined charge states | TRUE |
| Exclude within cycle | TRUE |
| Dynamic exclusion duration (s) | 25 |
| DE: exclude isotopes | TRUE |
| **ddMS2** |  |
| Scan range mode | Define first mass |
| First mass (m/z) | 100 |
| Isolation mode | Quadrupole |
| Isolation window (m/z) | 1.4 |
| Isolation offset (m/z) | 0.3 |
| Normalized collision energy | 27 |
| Orbitrap resolution | 15000 |
| Scan range mode | Define first mass |
| Fixed first mass | - |
| Normalized AGC target (%; ions) | 200%; 1.0e5 |
| Max IT (ms) | 50 |
| Microscans | 1 |
| Desired min. points across the peak | 6 |
| Polarity | Positive |
| TopN | 15 |

**TABLE S9. Software and packages used in this study.**

| **Tool/Software** | **Version** | **Use** | **Reference** |
| --- | --- | --- | --- |
| **ProteomeDiscoverer** | 2.3 | database search, protein identification and quantification |  |
| **DeepLoc** | 2.1 | protein annotation | (Ødum et al., 2024) |
| **TMHMM** | 2.0 | protein annotation | (Krogh et al., 2001) |
| **SignalP** | 5.0 | protein annotation | (Teufel et al., 2022) |
| **TargetP** | 2 | protein annotation | (Almagro Armenteros et al., 2019a) |
| **MANTIS** |  | protein annotation | (Queirós et al., 2021) |
| **InterProScan** |  | protein annotation | (Blum et al., 2025; Jones et al., 2014; Paysan-Lafosse et al., 2025) |
| **BLAST Koala** |  | protein annotation | (Kanehisa et al., 2016) |
| **R** | 4.2.2 | data analysis, visualization, and statisitics |  |
| **Adobe Illustrator** | 25.2.3 | data visualization |  |
| **R::dplyr** | 1.1.4 | data cleaning |  |
| **R::tidyr** | 1.3.1 | data cleaning |  |
| **R::forcats** | 1.0.0 | data cleaning |  |
| **R::broom** | 1.0.8 | data cleaning |  |
| **R::ggplot2** | 3.5.2 | data visualization |  |
| **R::ggpubr** | 0.6.1 | data visualization |  |
| **R::pheatmap** | 1.0.13 | data visualization |  |
| **R::compositions** | 2.0-8 | data normalization |  |
| **Gene Graphics** |  | data visualization | (Harrison et al., 2018) |
| **BioEdit** | 7.2.5 | data analysis | (Hall, 1999) |
| **Jalview** | 2.11.5.1 |  | (Troshin et al., 2018, 2011; Waterhouse et al., 2009) |

**Table S10. Plasmids used in Gateway cloning for preparation of constructs used in heterologous expression yeast assays.**

|  | **pYES-DEST52** | **pAG415GAL-ccdb-EGFP** |
| --- | --- | --- |
| **Abbreviation** | pYES | pAG (pAG415) |
| **Plasmid type** | 2- micron | CEN (centromeric) |
| **Copy number** | High (40-60 copies) | Low (1-4 copies) |
| **Selective marker (yeast)** | Uracil (URA3 gene) | Leucine (LEU2 gene) |
| **Selective marker (bacteria)** | | |
| *Empty plasmid* | chloramphenicol (30 μg/mL) and ampicillin (100 μg/mL) | |
| *With gene insert* | ampicillin (100 μg/mL) | |
| **Counter-selection (bacteria)** | *E. coli* ccdb toxin on empty vector; need ccdb-resistant *E. coli* for maintenance of empty plasmid | |
| **Promoter** | GAL1 (galactose-inducible) | |
| **Terminator** | CYC1 | |
| **Tags** | C-terminal V5 epitope and polyhistidine (6x His) tags (not used) | C-terminal eGFP (enhanced green fluorescent protein) tag (used only for localization assays) |
| **Source** | ThermoFisher (12286019) | Addgene plasmid #14193 |

**TABLE S11. Yeast strain and constructs used in this study.**

| **Construct** | **Confirmation Method** | **Codon optimized for *S. cerevisiae*?** | **Description** | **Experiment notes** |
| --- | --- | --- | --- | --- |
| **High-copy single plasmid constructs** | | | | |
| pYES-DEST52 empty | PCR + band size | N/A | High-copy plasmid with no modifications or gene additions | High-copy yeast vector (2-micron) |
| pYES-DEST52 + wPip CidB | PCR + band size | No | High-copy plasmid with unaltered Wolbachia wPip cidB | Pos. control; >Wolbachia_wPip_CidB_WP0283_AM999887.1:289303-292827 |
| pYES-DEST52 + CAHE_p0043 | PCR + band size | No | High-copy plasmid with unaltered CAHE_p0043 |  |
| pYES-DEST52 + CAHE_p0044 | PCR + band size | No | High-copy plasmid with unaltered CAHE_p0044 |  |
| pYES-DEST52 + CAHE_0757 | PCR + band size | No | High-copy plasmid with unaltered CAHE_0757 |  |
| pYES-DEST52 + CAHE_0405 | PCR + band size | No | High-copy plasmid with unaltered CAHE_0405 |  |
| pYES-DEST52 + CAHE_0406 | PCR + band size | No | High-copy plasmid with unaltered CAHE_0406 |  |
| pYES-DEST52 + cEper2 CAHE_0406 homolog | PCR + band size | No | High-copy plasmid with unaltered cEper2 MGI2262105.1 | CAHE_0406 homolog |
| pYES-DEST52 + cEina2 CAHE_0406 homolog | PCR + band size | No | High-copy plasmid with unaltered cEina2 MGI2299658.1 | CAHE_0406 homolog |
| pYES-DEST52 + cEina3_1 CAHE_0406 homolog | PCR + band size | No | High-copy plasmid with unaltered cEina3 MGI2299057.1 | CAHE_0406 homolog |
| pYES-DEST52 + cEina3_2 CAHE_0406 homolog | PCR + band size | No | High-copy plasmid with unaltered cEina3 MGI2299058.1 | CAHE_0406 homolog |
| pYES-DEST52 + CAHE_0405-signal | PCR + band size + Sanger | Yes & Kozak sequence added | High-copy plasmid with codon-optimized CAHE_0405 after removing signal peptide | Signal peptide removed at SignalP predicted cut site |
| pYES-DEST52 + CAHE_0406-signal | PCR + band size + Sanger | Yes & Kozak sequence added | High-copy plasmid with codon-optimized CAHE_0406 after removing signal peptide | Signal peptide removed at SignalP predicted cut site |
| pYES-DEST52 + CAHE_0757-signal | PCR + band size + Sanger | Yes | High-copy plasmid with codon-optimized CAHE_0757 after signal peptide removal | Signal peptide removed at SignalP predicted cut site |
| pYES-DEST52 + CAHE_p0044+signal | PCR + band size + Sanger | Yes | High-copy plasmid with codon-optimized CAHE_p0044 with added potential signal peptide | Potential signal peptide added to N-terminus of protein by including amino acids beginning with an alternate upstream start codon |
| pYES-DEST52 + CAHE_p0043-signal | PCR + band size + Sanger | Yes | High-copy plasmid with codon-optimized CAHE_p0043 after signal peptide removal | Signal peptide removed at SignalP predicted cut site |
| **Co-expression constructs** | | | | |
| pAG415+CAHE_0405 & pYES+CAHE_0406 | PCR + band size |  | Coexpression strain with unmodified CAHE_0405 on the low-copy plasmid and unmodified CAHE_0406 on the high-copy plasmid |  |
| pAG415-empty & pYES-empty | PCR + band size |  | Coexpression strain with empty high and low-copy plasmids (no genes inserted); coexpression negative control |  |
| pAG415-empty & pYES+CAHE_0406 | PCR + band size |  | Coexpression strain with unmodified CAHE_0406 on the high-copy plasmid and an empty low-copy plasmid (no genes inserted) |  |
| pAG415+CAHE_p0043-signal & pYES empty | PCR + band size |  | Coexpression strain with CAHE_p0043 with signal peptide removed on the low-copy plasmid and an empty (no genes inserted) high-copy plasmid |  |
| pAG415+CAHE_p0043-signal & pYES + CAHE_p0044 | PCR + band size |  | Coexpression strain with CAHE_p0043 with signal peptide removed on the low-copy plasmid and unmodified CAHE_p0044 on the high-copy plasmid |  |
| pAG415+CAHE_p0043-signal & pYES + CAHE_p0044+signal | PCR + band size |  | Coexpression strain with CAHE_p0043 with signal peptide removed on the low-copy plasmid and codon optimized CAHE_p0044 with added potential signal peptide on the high-copy plasmid |  |

### Supplementary Figures

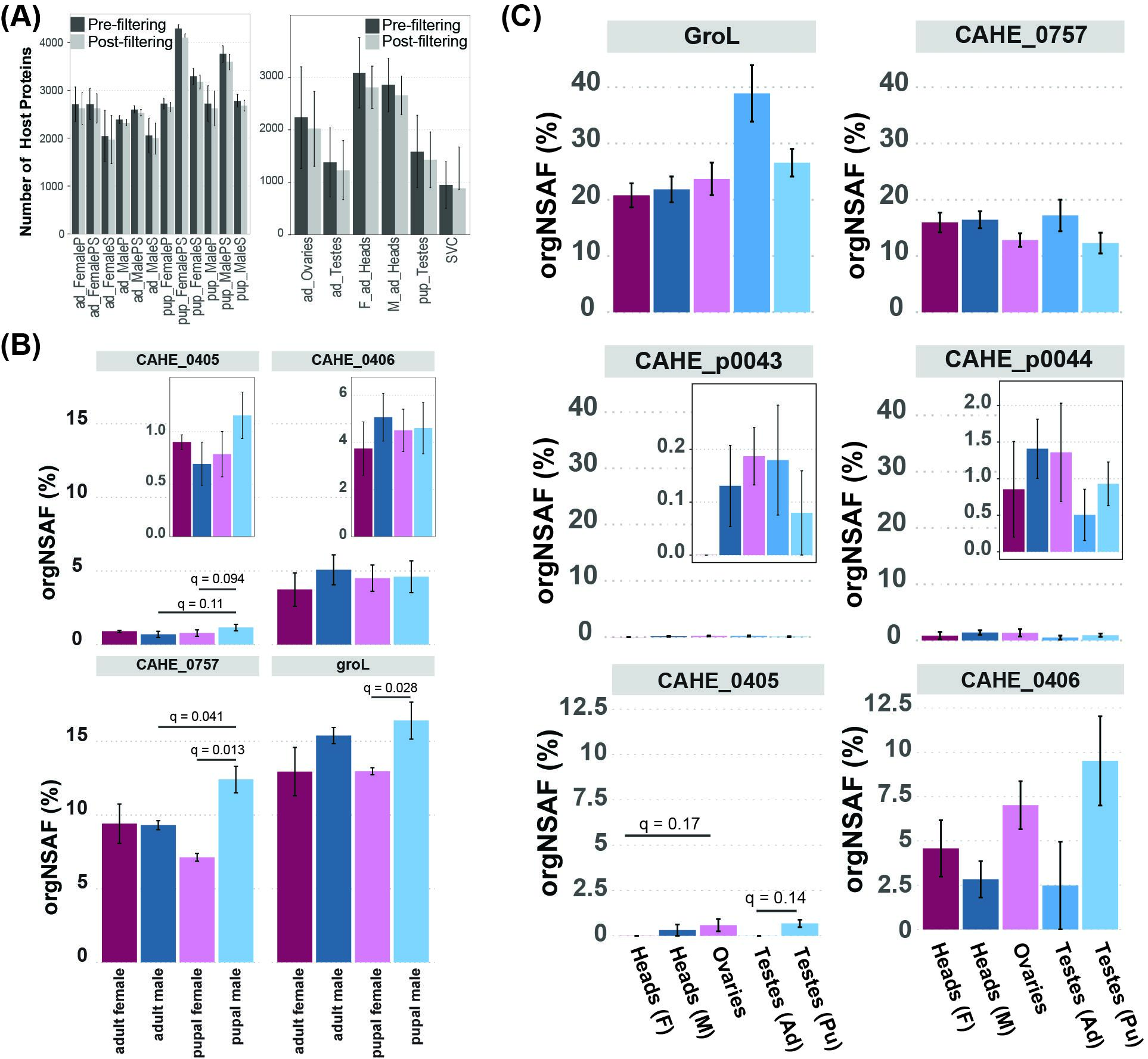

**Figure S1. Detection of host proteins and additional proteomic evidence for CI effector candidates. (A)** Mean number of *Encarsia* (host) proteins detected per sample type at a false discovery rate of 5% and classified as a master protein before and after filtering for detection in at least 50% of at least one sample type. Error bars indicate standard deviations (n =4, except for S and P fractions of pupal males where n = 5). SVC = adult seminal vesicle contents, ad = adult, pup = pupae, F = female, M = male, and P, PS, and S = sample fraction. **(B)** Mean abundance of CAHE_0405, CAHE_0406, CAHE_0757 and groL in the symbiont enrichment fraction (PS) with vertical error bars indicating standard error. **(C)** Mean relative abundance (orgNSAF%) of CAHE_p0043, CAHE_p0044, CAHE_0405, CAHE_0406, CAHE_0757, and GroL across tissues, with vertical error bars indicating standard error. For bar plots in (**B)** and (**C)**, **s**ignificance values (q < 0.2) are indicated by horizontal bar with q-value (Welch’s t-test with Benjamini-Hochberg FDR correction).

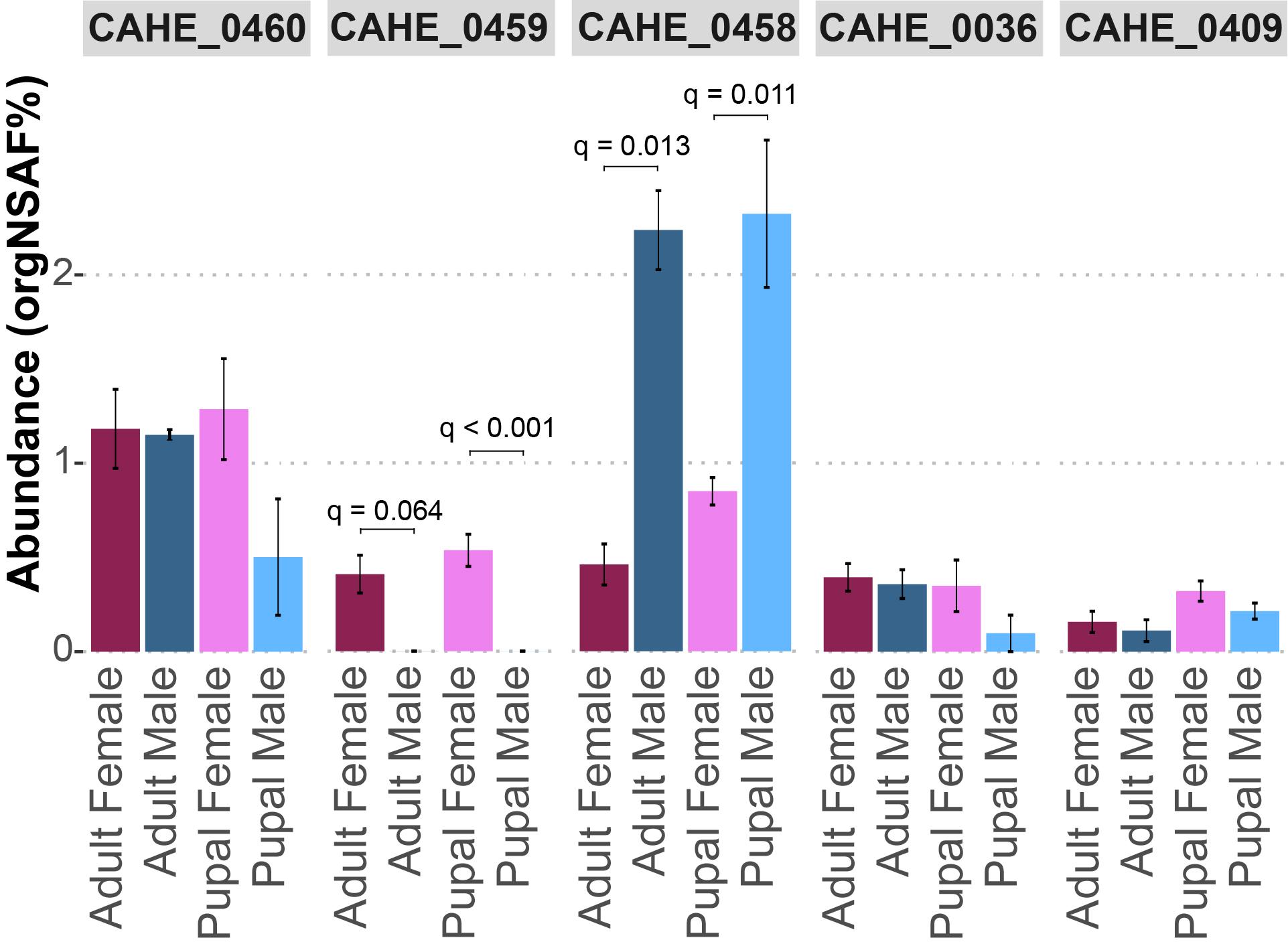

**Figure S2. Detection of the T6SS system of *Cardinium* across host sex and life stage.** Abundance (organismal normalized spectral abundance factor percentage, orgNSAF) of detected T6SS/Afp-like components. Significance was assessed using a Welch T-test on CLR normalized data with zeros imputed using a small constant (0.5 times the lowest value in the comparison). P-values were adjusted for multiple hypothesis testing using Benjamini-Hochberg FDR correction. Adjusted p-values are reported for q-values less than 0.1.

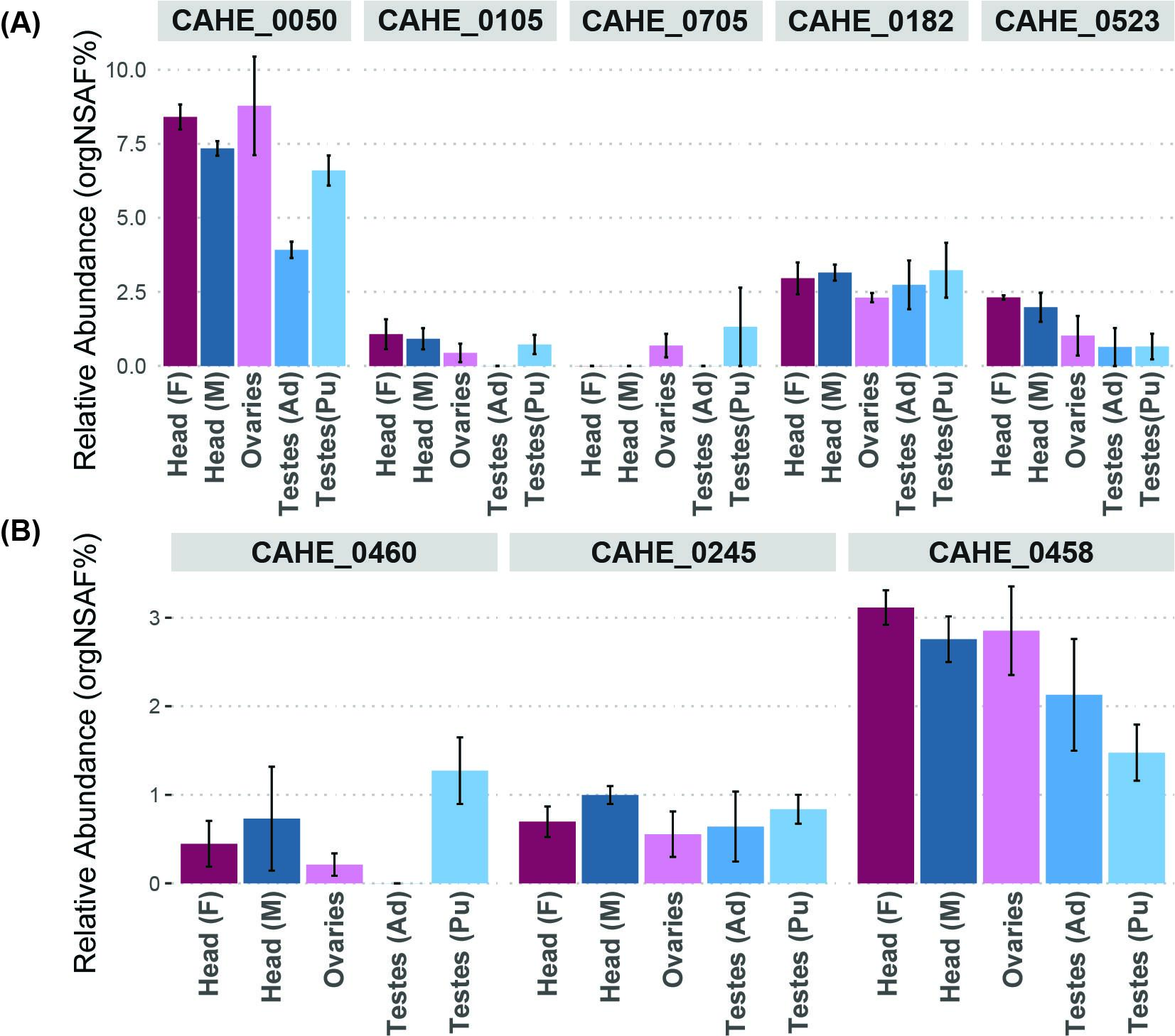

**Figure S3. Detection of additional proteins of interest in the 20 most abundant proteins of the pupal testis metaproteome across tissue types.** **(A)** Detection of uncharacterized proteins. **(B)** Detection of proteins likely involved in transport, e.g., T6SS (CAHE_0458, CAHE_0460) and solute binding domain containing protein (CAHE_0245).

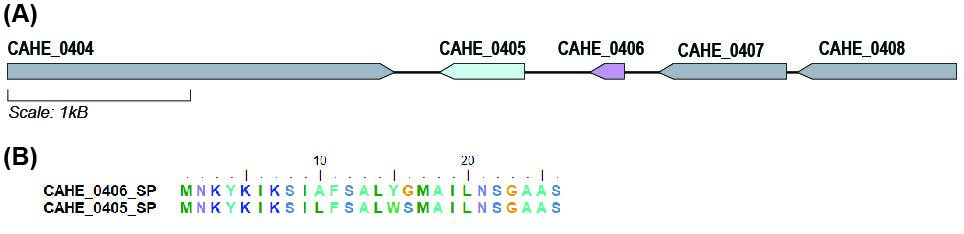

**Figure S4. Genomic structural evidence for CAHE_0405 and CAHE_0406. (A)** Gene neighborhood of CAHE_0405 (green) and CAHE_0406 (red) visualized with Gene Graphics (Harrison et al., 2018). **(B)** CAHE_0405 and CAHE_0406 signal peptide alignment (Alignment score: 109, Identities: 0.8846154, Similarities: 0.9230769, Similarity Matrix: BLOSUM62) performed with BioEdit.

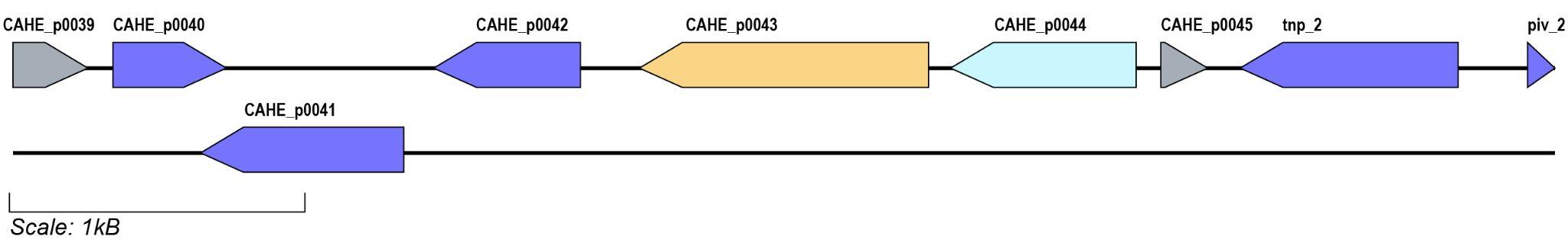

**Figure S5. Gene neighborhood of CAHE_p0043 and CAHE_p0044.** This was visualized using Gene Graphics (Harrison et al., 2018). Mobile genetic elements in this 5000 bp window (CAHE_p0040, CAHE_p0041, CAHE_p0042, tpn_2, and piv_2) are represented by purple tags.

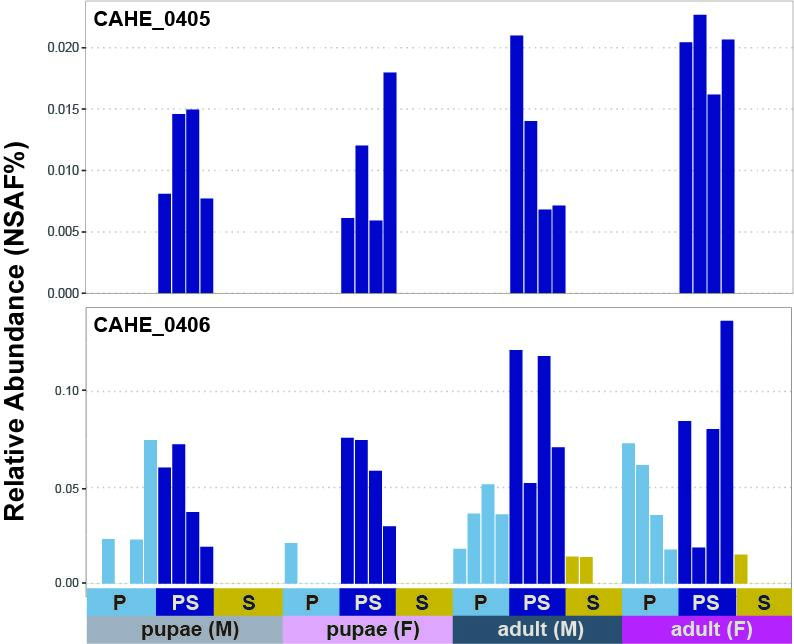

**Figure S6**. **Detection patterns of CAHE_0406 and CAHE_0405 across fractionated samples.** The x-axis shows individual samples grouped by fraction, host sex, and host life stage. Yellow = symbiont secretome (S), dark blue = symbiont cell enrichment (PS), and light blue = host cell debris fraction (P).

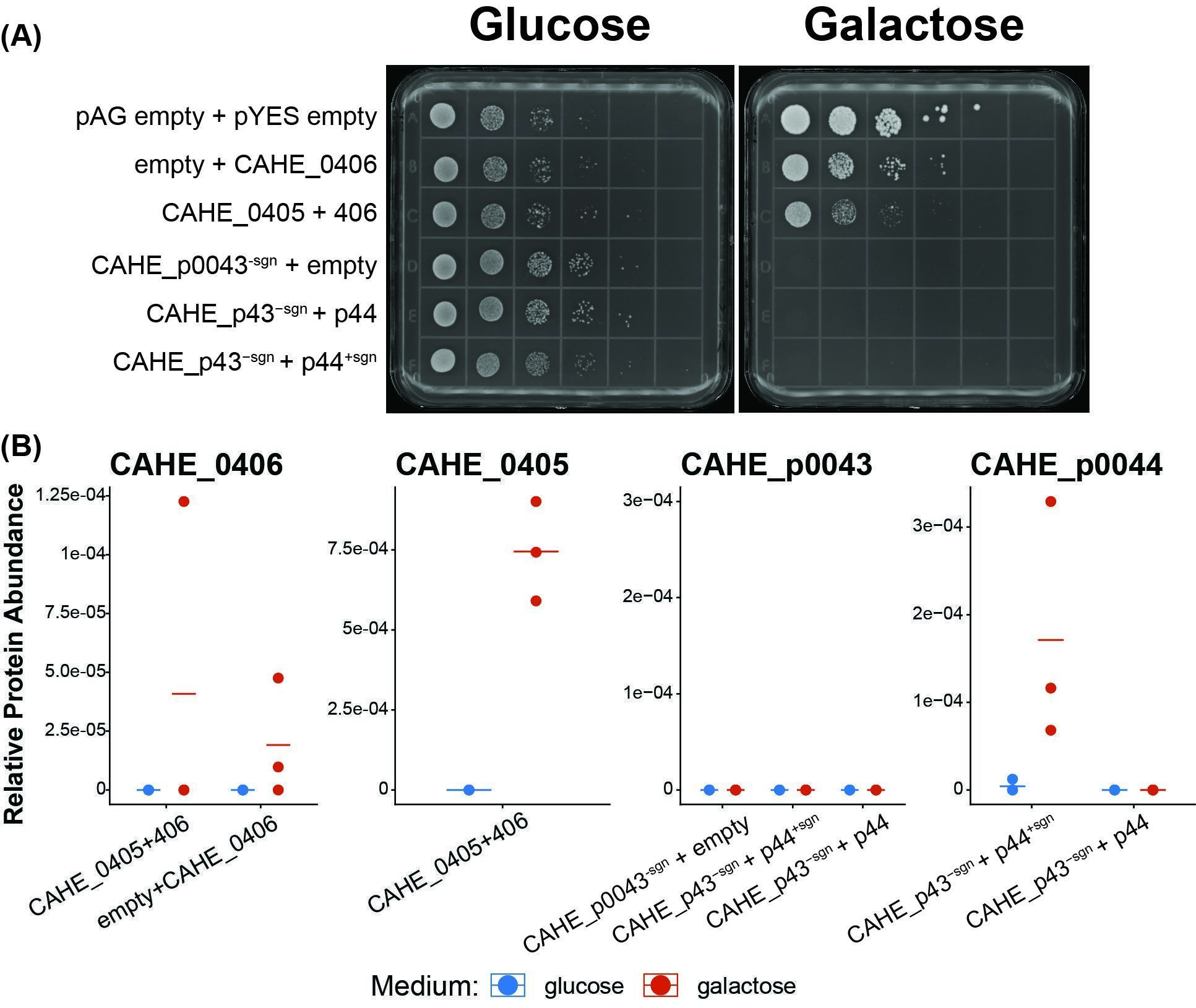

**FIGURE S7. Co-expression assay to attempt rescue yeast toxicity in heterologous expression assay.** (A) Spot plates for coexpression of CI candidate proteins with potential rescue factors using a heterologous expression assay in *Saccharomyces cerevisiae*. Left plate shows growth on DOB agar + 2% glucose where the genes of interest are turned off and serve as negative controls. Right plates are DOB agar with 2% galactose for induction of the gene of interest. Lack of growth on these plates indicates the expressed gene is toxic to yeast growth. Rescue is indicated by increased growth in rows with the rescue candidate expressed, relative to the rows with toxin candidates expressed with an empty vector. (B) Detection of constructs expressed in *S. cerevisiae* in LC-MS/MS measured proteome where each point is the abundance (TSS normalized AUC quantification) of the protein in one replicate and horizontal lines indicate the mean of three replicates.

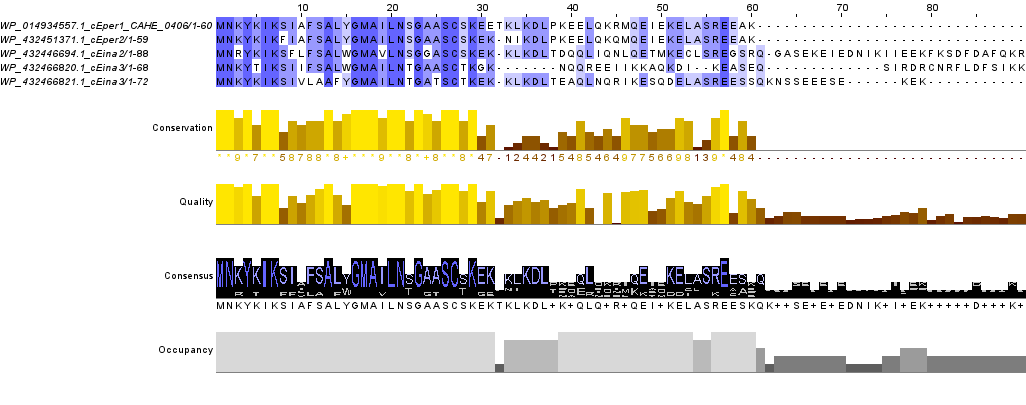

**FIGURE S8.** **Alignment of the *Cardinium* *c*Eper1 CI candidate protein CAHE_0406 to homologs from other *Cardinium* strains tested in yeast.** Amino acid sequences were retrieved from RefSeq and aligned using MAFFT with default settings in Jalview v. 2.11.5.1.

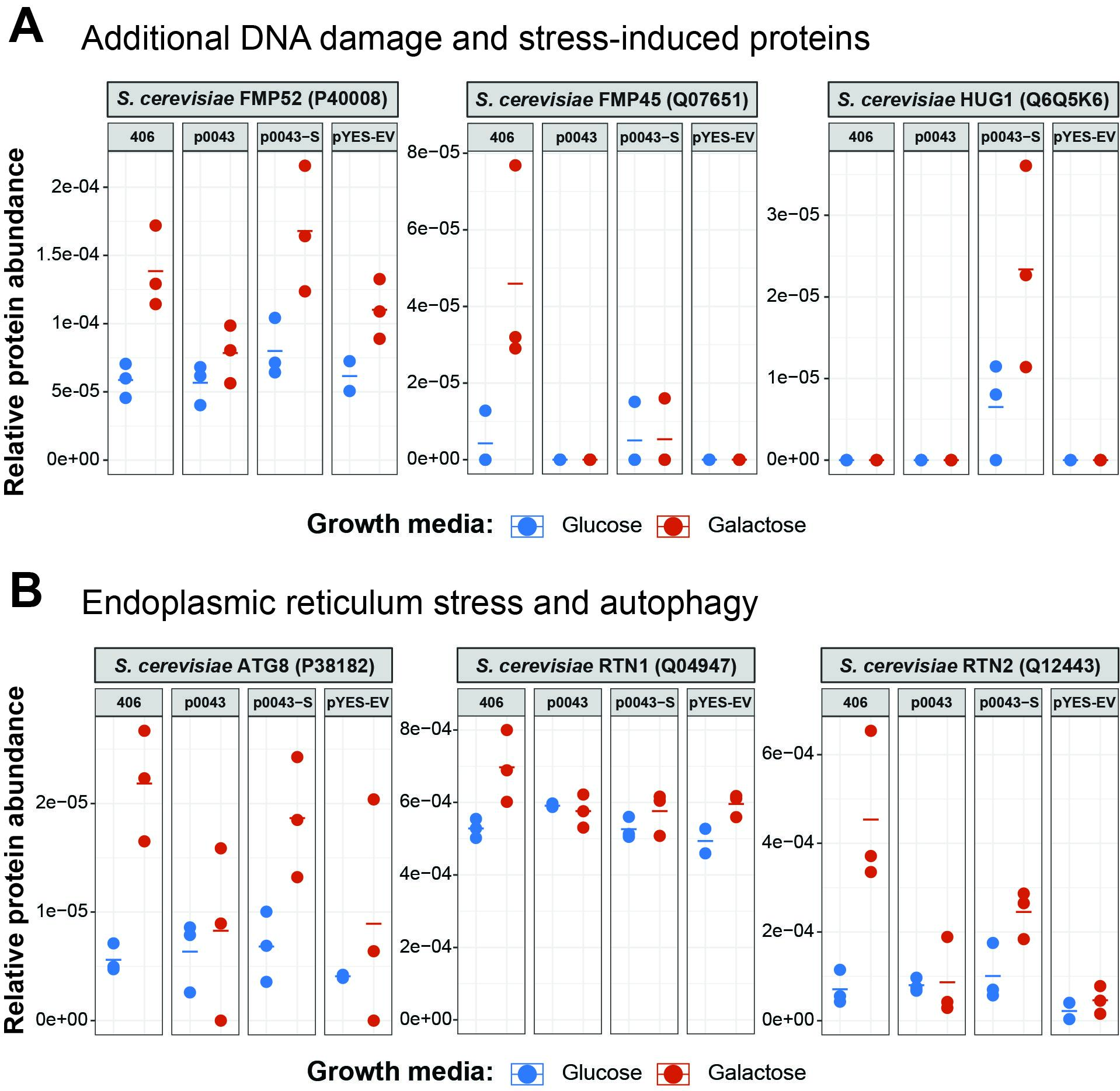

**FIGURE S9. *Saccharomyces cerevisiae* expression response to toxin induction.** (A) Expression of FMP52, FMP45, and HUG1. (B) Expression of ATG8, RTN1, and RTN2. Constructs: 406 = CAHE_0406, p0043 = CAHE_p0043, p0043-S = CAHE_p0043^-signal^, and pYES-EV = empty pYES-DEST52 vector. Each point is the abundance (TSS normalized AUC quantification) of the protein in one replicate and horizontal lines indicate the mean (n=3).

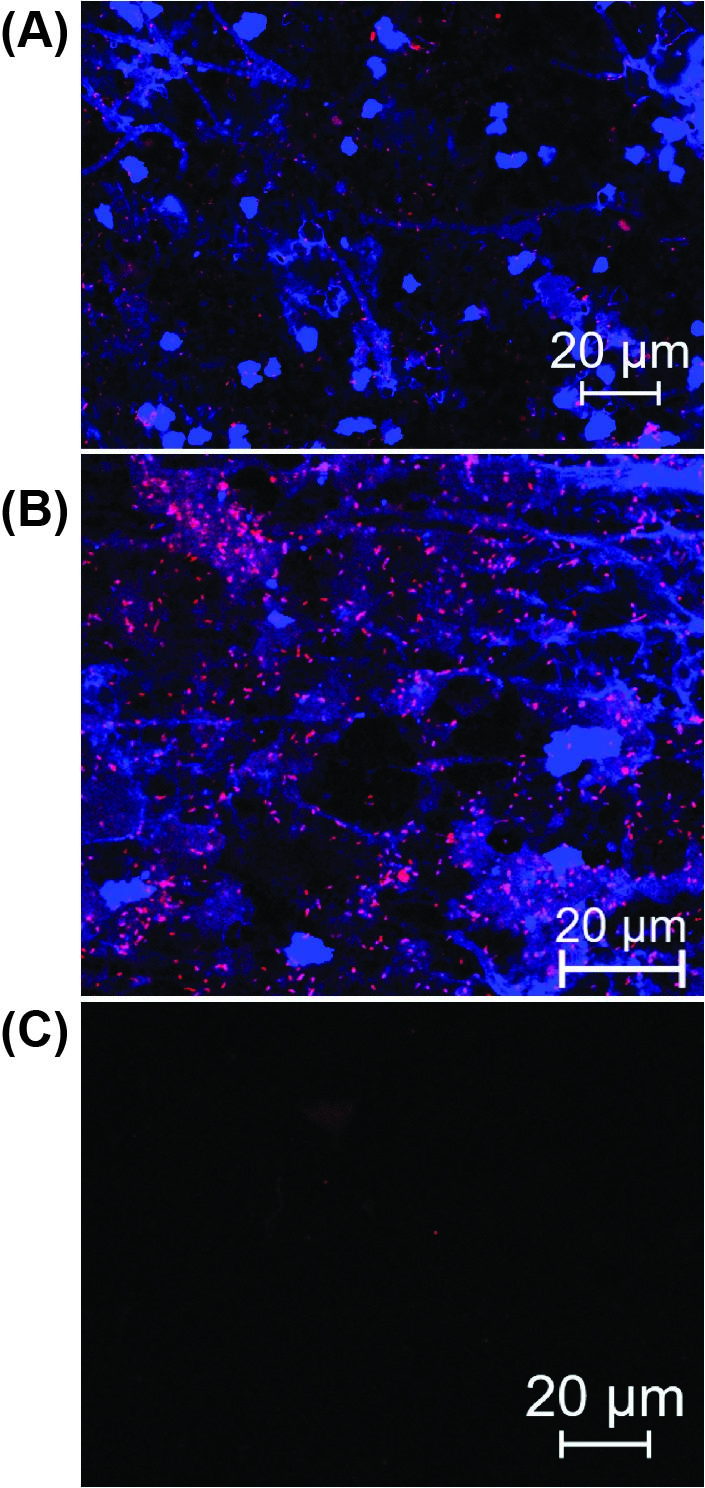

**Figure S10.** **Fluorescence microscopy of fractionated *Encarsia suzannae* wasp homogenates.** Host nuclei are stained with DAPI (blue) and symbionts are tagged with *Cardinium* 16S rRNA specific Cy3 double-labeled probes (pink). Panels: A = P fraction, B = PS fraction, C = S fraction.
