## Supplementary Data 2 for "Identification of cytoplasmic incompatibility effectors of the reproductive manipulator *Cardinium hertigii*"

**Supplementary Data 2.** Abundance of *c*Eper1 proteins across fractionated samples quantified by peptide spectral match counts normalized to Normalized Spectral Abundance Factors Percentages (NSAF%). Bars are colored according to fraction (P = light blue, PS = dark blue, S = gold).

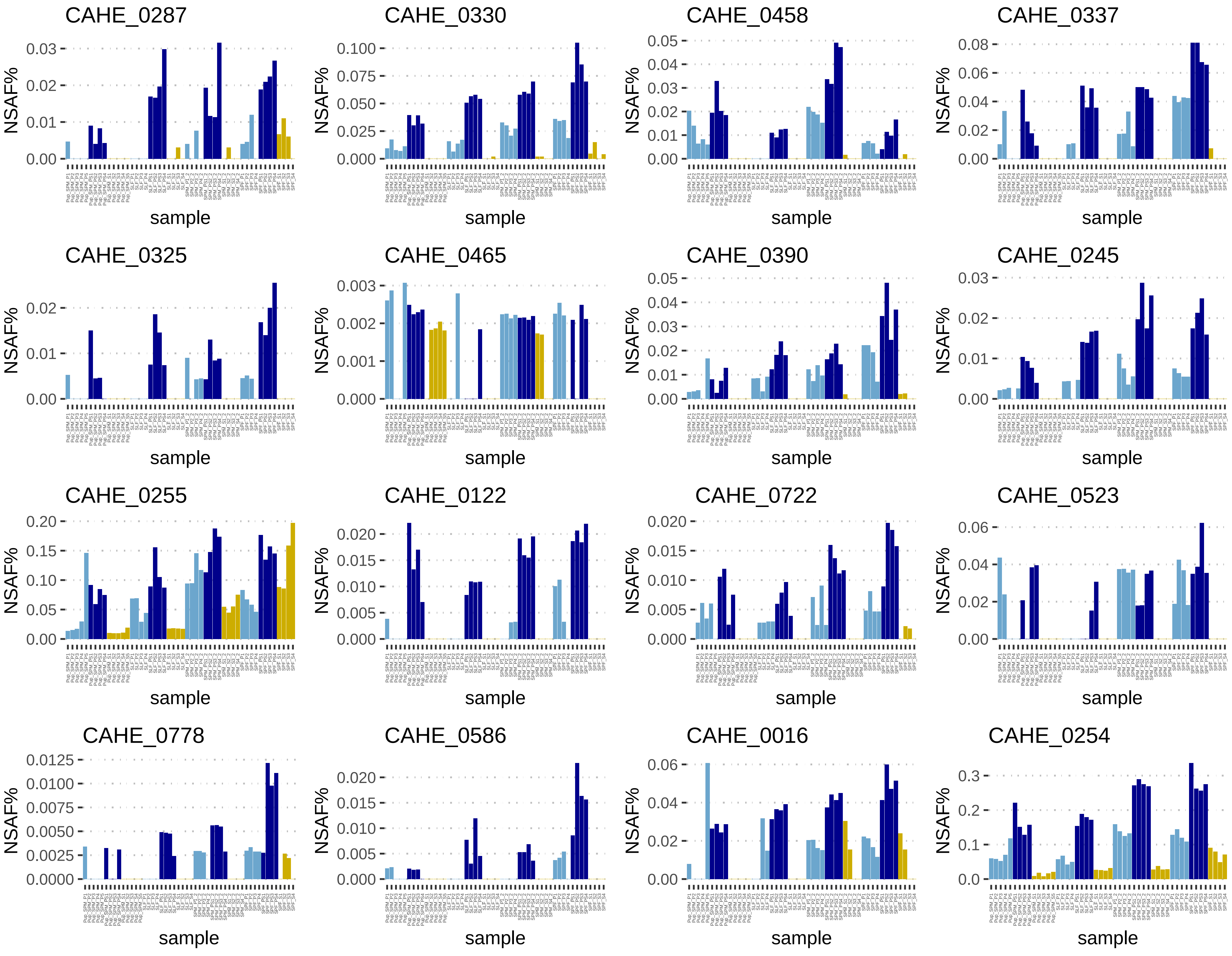

CAHE_0287
CAHE_0330
CAHE_0458
CAHE_0337
CAHE_0325
CAHE_0465
CAHE_0390
CAHE_0245
CAHE_0255
CAHE_0122
CAHE_0722
CAHE_0523
CAHE_0778
CAHE_0586
CAHE_0016
CAHE_0254
NSAF%
sample

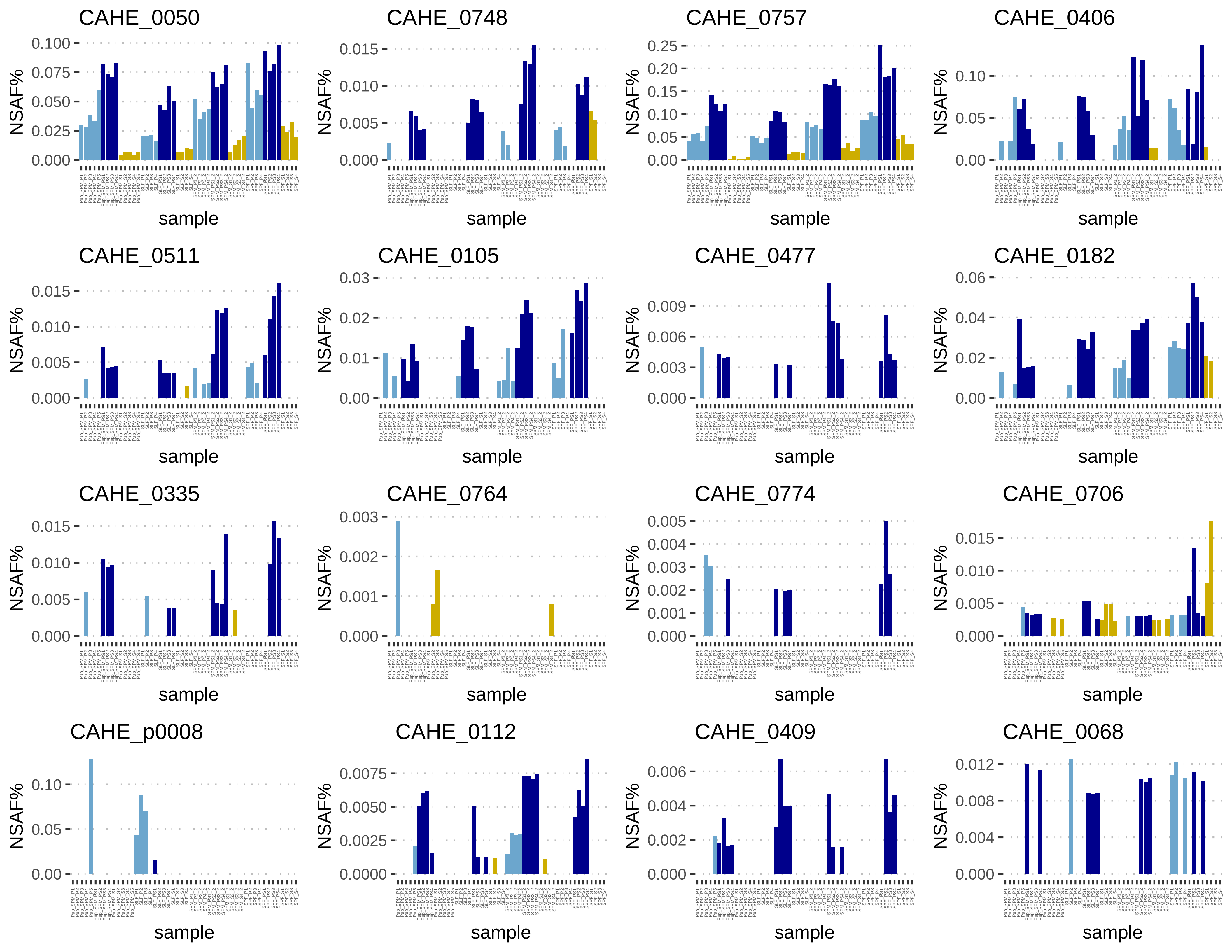
CAHE_0050
0.100
0.075
0.050
0.025
0.000
NSAF%
sample
CAHE_0748
0.015
0.010
0.005
0.000
NSAF%
sample
CAHE_0757
0.25
0.20
0.15
0.10
0.05
0.00
NSAF%
sample
CAHE_0406
0.10
0.05
0.00
NSAF%
sample
CAHE_0511
0.015
0.010
0.005
0.000
NSAF%
sample
CAHE_0105
0.03
0.02
0.01
0.00
NSAF%
sample
CAHE_0477
0.009
0.006
0.003
0.000
NSAF%
sample
CAHE_0182
0.06
0.04
0.02
0.00
NSAF%
sample
CAHE_0335
0.015
0.010
0.005
0.000
NSAF%
sample
CAHE_0764
0.003
0.002
0.001
0.000
NSAF%
sample
CAHE_0774
0.005
0.004
0.003
0.002
0.001
0.000
NSAF%
sample
CAHE_0706
0.015
0.010
0.005
0.000
NSAF%
sample
CAHE_p0008
0.10
0.05
0.00
NSAF%
sample
CAHE_0112
0.0075
0.0050
0.0025
0.0000
NSAF%
sample
CAHE_0409
0.006
0.004
0.002
0.000
NSAF%
sample
CAHE_0068
0.012
0.008
0.004
0.000
NSAF%
sample

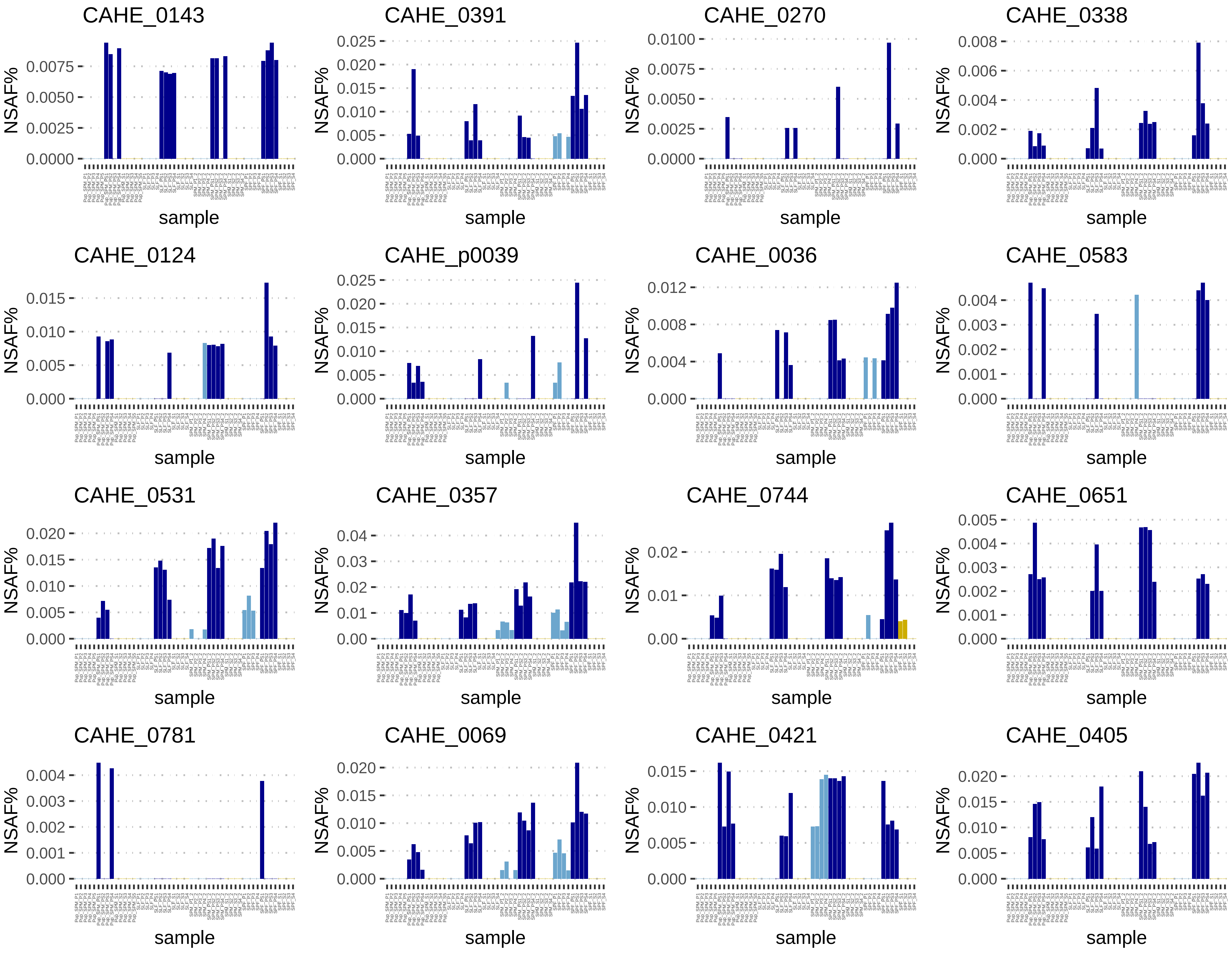

CAHE_0143
0.0075
0.0050
0.0025
0.0000
NSAF%
sample
CAHE_0391
0.025
0.020
0.015
0.010
0.005
0.000
NSAF%
sample
CAHE_0270
0.0100
0.0075
0.0050
0.0025
0.0000
NSAF%
sample
CAHE_0338
0.008
0.006
0.004
0.002
0.000
NSAF%
sample
CAHE_0124
0.015
0.010
0.005
0.000
NSAF%
sample
CAHE_p0039
0.025
0.020
0.015
0.010
0.005
0.000
NSAF%
sample
CAHE_0036
0.012
0.008
0.004
0.000
NSAF%
sample
CAHE_0583
0.004
0.003
0.002
0.001
0.000
NSAF%
sample
CAHE_0531
0.020
0.015
0.010
0.005
0.000
NSAF%
sample
CAHE_0357
0.04
0.03
0.02
0.01
0.00
NSAF%
sample
CAHE_0744
0.02
0.01
0.00
NSAF%
sample
CAHE_0651
0.005
0.004
0.003
0.002
0.001
0.000
NSAF%
sample
CAHE_0781
0.004
0.003
0.002
0.001
0.000
NSAF%
sample
CAHE_0069
0.020
0.015
0.010
0.005
0.000
NSAF%
sample
CAHE_0421
0.015
0.010
0.005
0.000
NSAF%
sample
CAHE_0405
0.020
0.015
0.010
0.005
0.000
NSAF%
sample

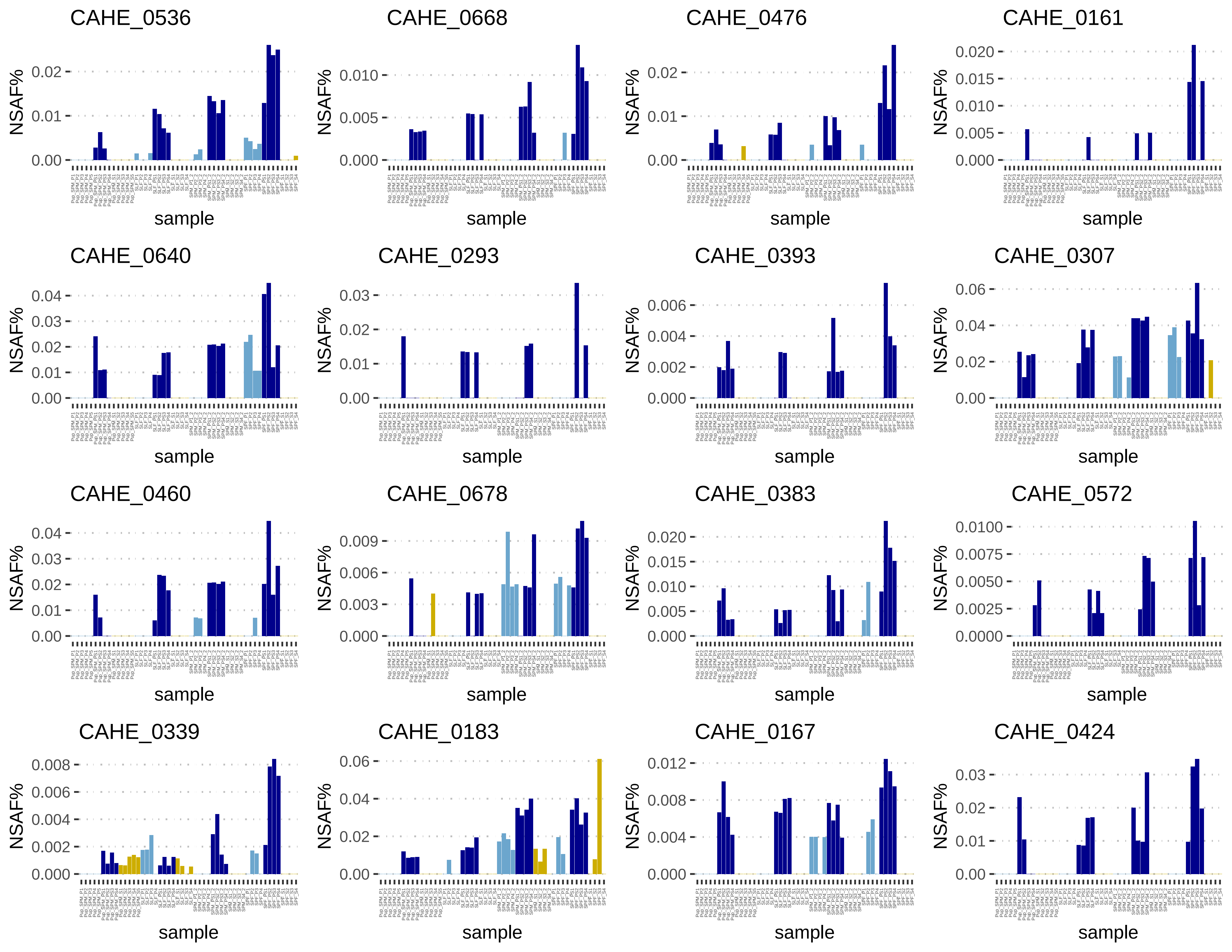
CAHE_0536
CAHE_0668
CAHE_0476
CAHE_0161
CAHE_0640
CAHE_0293
CAHE_0393
CAHE_0307
CAHE_0460
CAHE_0678
CAHE_0383
CAHE_0572
CAHE_0339
CAHE_0183
CAHE_0167
CAHE_0424
NSAF%
sample

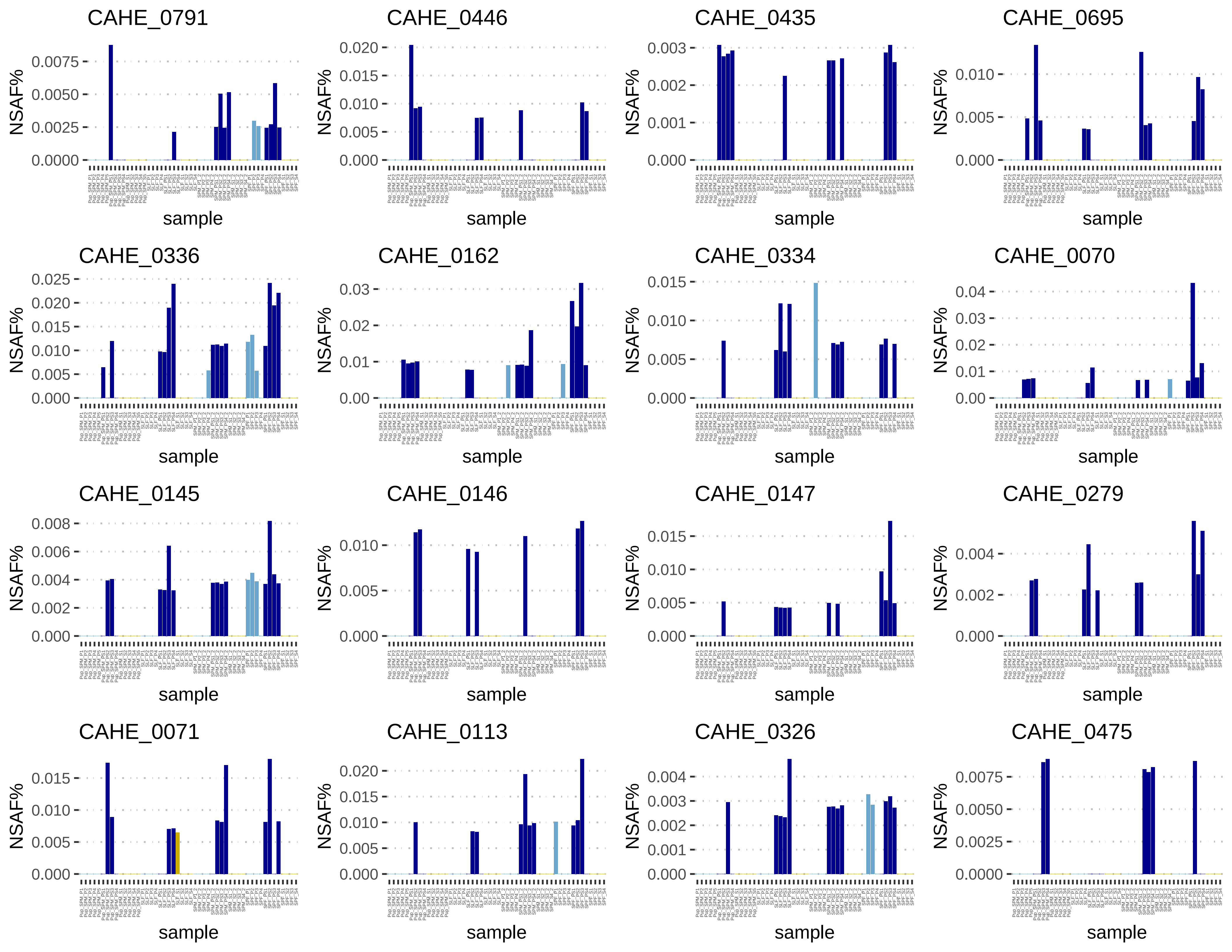

CAHE_0791
NSAF%
0.0075
0.0050
0.0025
0.0000
sample
CAHE_0446
NSAF%
0.020
0.015
0.010
0.005
0.000
sample
CAHE_0435
NSAF%
0.003
0.002
0.001
0.000
sample
CAHE_0695
NSAF%
0.010
0.005
0.000
sample
CAHE_0336
NSAF%
0.025
0.020
0.015
0.010
0.005
0.000
sample
CAHE_0162
NSAF%
0.03
0.02
0.01
0.00
sample
CAHE_0334
NSAF%
0.015
0.010
0.005
0.000
sample
CAHE_0070
NSAF%
0.04
0.03
0.02
0.01
0.00
sample
CAHE_0145
NSAF%
0.008
0.006
0.004
0.002
0.000
sample
CAHE_0146
NSAF%
0.010
0.005
0.000
sample
CAHE_0147
NSAF%
0.015
0.010
0.005
0.000
sample
CAHE_0279
NSAF%
0.004
0.002
0.000
sample
CAHE_0071
NSAF%
0.015
0.010
0.005
0.000
sample
CAHE_0113
NSAF%
0.020
0.015
0.010
0.005
0.000
sample
CAHE_0326
NSAF%
0.004
0.003
0.002
0.001
0.000
sample
CAHE_0475
NSAF%
0.0075
0.0050
0.0025
0.0000
sample

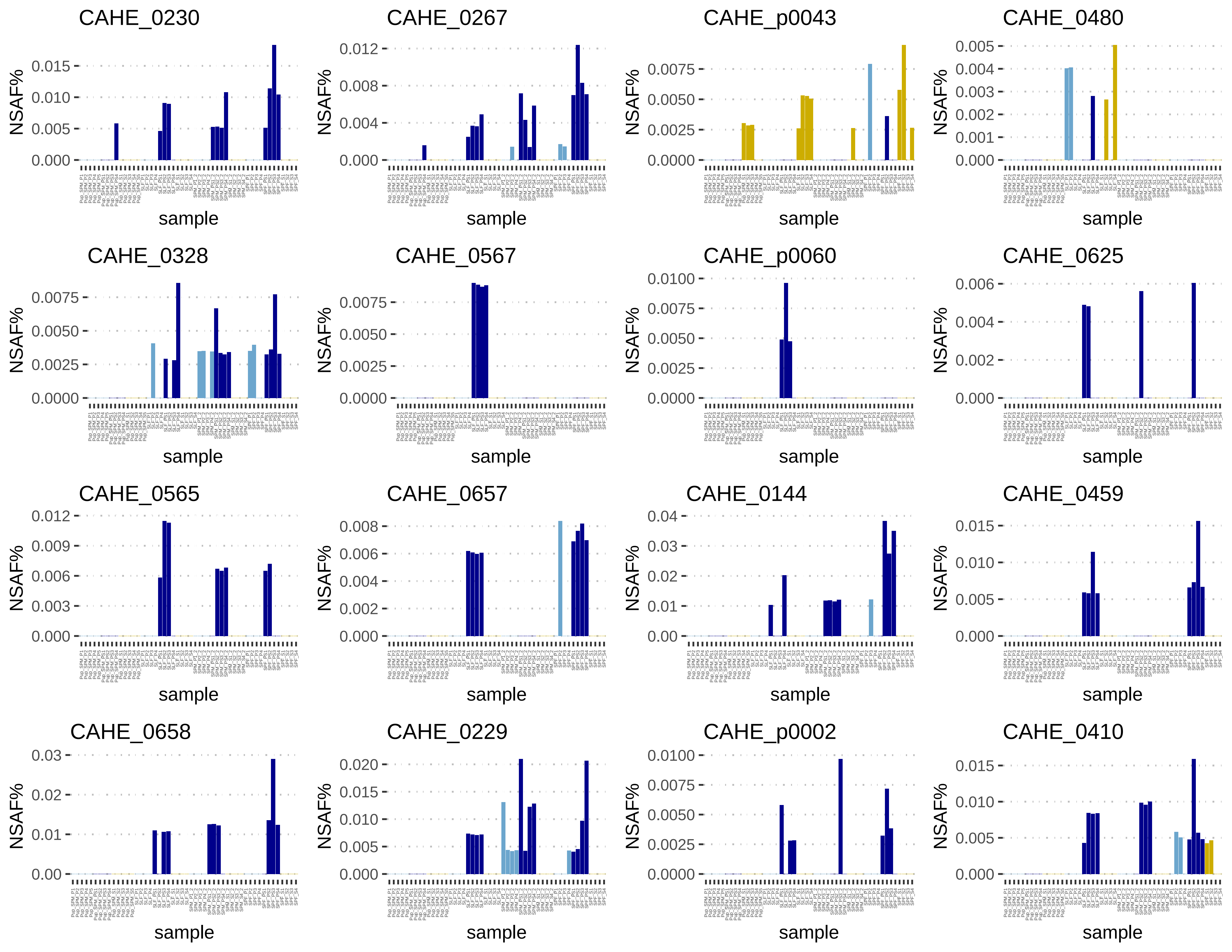
CAHE_0230
CAHE_0267
CAHE_p0043
CAHE_0480
CAHE_0328
CAHE_0567
CAHE_p0060
CAHE_0625
CAHE_0565
CAHE_0657
CAHE_0144
CAHE_0459
CAHE_0658
CAHE_0229
CAHE_p0002
CAHE_0410
NSAF%
sample

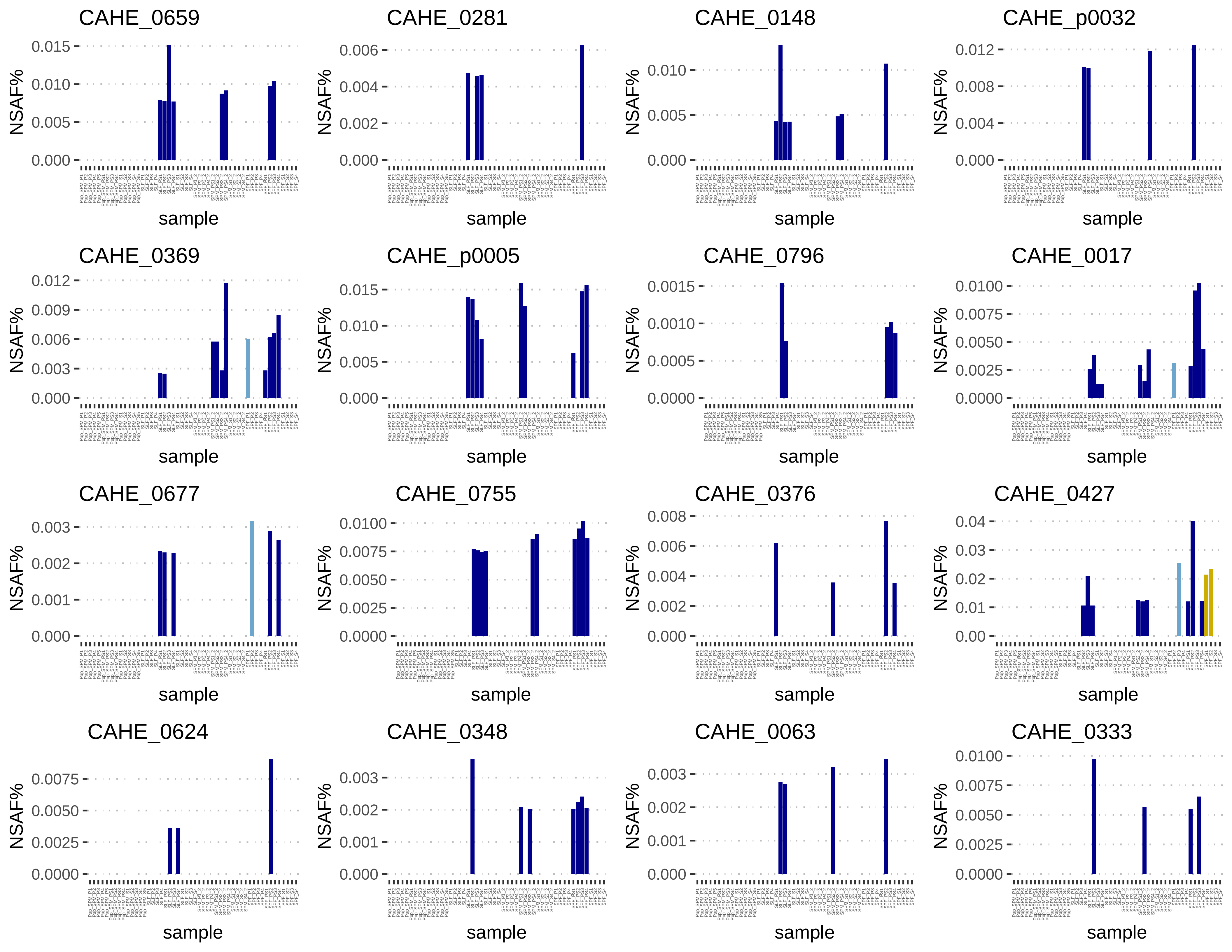
CAHE_0659
CAHE_0281
CAHE_0148
CAHE_p0032
CAHE_0369
CAHE_p0005
CAHE_0796
CAHE_0017
CAHE_0677
CAHE_0755
CAHE_0376
CAHE_0427
CAHE_0624
CAHE_0348
CAHE_0063
CAHE_0333
NSAF%
sample

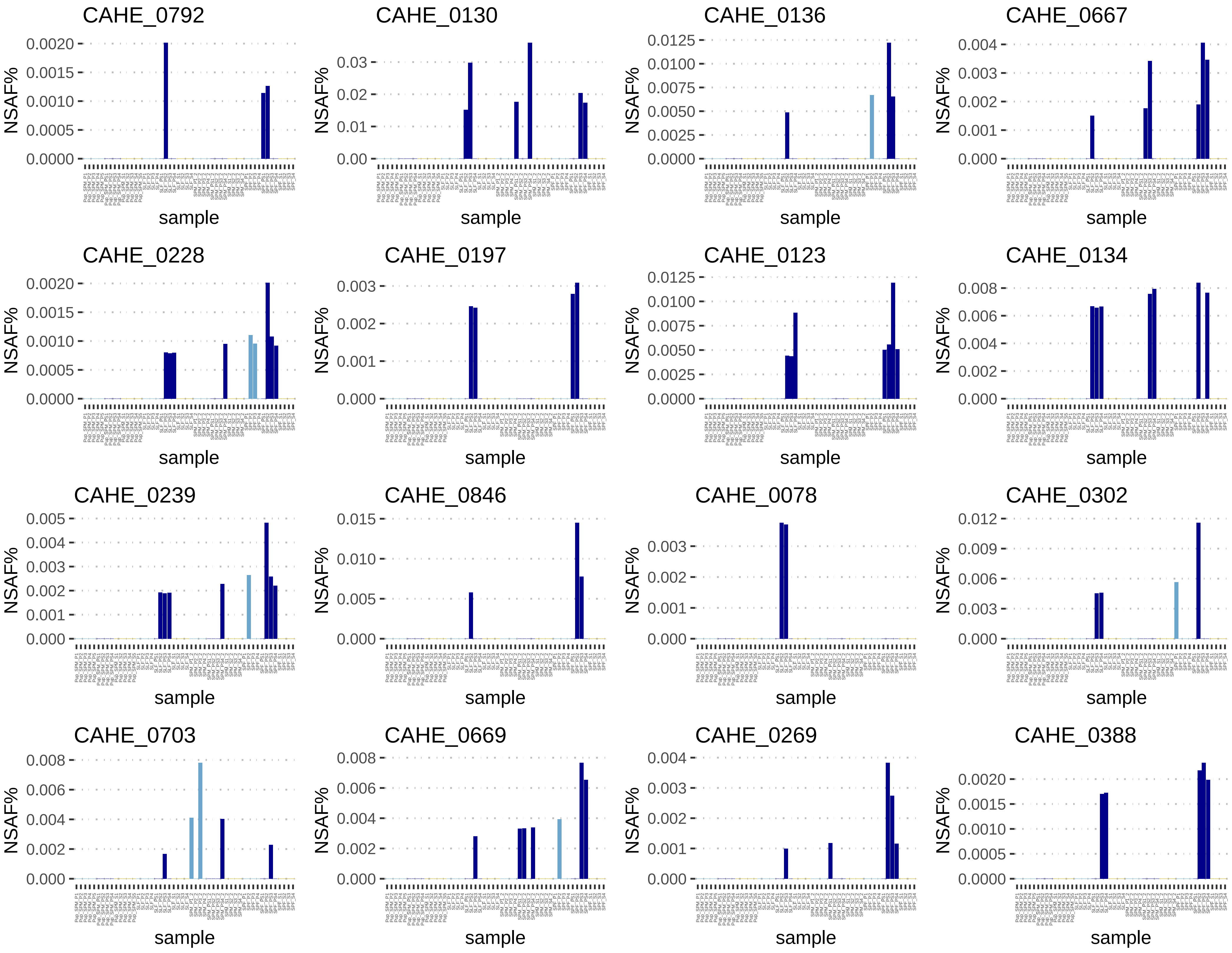
CAHE_0792
CAHE_0130
CAHE_0136
CAHE_0667
CAHE_0228
CAHE_0197
CAHE_0123
CAHE_0134
CAHE_0239
CAHE_0846
CAHE_0078
CAHE_0302
CAHE_0703
CAHE_0669
CAHE_0269
CAHE_0388
NSAF%
sample

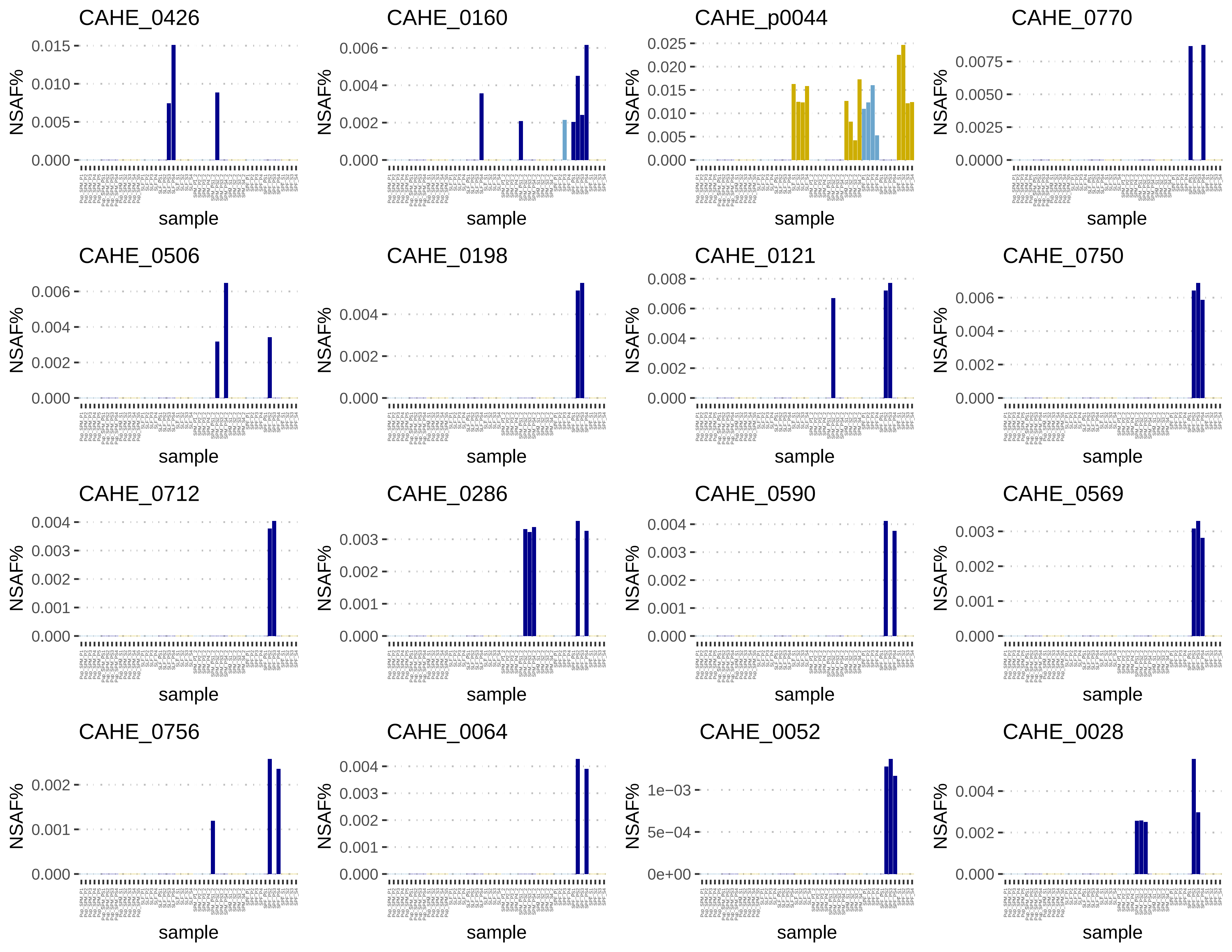
CAHE_0426
CAHE_0160
CAHE_p0044
CAHE_0770
CAHE_0506
CAHE_0198
CAHE_0121
CAHE_0750
CAHE_0712
CAHE_0286
CAHE_0590
CAHE_0569
CAHE_0756
CAHE_0064
CAHE_0052
CAHE_0028
NSAF%
sample
0.000
0.005
0.010
0.015
0.002
0.004
0.006
0.020
0.025
0.0000
0.0025
0.0050
0.0075
0.008
0.001
0.003
0e+00
5e−04
1e−03

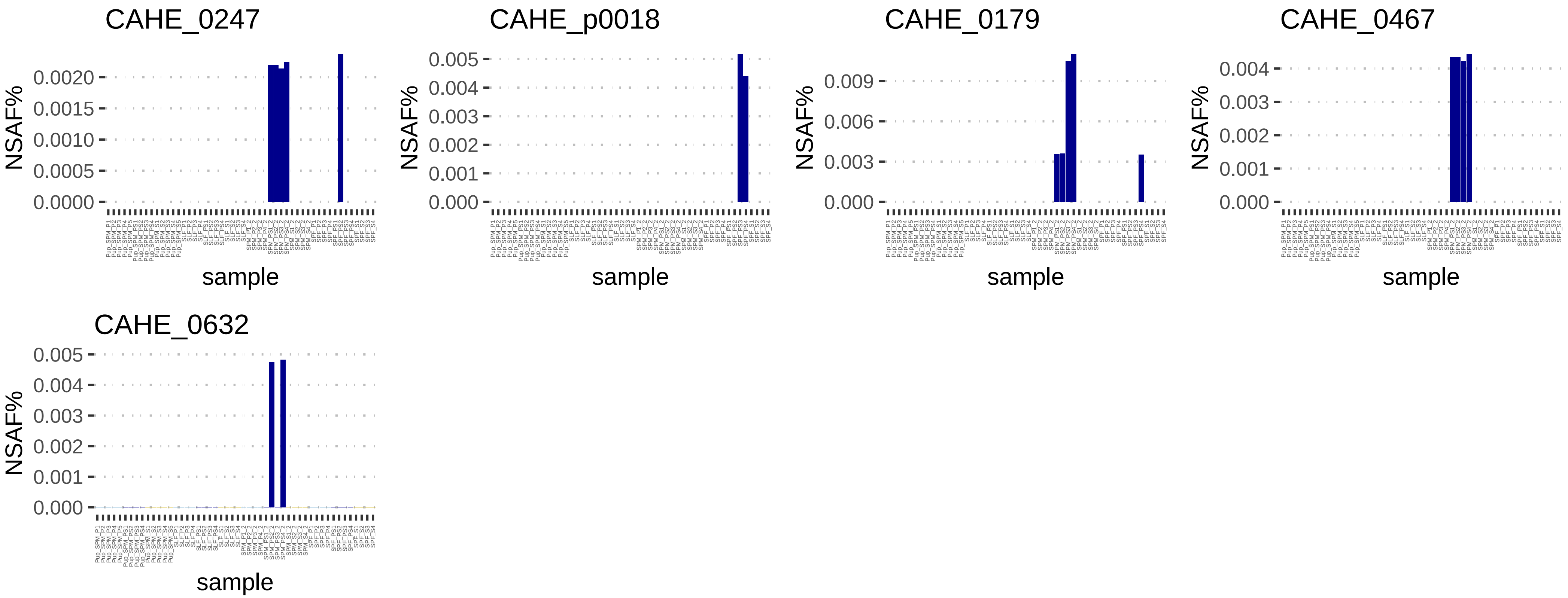
CAHE_0247
NSAF%
0.0020
0.0015
0.0010
0.0005
0.0000
sample
CAHE_p0018
NSAF%
0.005
0.004
0.003
0.002
0.001
0.000
sample
CAHE_0179
NSAF%
0.009
0.006
0.003
0.000
sample
CAHE_0467
NSAF%
0.004
0.003
0.002
0.001
0.000
sample
CAHE_0632
NSAF%
0.005
0.004
0.003
0.002
0.001
0.000
sample
